# DynaMorph: self-supervised learning of morphodynamic states of live cells

**DOI:** 10.1101/2020.07.20.213074

**Authors:** Zhenqin Wu, Bryant B. Chhun, Galina Popova, Syuan-Ming Guo, Chang N. Kim, Li-Hao Yeh, Tomasz Nowakowski, James Zou, Shalin B. Mehta

**Affiliations:** Department of Chemistry, Stanford University; Chan Zuckerberg Biohub; Department of Anatomy, University of California, San Francisco; Department of Biomedical Data Science, Stanford University

**Keywords:** Label-free imaging, self-supervised learning, human microglia, morphodynamic states

## Abstract

The cell’s shape and motion represent fundamental aspects of the cell identity, and can be highly predictive of the function and pathology. However, automated analysis of the morphodynamic states remains challenging for most cell types, especially primary human cells where genetic labeling may not be feasible. To enable automated and quantitative analysis of morphodynamic states, we developed DynaMorph – a computational framework that combines quantitative live cell imaging with self-supervised learning. To demonstrate the fidelity and robustness of this approach, we used DynaMorph to annotate morphodynamic states observed with label-free measurements of density and anisotropy of live microglia isolated from human brain tissue. These cells show complex behavior and have varied responses to disease-relevant stimuli. DynaMorph generates quantitative morphodynamic representations that can be used to evaluate the effects of disease-relevant perturbations. Using DynaMorph, we identify distinct morphodynamic states of microglia polarization and detect rare transition events between states. The methodologies presented here can facilitate automated discovery of functional states of diverse cellular systems.

## Introduction

Organs and tissues of the human body consist of an astonishing diversity of cell types, classically described by their anatomical location, shape, dynamic behavior, gene expression, and protein expression. Morphometry of human cells is widely used to analyze healthy and disease states of cells in clinical pathology and to discover fundamental biological mechanisms. However, automated analysis of morphodynamic states of human cells still remains an unsolved problem, because morphodynamic states can be difficult to identify either visually or with molecular markers. While recent advances in single cell genomics have significantly advanced our understanding of molecular diversity of types of human cells, morphological and behavioral properties of cells cannot be predicted from gene expression data alone. As a result, quantitative analysis of morphodynamic states of human cells with high-throughput methods is a timely area of research.

Recent studies of morphological states of cells have relied on images of fixed cells labeled with a panel of fluorescent markers (1), live three-dimensional imaging of the membrane labeled with genetic markers (2), and phase contrast imaging of live cells (3–6). The morphological states have been analyzed with low dimensional representations computed with geometric or biophysical models (3, 7–11), supervised learning of morphological labels (4, 12–17), and, recently, self-supervised learning of latent representations of shape (5, 6). These analytical approaches have been motivated by the unmet need for quantitative descriptions of specific, complex biological functions, such as motility of single cells (2, 3, 7, 8, 18), collective cell migration (9, 11), cell cycle (4, 12, 13), spatial gene expression (17), and spatial protein expression (14, 16, 19). In addition, data-driven integration of the shape and gene expression (13, 17, 20–23) is now enabling rapid analysis of functional roles of genes. Collectively, these studies illustrate that deep learning is poised to enable quantitative and interpretable analysis of complex cell phenotypes that are too high-dimensional and too subtle for human vision to discern.

Despite this progress, we currently lack automated analysis methods for high-throughput screening of morphodynamic states of human cells, including their temporal dynamics, due to few key technological limitations:

- Measurements of morphodynamics of live human cells can be challenging, because they are difficult to label consistently and without perturbing their native function. While dozens of molecular reporters have been developed to visualize cell structure based on gene expression studies, introducing molecular reporters in primary human cells is frequently impractical.
- Supervised learning from large-scale multidimensional imaging datasets remains challenging, because identifying subtle changes in cell behaviors requires large amount of annotation by biological experts, which is time consuming and hence not scalable.
- While the description of the molecular states of cells in terms of their gene expression profiles is now well established, similar morphodynamic states of cell behavior remain to be established.

Learning the latent representation of cell shapes via self-supervised reconstruction (5, 6, 19) of images is emerging as a promising method of transforming complex phenotypes into quantitative and interpretable shape modes. These shape modes provide a reduced set of cell shapes from which a large variety of cell shapes observed in the data can be generated. The shape modes that can be learned from the imaging data fundamentally depend on the information captured by the imaging modality. Self-supervision of cell contours with a variational autoencoder (VAE) (3) has enabled analysis of shape modes of motile cells. Self-supervision of qualitative phase contrast images with an adverserial autoencoder (AAE) has enabled prediction of the metastatic potential of melanoma cells (5). Self-supervision of fluorescence images of labeled proteins with a regularized vector-quantized variational autoencoder (VQ-VAE) has enabled automated clustering of localization patterns of proteins (19). These papers suggest that self-supervised reconstruction of time-lapse images may enable automated analysis of morphodynamic states of complex cells. In this paper, we demonstrate analysis of the morphodynamic states of human microglia with self-supervised reconstruction of quantitative label-free time-lapse images (16) using VQ-VAE regularized by temporal continuity of the cell shape.

Microglia are the resident macrophages of the central nervous system that are involved in brain development and homeostasis, as well as immune responses (24). Microglia’s response to secreted cytokines or viral infections elicits profound changes in the transcriptome and dynamics of cells that are unique to perturbations (25, 26). Microglia survey brain parenchyma with their highly motile processes and respond to changes in the brain homeostasis by altering cell shape and motility (27, 28), but large-scale features of their dynamic behavior are not well characterized partly due to the lack of tractable molecular labeling tools. For other types of motile cells, either learned features or the manually curated features (3, 5, 6, 18) have been used to describe complex cell dynamics, including random search, persistent migration towards target, changes in cell shape due to interaction with other cells and targets, and endocytosis, among others. Therefore, analyzing the dynamic behavior of microglia under different perturbation conditions represents an ideal use-case for methods of interpreting complex cell dynamics.

We acquired reproducible measurements of cellular architecture and dynamics of microglia under immunogenic perturbations (Figure 1A). To avoid the necessity to virally transduce the cells with reporters, as has been done traditionally(29, 30), we used quantitative label-free imaging with phase and polarization (QLIPP) (16). QLIPP measures phase and retardance, which report physical properties of the density and molecular anisotropy, respectively, of the cells (Figure 1B). Trajectories of live cells (Figure 1C) were used as an input to for DynaMorph, a deep-learning enabled framework that automatically extracts descriptors of shape and dynamics. DynaMorph relies on vector-quantized variational autoencoder to learn a latent representation of cell shape (Figure 1D). Enforcing temporal continuity of the latent shape modes in the neighboring frames leads to a quantitative representation of cell shape. We found that latent shape space regularized by the temporal continuity generalized well to new experiments. We evaluated dependence of the latent shape space on the imaging modality, and found that shape modes learned from quantitative measurements were consistent across experiments. In this study, we combine shape and motion descriptors of cells to represent cell behavior and employed mixture modeling to identify morphodynamic states (Figure 1E). From the trajectory of cells in the latent shape space, we identified transitions among morphodynamic states of single cells. The same approach enabled detection of transitions in the morphodynamic states of cells as a result of immunogeneic perturbations.

**Fig. 1.**
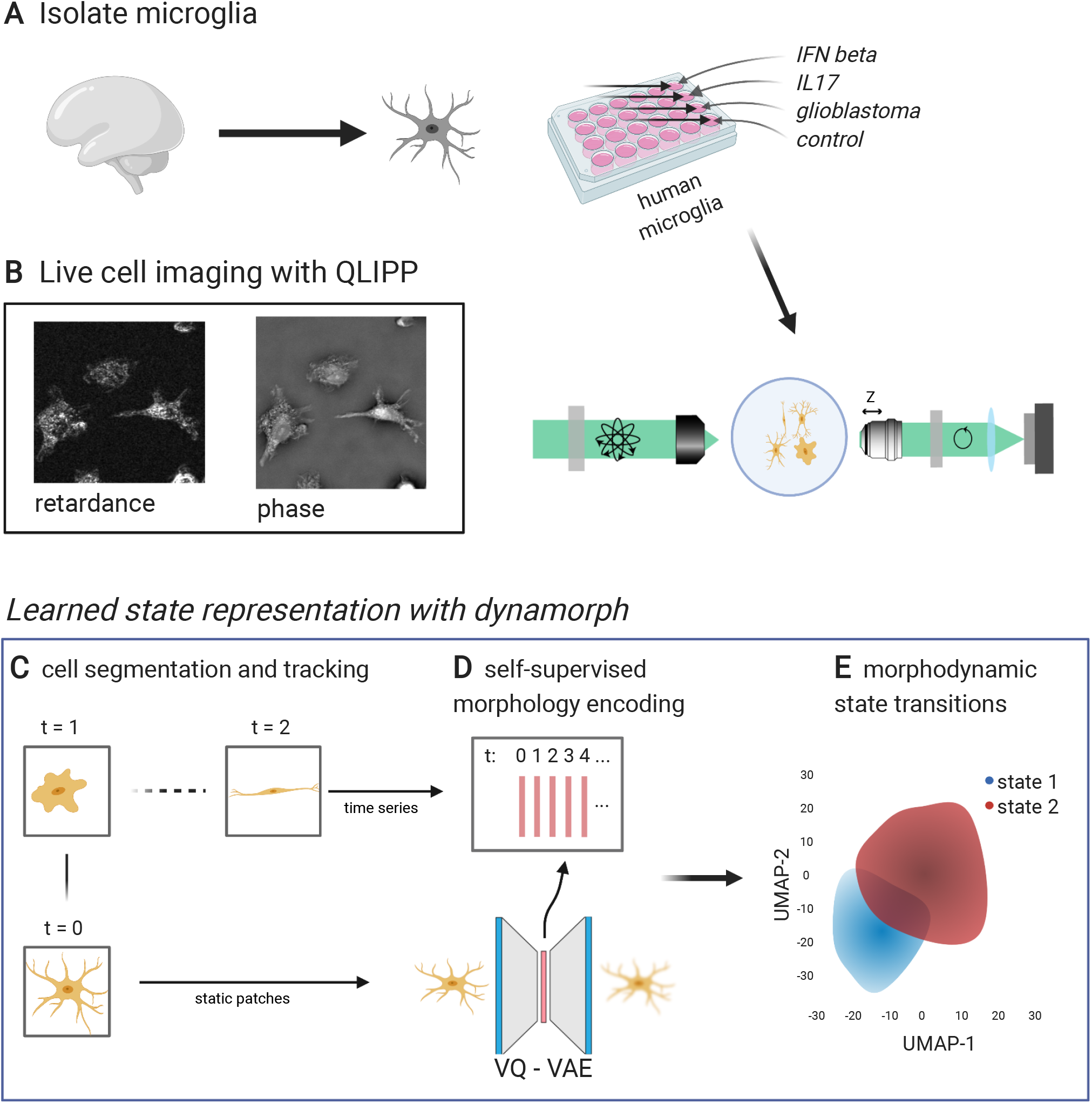
DynaMorph enables automated discovery of morphodynamic transitions in human microglia: **(A)** Human microglia are isolated from brain tissue and plated in 24-well plates and perturbed with cytokines of relevance to infection (IFN beta, IL17) or cancer (glioblastoma supernatant), (B) Morphodynamics of perturbed microglia, along with control cells, are imaged using quantitative label-free imaging with phase and polarization imaging (QLIPP), which measures isotropic and anisotropic optical path lengths of cells. **(C)** Cells were segmented and tracked by in-house developed tools. (**D**) Generalizable and quantitative shape modes were learned from the thousands of tracked cells using a self-supervised model that reconstructs cell shape. **(E)** Morphodynamic states and transition among states under each perturbation were revealed via dimensionality reduction algorithms and clustering of most significant features.

## Results

### A: Learning a latent representation of shape that generalizes across experiments

We evaluated fluorescent probes that label membrane and QLIPP for imaging shape and dynamics of primary microglia. We observed that the distribution of membrane probes can vary over time due to the turnover of the membrane. On the other hand, phase and retardance images acquired with QLIPP provided consistent (Fig. 1 - supplement 1) readout of the cell shape and dynamics. We acquired label-free time-lapse images from primary human microglia over two sets of experiments. First set of experiments was done with purified microglia without any perturbation and provided the training data for the model. The second set of experiments was done in the presence of immunogeneic perturbations, and provided the data used for testing the model and discovery of cell states. In the perturbation experiment (test set), multiple treatments relevant to infection or cancer were applied to mimic different inflammatory states in neural tissue. The time-lapse data was processed by segmentation and tracking modules developed in-house to extract patches/movies of individual cells (see Methods, Fig. 1 - supplement 2, and Fig. 1 - supplement 3).

We observed significant diversity in shape and behavior of microglia across perturbations (see videos in Video Set 1). While microglia are classically described as amoeboid, ramified, mobile, or immobile, depending on their cell state, they exhibited a variety of complex shapes and state transitions in our videos that we attempt to capture in our model. We used self-supervised deep learning to learn quantitative description of complex shapes of microglia.

Patches of individual microglia from the training set were used to train a variant of autoencoder: Vector Quantized Variational Autoencoder (VQ-VAE) (31)(Figure 2, Fig. 2 - supplement 1). The VQ-VAE was trained to reconstruct phase and retardance channels of the patches. The reconstruction of images serves as an auxiliary task to compute a latent vectors for all patches. The latent vectors of all patches constitute the latent (or learned) shape space of microglia, which we abbreviate to *shape space* in the following text.

**Fig. 2.**
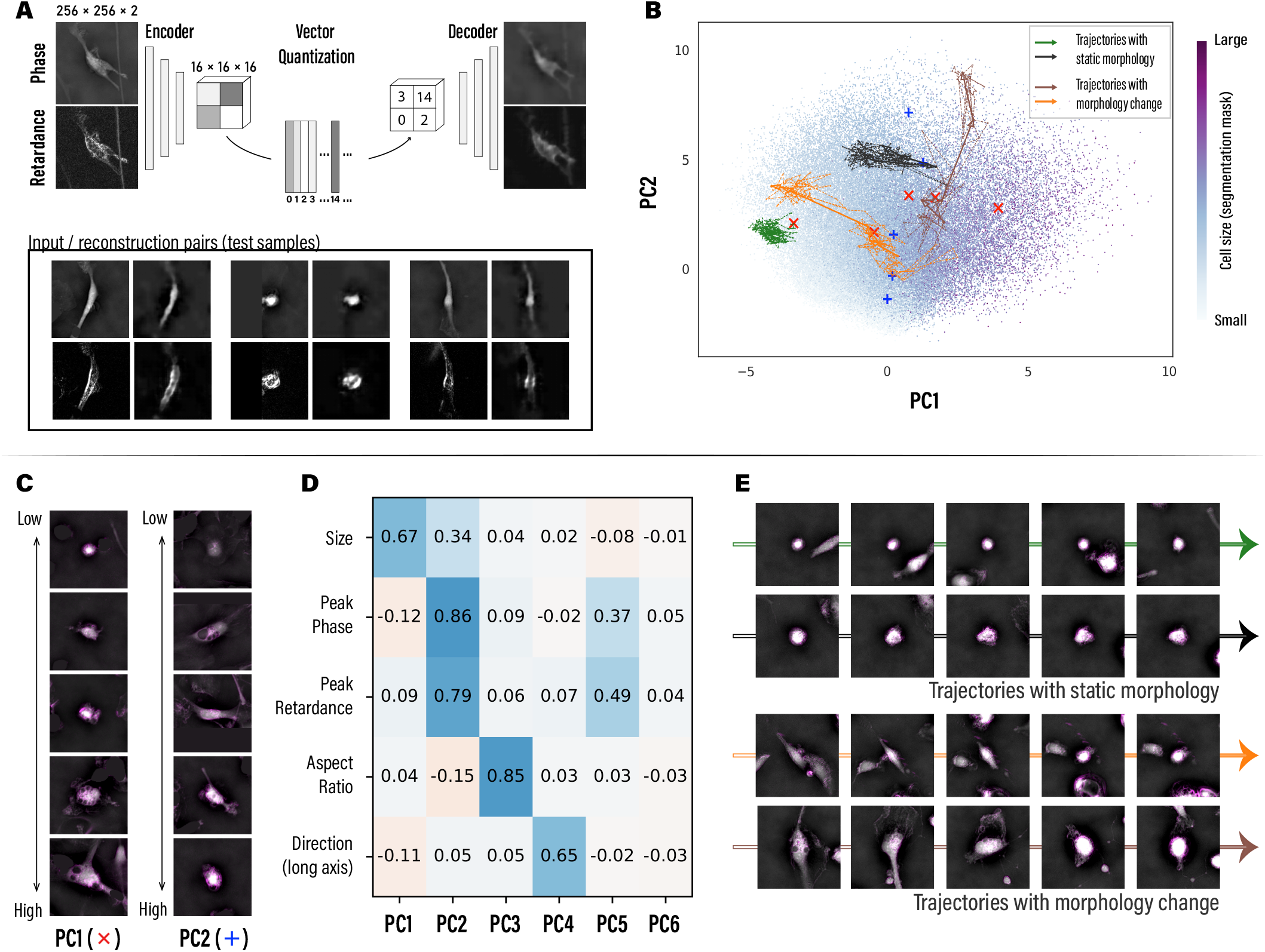
DynaMorph learns a generalizable and quantitative representation of cell shape via self-supervision: **(A)** We encoded the shape of microglia in quantized latent vectors and reconstructed the shape from the latent vectors through a vector quantized variational autoencoder (VQ-VAE), which is trained in a self-supervised fashion. The phase and retardance images from the test experiment (test set) shown here were encoded and reconstructed using models trained on the data from control experiment (training set). Comparison of reconstructions from training set Fig. 2 - supplement 1 demonstrates generalizability of the model. **(B)** Shape modes that describe the most significant differences were computed with principal component analysis (PCA) of the latent vectors. Resulting top 2 principal components (PC) are visualized, in which each dot represents a single cell patch from the test set and its color indicates the size of cell segmentation mask. 4 representative trajectories: two cells with static shape (green and black) and two cells undergoing changes in shape (orange and brown) are plotted in the PCA space as arrows. **(C)** The first two shape modes (PC1 and PC2) were interpreted by sampling representative patches along each PC axis (cross markers in panel B). Clear trends of changing size/cell density could be observed. Note that patches shown combine both phase channel (as grey scale) and retardance channel (as magenta shades). **(D)** We explored correlation between the top 6 PCs and selected geometric properties of the test set using Spearman’s rank correlation coefficients. PC1 and PC2 both show positive correlations with cell size; PC2 is also highly correlated with cell’s peak phase and retardance that measure isotropic and anisotropic density, respectively; PC3 is correlated with cell’s aspect ratio; PC4 is correlated with the orientation of the cell (long axis of the cell body). Note that all results are consistent between training set cells and test set cells (Fig. 2 - supplement 1C). **(E)** Representative cells plotted in **(B)** are visualized. Significant changes in shape could be linked to jumps in the latent space.

Multiple variants of autoencoder were tested, and the VQ-VAE appeared as the best-performing model architecture with the lowest reconstruction loss and consistent shape modes (see Supplementary Notes, Fig. 2 - supplement 2, and Fig. 2 - supplement 3), in accordance with results reported for the general image reconstruction task (31, 32). The self-supervised encoding led to a noise-robust and interpretable learned shape space of cells as discussed in sections below.

In addition to the regular reconstruction and regularization loss, we imposed a matching loss in the model to minimize frame-to-frame distance between latent vectors of the same cell along its trajectory, based upon the prior that cell architecture changes smoothly between consecutive frames. This approach is motivated by the previous work that enforced temporal slowness constraint on the feature representation of adjacent frames (33). Similar concepts have been applied to learning similarities among human face images (34). As a result of applying the temporal matching loss, cell trajectories with small changes in cell morphology tend to stay localized in shape space, while trajectories with significant size/density/shape changes can be easily identified as trajectories showing large jumps in shape space. For full details of the training procedure, see the Methods section and loss curves during training Fig. 2 - supplement 4A.

To evaluate how the shape space depends on the input data, we trained and evaluated models using individual label-free channels (brightfield, phase, and retardance). Models trained on single channels show worse performance than the joint model (see Supplementary Notes and Fig. 2 - supplement 4).

Overall, we find that the shape space of the DynaMorph autoencoder substantially compresses the data in the raw imaging– by over 680 times. The model trained on normalized cell patches has reconstruction loss 0.19 *±* 0.21SD (cell/foreground are usually 3-4 SD brighter than background). This was further validated on the test set, on which model reconstruction loss is 0.25 *±* 0.20SD with no detectable signs of overfitting.

Comparison of reconstructed shapes from the test set (Figure 2A and Fig. 2 - supplement 5) and training set (Fig. 2 - supplement 1A) along with the analysis of the shape space described in the next section show that our self-supervised model trained on control experiment generalized well to unseen cells that were treated with multiple perturbations.

### B: Learned Shape Space provides a quantitative description of shape

The learned shape space could provide a quantitative reduction of complex shapes of cells. We sought to evaluate if the shape space provides quantitative metric of shape, i.e., we sought to answer the following questions: Are the most significant modes of the shape space interpretable? Does the distance in the shape space capture shape similarity?

#### Interpretation of the learned shape space

We use principal component analysis (PCA) to extract the top shape modes and visualize the shape space. Distributions of the first two principal components (PCs) are displayed in Figure 2B, in which each individual dot represents an encoded cell patch and the color indicates the cell size measured from its segmentation mask. A horizontal gradient of cell size is observed, which suggests quantitative relationships between top PCs and cell properties. Fig. 2 - supplement 1B shows similar results for the cells from the training set.

To further unravel these relationships, we sought to link the top PCs with heuristic cell geometric properties. We first sampled cell patches along PC axes (Figure 2C and Fig. 2 - supplement 6) and interpreted the cell properties visually. We were able to identify significant correlations in each PC: PC1 relates with cell size; PC2 relates with peak phase and retardance, which measure cell density; PC3 and PC4 relate with cell orientations. Quantification using Spearman’s rank correlation coefficients (Figure 2D) is consistent with our visual interpretation of each PC. Thus, the first two principal components of the shape space correlate strongly, but not exclusively, with cell size and cell density.

The correlation of the first PC with the cell size intriguingly parallels the analysis of cell masks of other immune cell types in Chan *et al*. (6). They also show that the most significant shape mode of three different motile cell types is cell size. In addition, the second most significant shape mode in our data relates with cell density as measured with phase and retardance, which comports with the result in Zaritsky *et al*. (5) that light scattering is a significant descriptor of cell behavior. Thus, our results show that the shape space learned by the VQ-VAE model indeed capture dominant shape modes that have been reported in distinct cell types. This correspondence suggests that the cell size and cell density may be highly conserved descriptors of cell states, due to mechanisms that are yet to be understood.

Intriguingly, while 80% of the shape variance of ameboid cells analyzed in Chan *et al*. (6) is captured by 5 shape modes, we found that the first 4 shape modes (or PCs) of microglia account for less than 20% of all variance (Fig. 2 - supplement 7A). The remaining variance can be due to more complex aspects of microglia shape, such as diversity of protrusions, sub-cellular structures and variations in cell density, location of nuclei in migrating cells, etc. UMAP(35) projections of the shape vectors of cells show no apparent clustering patterns (Fig. 2 - supplement 7B and C), suggesting that microglia shape undergoes continuous change and hence no discrete states could be defined based on static frames.

To validate that the learned shape space is uniquely informative, we applied PCA to the input images and followed the same analysis procedure. The derived top PCs have weaker and less clear correlations (Fig. 2 - supplement 8A) and consequently are harder to interpret. The principal components of input images could not be separated by clustering analysis (Fig. 2 - supplement 8B and Supplementary Table 1). Thus, shape space learned with VQ-VAE indeed provides an interpretable and quantitative representation of the complex shape of microglia.

#### Trajectories in the Learned Shape Space characterize dynamics

To better understand how the shape space captures dynamics, we generated single-cell trajectories of latent representations by concatenating representations of static frames. As a control, we compared the distance between the latent representations of temporally adjacent and non-adjacent frames (both before and after quantization). We found that the distance between temporally adjacent frames is substantially less than non-adjacent frames and random baseline, indicating that our shape space captures the continuity of cellular dynamics (Fig. 2 - supplement 9).

We further compared the distance distribution between latent representations generated with and without matching loss. Results show that the reconstruction task already reduces the distances between adjacent frames, as compared to random pairs, while including the matching loss further regularized the shape space such that it reliably represents the temporal evolution of the shape (Fig. 2 - supplement 9).

Visualization of individual cell trajectories in the shape space show that majority of cells had stable shape modes throughout the imaging period. In Figure 2E we illustrate some representative trajectories (marked in green and black) that are in static cell states and have stable appearances. Their top shape modes are relatively confined, suggesting that the cell states, as defined by their visual features, are stable throughout time. However, we also observed a number of “leap” events in the shape space, which coincide with cells undergoing significant transitions in shape. Representative examples (marked in orange and brown) are also shown in Figure 2E, in which transitions in size or density could be observed. Full video clips of these example cell trajectories could be found in Video Set 2.

### C: DynaMorph discerns between pro- and anti-inflammatory microglia states

Microglia can respond to pro- and antiinflammatory stimuli by changing their shape, dynamics, and gene expression. While a range of small molecules, including cytokines and chemokines and their individual effect on microglia has been well described, cell conditions with both pro- and anti-inflammatory signaling molecules in the microenvironment present a more challenging task. One example is a brain tumor micro-environment, such as glioblastoma (GBM), where a wide array of signaling molecules can simultaneously promote pro-inflammatory states of microglia, and anti-inflammatory states. Characterizing the nature, as well as heterogeneity of microglia states under defined perturbation conditions could enable a systematic, data-driven method for defining the complex responses of microglia to disease states, and could ultimately serve as a benchmark for evaluating the effects of pharmacological perturbations in a high-throughput manner.

We treated cells with a pro-inflammatory cytokine IL17, which has been implicated in several brain diseases(36–38), and IFN beta, an anti-viral and anti-inflammatory cytokine (39, 40). In parallel, we treated microglia with supernatant from primary human GBM cells. Cell patches from the test conditions were processed and encoded through the DynaMorph pipeline. Video clips of some representative cell trajectories from each condition can be found in Video Set 3. As already shown in fig. 2, reconstruction of patches and interpretation of shape space are reliable across perturbations.

We first explored distribution of shape modes and speed of cells under different perturbations. For each cell, we collected and averaged its PC1 and PC2 values along the trajectory (Figure 3A). Distributions of these trajectory-averaged PC1 and PC2 values are shown for each treatment in Figure 3B and Fig. 3 - supplement 1. PC1 of shape showed a similar distribution across all four conditions, suggesting that the size distribution of the cells is consistent across different conditions. In the PC2 space, which correlates closely with cell density, IL17 and IFN beta subsets had similar distributions, which differed from control and GBM subsets. This suggests that microglia under IL-17 and IFN-beta treatment tend to have lower density while maintaining a similar size. Speed is plotted separately (Figure 3B) and against 2 top shape modes as kernel density estimation (KDE) plots (Figure 3C and D). Microglia display a broad range of motility under different conditions, with the fast-moving population (control subset) traveling over 2 times faster than the slow-moving population under an anti-inflammatory IFN beta treatment. Note that this measurement is based on centroids of cells and hence could be confounded by shape changes, though the scale is much smaller than typical cell movement.

**Fig. 3.**
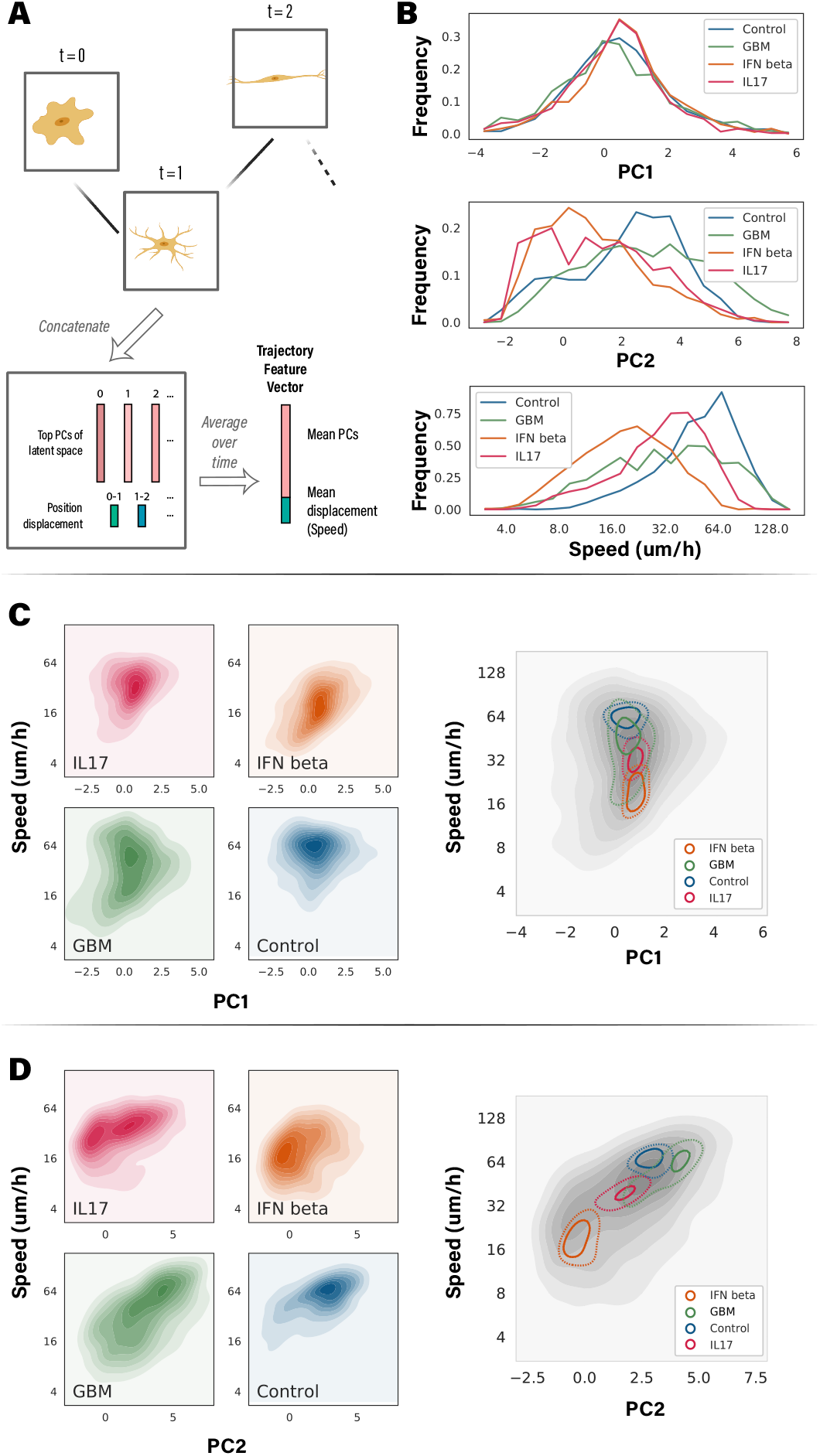
DynaMorph reveals differences in morphodynamics of microglia under different perturbations. **(A)** To compare how the morphodynamics of microglia change in response to different perturbations, we computed trajectory-averaged features as follows: principal components of the shape space of the frames of a tracked cell were concatenated with position displacements between frames and averaged over each trajectory. **(B)** We analyzed the distributions of Trajectory Feature Vectors under multiple perturbations by plotting probability densities of trajectory-averaged PC1, PC2 and mean displacement, i.e., mean speed. The geometric properties of cell size and density correlated highly with PC1 and PC2, respectively. Trajectories under IFN beta treatment show significant differences from trajectories under control treatment in cell density and speed. **(C)** and **(D)** We analyzed changes in average cell size (PC1) and average cell speed, as well as average cell density (PC2) and speed by plotting their joint distributions. Left panels show density plots of distributions under different perturbations. Right panels show the combined distributions, with apex contours under each perturbations marked. Separations between IFN and control conditions observed in **(B)** were confirmed.

To better understand the interplay between shape and dynamic behavior, we constructed Trajectory Feature Vectors (TFVs) by first concatenating the Principal Components of the shape space with the instantaneous speed of the cell (between two consecutive frames), and then averaging them along the trajectory(Figure 3A). These vectors are then be clustered to distinguish morphodynamic states of the cell population (Figure 1F) and analyze how cells transition between these states.

Interestingly, while microglia treated with a single cytokine (IL17 or IFN beta) displayed unimodal speed distribution (Figure 3B), GBM supernatant induced a much wider range of speeds, with clearly distinguishable individual peaks around 16, 32 and 60 um/h. This finding suggests that several intermixed populations of microglia may exist under the GBM cell microenvironment, further highlighting the importance of single-cell tracking in understanding cell behavior under complex cell signaling milieu. Given the striking differences in the cell speed, we plotted changes in top shape modes and average cell speed (Figure 3C, D). Our data reveal that microglia under pro-inflammatory stimuli had a greater proportion of faster and denser cells, while under the IFN beta treatment the cell distribution was shifted towards slower moving cells. Control microglia had a distribution resembling the trend observed in IL17 treatment, suggesting that *in vitro* culture conditions induce microglia activation, as have been previously noted(41). In contrast, cells treated with the GBM supernatant had a shape distribution that simultaneously resembled both control, IL17 and IFN treatment, suggesting that different cell populations within this condition would adopt either pro- and anti-inflammatory morphodynamic cell profile, rather than being a homogeneous population of an intermediate response.

The separation was further revealed in a discriminative study that predicts treatment/condition purely based on TFVs (Fig. 3 - supplement 2, see Supplementary Notes for details). In the results from gradient boosted decision tree models (42), confusion between control subset and IFN beta subset is especially small, while confusion between IL-17 and GBM is larger.

Together, this analysis suggests that at least two broad morphodynamic states of microglia can be detected using DynaMorph. In particular, exposure of microglia to IFN beta induces a state characterized by reduced density and migration speed. In contrast, exposure to GBM supernatant, which likely contains pro-inflammatory and anti-inflammatory cytokines, induces a broader distribution that might consists of multiple morphodynamic state.

### D: Identification of morphodynamic states and transitions between them

To further probe how learned shape space and motion of microglia under different perturbations can reveal morphodynamic states, we employed mixture modeling of the trajectory feature vectors (TFVs) of test cell patches to locate and quantify the underlying cell states. We found that trajectory feature vectors formed a continuous spectrum rather than discrete clusters. Therefore, instead of using clustering algorithms such as hierarchical or K-means clustering, we employed a Gaussian Mixture Model (GMM) to identify morphodynamic states in the test population, which could be further used to explain distributional differences between perturbations. The model revealed two components in the combined test population (See Methods for details). We refer them as state-1 and state-2 respectively in the following text.

Comparing with inter-condition differences, inter-state separation is much stronger (Fig. 4 - supplement 1) which suggests the states identified by GMM are superior representations of morphodynamic modes of microglia. Representative samples from the two states are illustrated in Figure 4B, with their trajectories plotted as colored lines. Representative cell trajectories could also be found in Video Set 4. Morphodynamic distributions of the two states are described in Figure 4F, details could be found in Supplementary Table 1. Qualitatively, based on the cell appearance and features of the two components, we describe state-1 (blue) cells as large-sized, low density and slow-moving population, and state-2 (red) cells as a higher density and fast-moving population. The two states differ the most on PC2 and mean distance, and have slight difference on PC1 as well.

**Fig. 4.**
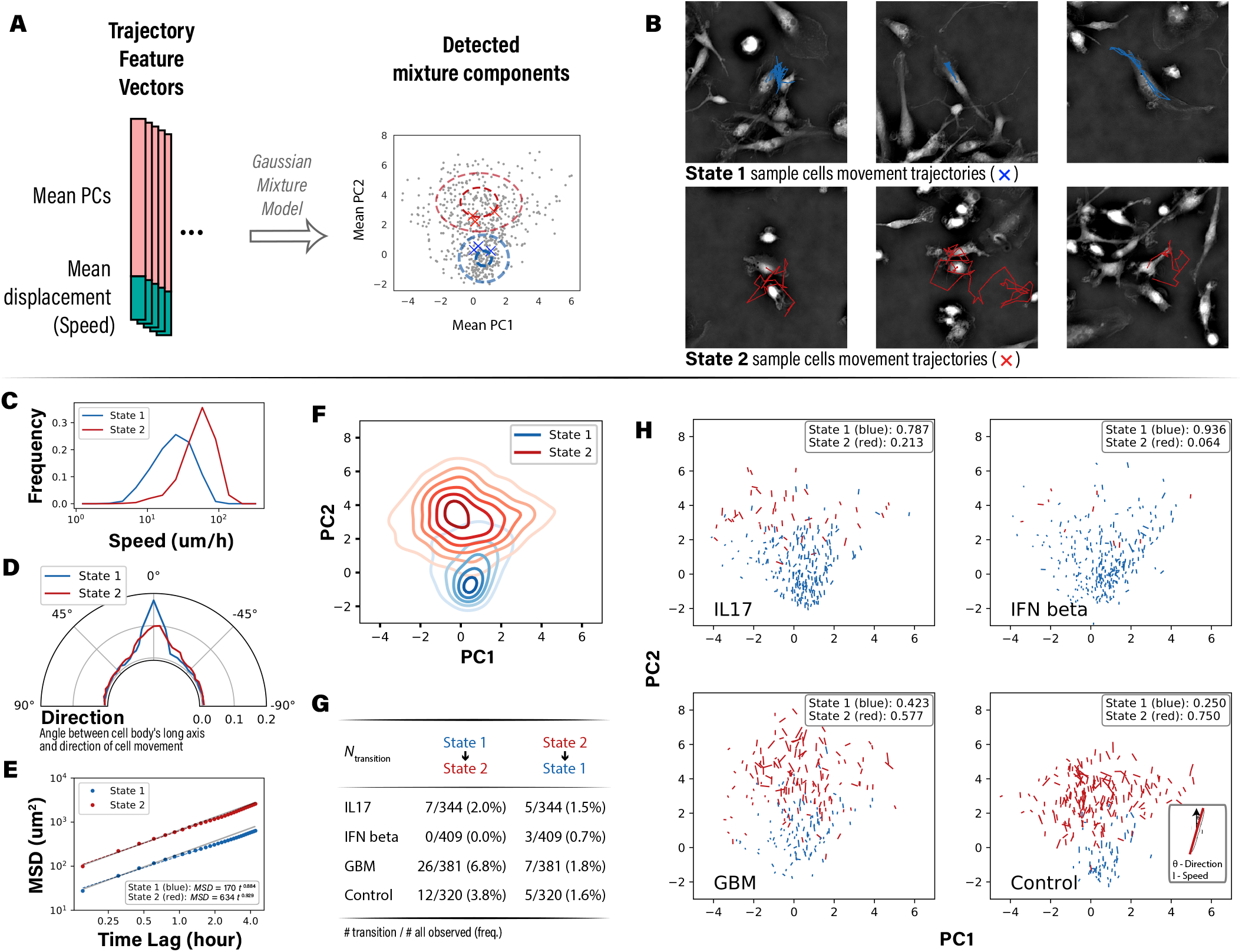
Cell trajectories are clustered into two states based on the morphodynamic features. **(A)** We identified two states of the trajectories by pooling their TFVs and clustering them with a Gaussian Mixture Model (GMM). **B** Representative trajectories from the two states (corresponding to cross markers in panel A) are shown: initial frames are visualized, and the movement of cells in the following frames are marked. **(C–F)** We compared two morphodynamic states with multiple metrics: **(C)** cells in state-1 were found to be slower than cells in state-2 from the probability densities of trajectory-averaged speeds. **(D)** cells in state-1 were found to align more with directions of migration than cells in state-2 from the probability densities of angles between cell body (long axis) and the directions of movement. **(E)** cells in state-1 and state-2 were found to undergo random motion (grey shades indicate standard random motion) over the time-scale of hours from the mean squared displacement (MSD) curves of the trajectories. Consistent with the speed distribution, state-2 cells travel longer distances on all time scales. **(F)** cells in state-1 had narrower distribution of size and density relative to cells in state-2, as seen from the Kernel density estimates (KDE) of PC1 (cell size) and PC2 (cell density). **(G)** We counted the number of trajectories that transitioned between two states under each perturbation and found rare instances of transition as noted in the table. Note that only trajectories longer than 4.5 hours are considered. **(H)** Metrics shown in **(C)–(F)** were elaborated per perturbation with scatter plots of the metrics: each marker represents a single trajectory, whose length indicates the speed of the cell. Direction of the marker is aligned with the angle between cell body and movement, with a vertical line indicating a perfect alignment. (See legend in the bottom right panel). Fraction of trajectories that occupy either state under each perturbation are noted in the insets.

A more quantitative view on cell motion demonstrates that state-2 cells, enriched in control subset, are on average 2.3 times faster than state-1 cells (Figure 4C) that appear mostly in IFN beta subset, in line with the subset-level speed differences we noted above. We also consider directionality in the two states, which represents an important feature of cell migration, noting how the movement trajectories of state-1 cell in Figure 4B aligned with the direction of its cell body. We validated this motion pattern on all microglia by calculating angles between movement and long axis of cell body in each frame and conducting vector sums over these angles. Resulting trajectory-summed directions are visualized in Figure 4H. Distributions of the directions shows dominating peaks at 0°(Figure 4D), indicating a very high preference for cell body-aligned motion. Notably, state-2 cells have less such tendencies, which is in part due to the morphology distinction as state-2 cells usually appear as dense, round shapes that do not have significant long axes, in contrast to state-1 cells that appear more in oval shapes. Analysis on the Mean Square Displacement (MSD) of the two states were then conducted. Figure 4E and Fig. 4 - supplement 2 plot the log-log fit of MSD over time lag. Both states have similar slopes (∼ 0.9) that are close to 1, indicating a near-random motion. State-2 have larger intercept, indicating a larger diffusion constant and hence faster migration. Taken together, the distinct morphodynamic properties exhibited by two cell states revealed by GMM suggests the two-state model is a good description of cell populations across perturbation conditions.

Among the different perturbation conditions in the test population, cells are widely distributed between two states depending on the treatment (Figure 4H and Supplementary Table 1): majority of the control subset cells are in state-2 (∼ 75%), while almost all IFN beta-treated cells are in state-1 (*>* 90%); GBM subset falls between the two extremes with near-equal split on the two states. Based on the previous observations that microglia adopt an amoeboid shape upon exposure to anti-inflammatory stimuli, we speculate that state-2 with faster moving, denser and round-shaped cells represents activated microglia, while state-1 that is characterized by ramified and slower moving cells represents homeostatic microglia. Note that due to the huge separation between control and IFN beta subsets, many of the cells in IL-17 subset are labeled as state-1. We treat this conflicting message as a side effect of the simplified two-state model, as feature distribution in Figure 3C, D clearly shows the distinction between IFN beta and IL-17 subsets in cell shape and motion.

The treatment of microglia with glioblastoma supernatant lead to a distribution that included nearly equal contribution from both states, suggesting a complex mixture of both pro- and anti-inflammatory cytokines. We further compared the detected morphodynamic states with scRNA measurements of the same cell population. Interestingly, the separation of stage-1 and state-2 cells from control and IFN group parallels the clustering pattern of cell transcriptome (see Supplementary Note, Fig. 4 - supplement 3 and Fig. 4 - supplement 4 for details), which hinted the potential of correlations between morphodynamic/behavior change and transcriptomic change in response to perturbation.

Furthermore, the quantitative definition of states allowed us to rigorously characterize state transition events such as the ones shown in Figure 2E (orange and brown trajectories). Given the shape and motion changes throughout the imaging period, we separated each trajectory into segments (2 hour each) and applied the unsupervised GMM to estimate posterior probabilities of cell states for each segment. Cells that have different states assigned to different segments would be of great interest to the study of dynamics and behavior of broad cell types under different conditions. We counted the observed transition cases in the test set and report the numbers in Figure 4G. In our analysis, transition events are very rare among cells treated with IFN beta, while cells treated with GBM supernatant occurred more frequently. While both directions of transitions were observed within the imaging period, cells in state-1 are more likely to transition to state-2 than vice versa within the chosen time frame. Given that cell states should have reached equilibrium after several days in culture at the time of the imaging experiments, these results suggest that the transitions from state 2 to state 1 occur at a different time scale (i.e., much slower). Representative state-transition cells can be found in Video Set 5.

### E: DynaMorph can be easily extended to other cell types

Building on the successful application of DynaMorph to the discovery of microglia cell states, we sought to evaluate the generalizability of our model by applying DynaMorph to non-microglia cells that were present in our cultures. Specifically, neural progenitors can be easily identified by their elongated cell bodies with radially extending processes, different from the amoeboid or ramified shape of microglia.

VQ-VAE trained with the control experiment cells generalizes well to the progenitor cells from the perturbation experiment, with a mean VAE reconstruction loss of 0.25 SD over patches containing progenitor cells, close to the reconstruction performance of microglia. Then through the same trajectory building and encoding pipeline, we extracted the TFVs for 125 identified progenitor cells. Sample cells along with their movement are illustrated in Figure 5A, corresponding cell trajectories can be found in Video Set 6.

**Fig. 5.**
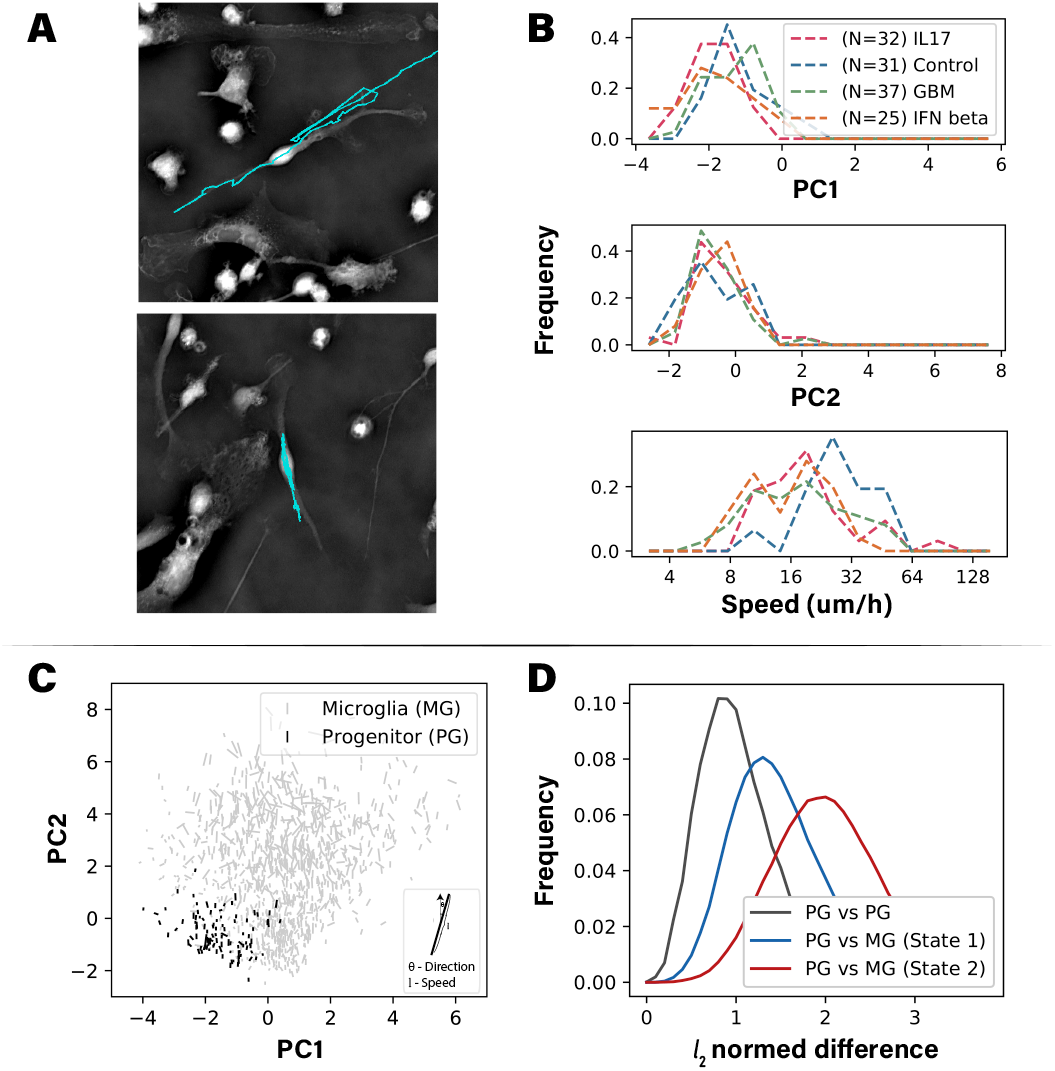
Morphodynamic state of neural progenitor cells in the test dataset. **(A)** Example trajectories of progenitor cells are shown; cell motions are marked as cyan lines. Note that the elongated cell bodies align with cell motions. **(B)** Probability densities of top principal components and cell speeds. Shape of progenitors is robust to perturbations. **(C)** Scatter plots comparing shape, speed, and orientation of progenitor cells and microglia (similar to Figure 4H). The two cell types occupied different regions of the shape space. **(D)** *l*_2_ normed differences of Trajectory Feature Vectors between progenitor cells and microglia in two states. Note that progenitor cells are closer to microglia that are in state-1.

Figure 5B shows the first two shape modes and motion features of progenitor cells. These results indicate that progenitor cells are smaller (lower PC1) and less dense (lower PC2) than microglia. Progenitor cells have a distinct cell motion as compared to microglia. Aligned with its polarized shape, progenitor cell’s movement usually follows the direction of its protrusions, as illustrated in Figure 5A. This results in a much stronger preference for directional motion as compared to microglia. And consistent with the fact that these are not immune cells, progenitor cells did not substantially alter their behavior when subjected to immunogenic perturbations.

In Figure 5C, we plot the average of shape features over trajectories for microglia and progenitor cells, which demonstrates a clear difference in shape between the two cell types. Finally to directly compare morphodynamic states of these two cell types, we used the TFVs which contain motion and shape information to calculate pairwise Euclidean distances. Figure 5D illustrates the distribution of distances between progenitors or between progenitor and microglia. Results reaffirm that microglia are in general different from progenitor cells, and state-1 microglia are more similar to progenitor cells. This result agrees with the visual evidence that state-1 microglia have similar elongated cell shape and higher tendency for directional movement (Figure 4B). Together, these results suggest that our model performs equally well on the highly motile immune cells and less dynamic progenitor population, and can be used to distinguish between different cell populations while maintaining the level of granularity to identify different states within the same cell type.

## Discussion

In this work, we present DynaMorph: a deep learning framework for the dimensionality reduction and for quantitative analysis of live cell imaging data. We demonstrate utility of the framework by analyzing morphodynamic states from two complementary physical properties of cells - their density and anisotropy. By combining the novel self-supervised learning with tracking and mixture modeling, the workflow allows us to analyze cell dynamics and behavior on both individual cell-level and population-level.

Our work formalizes an analytical approach for data-driven discovery of morphodynamic cell states based on the quantitative shape and motion descriptors. All components in the pipeline can be adapted based on the goals of analyses and characteristics of the imaging system. Here, we specifically demonstrate the utility of this method to discover and classify morphodynamic states of human primary microglia cultured *in vitro*, and to transitions between these states upon exposure to polarizing, disease-relevant environments. We show that glioblastoma microenvironment differently affects microglia cells by inducing a wide range of states consistent with both pro- and anti-inflammatory response. While it is not known what factors predispose individual microglia cells to adopt one or another state under complex cell environments such as brain tumors, these specific cells could be used for downstream single cell profiling to discover molecular correlates of those events and consolidate the relationship between morphodynamic behavior and transcriptomic alteration. Once generalizable models are trained and evaluated as we describe, label-free imaging and DynaMorph can enable online inference of changes in the morphodynamic states in cell population with single-cell resolution, opening new avenues for single-cell analyses.

In addition to the application presented in this work, DynaMorph can be used with other cell types and other quantitative measurements of dynamic architecture to delineate morphodynamic states. For example, the physical measurements of cell density and cell anisotropy can be complemented with molecular measurements of metabolic states, cell cycle, or proteins of relevance to specific cell types. Furthermore, While we demonstrated this proof-of-principle on microglia and neural progenitor cells, this approach can be used to track cell transitions during cell differentiation and lineage specification, as well as broad cellular functions like viral infections and response and apoptosis.

Following are areas of improvement in DynaMorph:

- Segmentation and tracking accuracy are currently limited by the density of cell culture. The denser environment significantly increases the difficulty of recognizing individual cells. Methods that incorporate more human supervision or models based on geometric representations of shapes can better extract and analyze cell behavior in a dense environment.
- We found good agreement between the gene expression changes and morphodynamics in response to different perturbations at the resolution of cell population. However, direct correlation between the single cell morphodynamic behavior and its transcriptomic profile is still an area of future work. The transitions in gene expression and morphodynamics driven by the perturbations can be elucidated by paired measurements of gene expression and morphodynamics per cell. Post-imaging staining of cells using markers that discriminate gene expression clusters seen in single cell transcriptomes can provide this information.
- The biological meaning of morphodynamic states of microglia we have identified from average of feature vector across trajectories (Figure 4) is confounded by two possible causes: a) the cell populations subjected to the same perturbations exhibit heterogeneous dynamics, and b) the dynamic behavior in the latent shape space does not always align with existing vocabulary of cell behavior. The heterogeneity in cell behavior can be distinguished by acquiring data at higher time-resolution or utilizing a ‘bag-of-features’ of latent shape vector rather than averaging the shape vector ((5)), which is an area of future work. How to ascribe a biological interpretation to the learned representation is an open question for the field. A mathematically rigorous and experimentally generalizable representation can be considered a quantitative vocabulary with which we describe the dynamics of cells.

Taken together, DynaMorph is a versatile and quantitative framework for automated discovery of morphodynamic states of cells from live imaging data. Extraction of shape and motility descriptors can be coupled with a broad spectra of downstream tasks, e.g., anomaly detection, unsupervised state discovery, and supervised phenotype classification. Further integration with proteomic and transcriptomic assays can enable a more complete understanding of cell types and cell states. Identifying morphodynamic states in the context of disease could help elucidate the natural and induced changes in cell behavior, potentially informing novel therapeutics or diagnostics.

## Supporting information

Video Set 1

Video Set 2

Video Set 3

Video Set 4

Video Set 5

Video Set 6

## Methods

### Label-free brightfield, phase, and retardance imaging

We acquired all data using a Leica DMI-8 inverted widefield microscope, with 20x objective magnification at 0.55NA (air), and 0.4NA condensor on a Hamamatsu Flash-4 LT camera (6.5 um pixels). The cells were held at a constant 37°C, 5% CO2 using the Okolab stage-top incubator (H101-K-Frame). Each of the five polarization states were acquired, in sequence, with 50 ms camera exposure, for each of 5 z-planes for a given field of view. For the unperturbed microglia time series (training set), we acquired 52 time points and 27 fields of view (9 per well) over 24 hours at 27 minute time intervals. For the perturbed microglia time series (test set), we acquired 159 time points, 9 fields of view (4 were used for analysis), over 24 hours at 9 minute time intervals.

Birefringence is an optical property of matter that describes a different refractive index for different orientation axes of polarized light. The differential phase shift caused by these refractive indices, and the orientation axes, allow us to decompose birefringence into two properties: retardance and orientation (respectively). Each of these properties can be represented in terms of a combination of the Stokes parameters, which are Cartesian projections of a vector from spherical coordinates that describes the full polarization state of light. With the application of Mueller matrices that are tuned to our instrument setup, we can translate the Stokes representation into image intensities and vice versa. Therefore, given a set of intensity images of the specimen with known polarization states, we can use the inverse model to compute the sample’s corresponding Stokes parameters, and from there compute physical properties of retardance and orientation. We estimate the background level of polarization with a 2D polynomial fit. Details for label-free hardware calibration, background correction and reconstruction can be found in previous work (16).

We used the open-source microscope control software Micro-Manager 1.4.22 (https://micro-manager.org/) for all image acquisition. The Micro-Manager plugin OpenPolScope (https://openpolscope.org/) performs the liquidcrystal-compensator (LCC) calibration by first finding voltages to achieve extinction (I_Ext_ , intensity minimum). A pre-defined “Swing” voltage (0.03) is applied to induce slight polarization ellipticity along one axis (I_0deg_). This “Swing” is empirically determined based on the sample, in order to produce maximum polarization contrast. Then, using the brent-optimizer minimization procedure, we find three more voltage states centered on I_0deg_ that sample other orientations (I_45deg_, I_90deg_, I_135deg_).

Following the reconstruction algorithm described previously (16) and documented on github (https://github.com/mehta-lab/reconstruct-order), phase reconstructions used the above hardware parameters plus the total variation regularizer with parameters: `rho=1`, `itr=50`, `absorption=1.0e-3`, `phase=1.0e-5`. Retardance reconstructions used default parameters plus “local fit” background correction, and retardance scaling of 1e4.

### Culture of primary microglia

De-identified tissue samples were collected with previous patient consent in strict observance of the legal and institutional ethical regulations. Protocols were approved by the Human Gamete, Embryo, and Stem Cell Research Committee (institutional review board) at the University of California, San Francisco.

Primary human microglia were obtained from second trimester (gestational week 18-22) cortical brain tissue using magnetic-activated cell sorting with CD11b magneteic beads. Tissue samples were dissected in artificial cerebrospinal fluid containing 125 mM NaCl, 2.5 mM KCl, 1 mM MgCl_2_, 1 mM CaCl_2_, and 1.25 mM NaH_2_PO_4_. Tissue was cut into 1 mm^3^ pieces, and the tissue was enzymatically digested using 0.25% trypsin (reconstituted from 2.5% trypsin, ThermoFisher 15090046) with addition of 0.5 mg/ml DNase (Sigma Aldrich, DN25) for 20 minutes at 37°C, mechanically dissociated and then passed through a 40 um mesh cell strainer (Corning 352340). The resulting cell suspension (100-150 million cells) was centrifuged for 5 minutes at 300 × *g* and washed twice with Ca^2+^/Mg^2+^ - free phosphate buffered solution with addition of 0.5 mg/ml DNase to prevent cell clumping. Cells were re-suspended in 900 ul of MACS buffer (PBS with 0.5% BSA) with addition of 0.5 mg/ml DNase and incubated with 100 ul of the CD11b magnetic beads (Milteniy Biotec, 130-049-601) for 15 minutes following the manufacturer’s instructions. After the magnetic beads incubation, cells were washed with 20 ml of PBS, span down at 300 × *g*, re-suspended in 0.5 ml of MACS buffer and loaded on a MACS LS column (Milteniy Biotec, 130-042-401). Cells on the column were washed three time with 3 ml of MACS buffer, the column was removed from the magnet, and remaining microglia cells were eluted in 5 ml of microglia culture media. Microglia culture media (50 ml) consisted of: 33 ml of phenol red-free basal media Eagle’s (BME), 12 ml Hank’s buffered solution, 1 ml B27 (ThermoFisher, A3582801), 0.5 ml N2 (ThermoFisher, 17502048), 2 ml of 33% glucose, 0.5 ml GlutaMax (ThermoFisher, 35050061), and 0.5 ml penicillin/streptomycin (ThermoFisher, 15240062). Purified microglia cells were span down at 300 × *g* and plated at 100 103 cells per well in microglia media supplemented with 100 ng/ml of rhIL34 (Peprotech, 200-34), 2 ng/ml TGFb2 (Peprotech, 100-35b) and 1x CD lipid concentrate (Lifetech, 11905031) to promote microglia cell survival in the monoculture. Prior to plating, glass-bottom 24 well plates (Cellvis, P24-1.5H-N) were coated with 0.1 mg/ml poly-d-lysine (Sigma-Aldrich, P7280) for 2 hours at room temperature followed by three double-distilled water washes and additionally incubated with laminin and fibronectin in PBS for 3 hours at 37°C. Cell culture media was changed twice a week, and was changed 24 hours prior to the start of the time lapse experiment to allow cells to re-equilibrate.

Time lapse imaging experiment was started on day 6 of culture and continued for 24 hrs in environmental control chamber (5%CO2, 37°C and relative humidity of 70%).

### DynaMorph Pipeline

*DynaMorph* is composed of a collection of machine learning/deep learning tools that operate on imaging data and automatically generate morphodynamic summary of target instances. Here we elaborate technical details of the preprocessing steps and DynaMorph pipeline. Note that two components of the pipeline required training in advance to state discovery applications: semantic segmentation model during preprocessing and the self-supervised encoding model. In this work we dedicated a separate set of experimental data on control microglia for model training and conducted majority of the analysis on perturbed microglia.

In order to identify and track microglia, DynaMorph first uses segmentation algorithms to extract individual cells from the imaging data. Without detailed annotation of the data, we built the segmentation method based upon the deep learning semantic segmentation model U-Net with weak supervision from human expert. It should be noted that a variety of similar deep learning based segmentation tools are proposed in recent literature, most of which will be fully applicable in our pipeline. For the application of DynaMorph, selection of segmentation tool is interchangeable as long as the segmentation results provide unbiased selection of target cell population.

#### Semantic segmentation

A single three-class U-Net classifier (43) was deployed for the cell semantic segmentation, whose training data was derived from a combination of human annotations and augmented background annotations. In the preparation of training data, we utilized interface from ilastik(44) and manually annotated microglia, contaminating non-microglia cells (mostly progenitor cells, also containing neurons, radial glia) and background/experiment artifacts in 30 frames selected from 13 fields of view. Manual annotation took in total approximately 8 hours, generating identifications for 1252 microglia and 312 non-microglia cells. The annotations were only partial, in which most of the background were not annotated (**??**B). To compensate for that (see Fig. 1 - supplement 3 for comparison), we utilized the internal segmentation module (random forest-based) of ilastik to separate foreground (microglia and contaminating cells) and background (under human supervision), which generates supplementary background annotations that could be used as data augmentation. 50 frames of ilastik-generated background annotations were inspected and used as additional training inputs.

All training frames (2048 × 2048 pixels): human annotated frames and augmented frames, were combined and cropped into small patches (256 × 256 pixels). Two strategies were utilized: a frame could either be cut into patch tiles by a sliding window with no overlap, or by random sampling centers and rotation angles to extract patches from the context. The second strategy served partially as a data augmentation technique and was designed to increase the diversity of cell orientation and position. No further augmentation was performed as all frames were imaged with the same scale and brightness.

U-Net was adapted from https://github.com/qubvel/segmentation_models, and a pretrained backbone of resnet34 (45) was used to initialize weights. The model was trained end-to-end with standard cross entropy loss and Adam optimizer (46). During training, weight scaling was employed to balance loss for different classes. Little hyperparameter search was performed due to limited amount of data and lack of validation metrics. Final model was evaluated visually based on criterions including quality of cell segmentation mask, prediction accuracy on microglia/contaminating cells, robustness against imaging artifact. It should be noted that *DynaMorph* uses segmentation results to extract single cell patches, for which task segmentation accuracy isn’t a major concern. We thus didn’t exhaustively validate and optimize segmentation performances to ease application-time usage. This step is also interchangeable with any off-the-shelf segmentation method/package that could provide unbiased selection of target cells.

To generate segmentation for a full video, we first separated the video into static frames. For each frame, the full field of view (2048 × 2048 pixels) was divided into patches (256 × 256 pixels) following the sliding window/tiling strategy, model was then applied on individual patches, results of which were tiled to generate the full prediction mask for this frame. To avoid edge effect, 20 repetitions with different offsets were performed, all 20 prediction masks were aligned and averaged to generate the final prediction.

#### Instance separation

We applied clustering on the pixels from segmentation masks to isolate and extract masks of each individual cell. In the procedure, all pixels predicted as foreground (microglia and progenitor cells) by U-Net were extracted, then clustered based on their 2D coordinates in the frame. We used DBSCAN (47) to detect core points of each cell as well as to exclude outlier points coming from prediction artifacts. Note that this is feasible when cells are cultured in a relatively sparse environment. In fields with densely populated cells, clustering would not be able to separate boundaries between overlapping/contacting cells, instead more robust end-to-end instance segmentation models (48, 49) would have better results. Implementation from scikit-learn was applied, the parameters of which were tuned to separate adjacent cells with small amount of contact. Masks with extreme sizes (*>*12000 pixels or *<*500 pixels) were excluded to avoid prediction/experimental artifact and overlapping/contacting cells that cannot be separated.

The identity of each cell was then determined based on the average classification score over its mask. We simply averaged the pixel-wise probabilities of both classes (microglia or progenitor cells) across the whole cell mask. The resulting ratio of probabilities represented the proportion/likelihood this specific cell is microglia or progenitor cells. Cells with microglia proportions larger than 0.9 were regarded as true microglia, and vice versa for progenitor cells. The rest would be regarded as ambiguous cells. These identities helped in determining validity of trajectories in the following tracking procedure.

#### Tracking

Within a full video, instance separation was performed on each static frame independently, results of which were used in this step to connect cells across time and form trajectories. The tracking procedure followed the classic work of (50), in which the core architecture is a linear assignment problem (LAP). In every two adjacent frames, each frame would have a list of cells generated from the instance separation step, along with their sizes and center positions calculated on the segmentation masks. Then a matching problem was set up between the two lists of cells, with a pairwise cost matrix calculated based on position displacements and shape differences between pairs of cells:

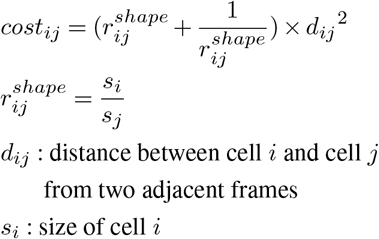

The cost favored cell pairs that were spatially close and had similar size. In practice, we also enforced thresholds (maximum displacement 100 pixels, maximum size difference 2 folds) to refine results. Resulting matching pairs along all frames in a video were connected, formulating a set of single cell trajectories that span multiple frames. We further adopted the gap closing strategy in (50) to connect short trajectories that were separated due to imaging/segmentation issues. A similar LAP set up with only squared distance cost was employed.

Identities of the trajectories obtained through the above procedure will be calculated by averaging static frame identities. Only trajectories with over 95% of all frames being labeled as microglia (proportion *>* 0.9) were kept for the main analysis. In total, 5731 microglia trajectories were collected, 3715 of which were extracted from the test dataset. Meanwhile, trajectories with over 50% of the frames labeled as progenitor cells formed another small collection of 125 cells, which were used in the extension analysis.

#### Self-supervised encoding with VQ-VAE

A variational autoencoder variant: VQ-VAE (31) was applied in this work to summarize shape of cells through an encoding-decoding process. The model was trained on microglia cell patches (phase and retardance channels) extracted from the training set and directly applied to the test set data to generate shape descriptors for cells under different treatment.

We took all individual microglia cells in static frames from the instance separation step and cropped patches of size 256 ×256 pixels around the cell centers. If the cell appeared on the border of a field of view, the out-of-border area would be filled with median background values of phase and retardance. Meanwhile, since we focused on analysis of single cells, we masked out other cells appearing in the patch area with median background values. Sample patches before the masking can be found in Figure 2. Samples after masking can be found in Fig. 2 - supplement 2 and Fig. 2 - supplement 5.

VQ-VAE was implemented in pytorch (51), using two deep convolutional networks as encoder and decoder respectively. Input patches were first down-scaled to 128 × 128 and normalized on both channels. The VAE encoder further down-sampled the image by 64 folds, resulting in a latent vector for each cell with shape of 16, with channel as the last dimension. Then, consistent with the standard setup of VQ-VAE, we regularized latent space by forcing a discretization step on the channel dimension, in which each vector at a given position out of the 16 × 16 grid was matched with 64 embedding vectors and the closest embedding was inserted back to the position and passed to the decoder. A matching loss was enforced in the later phase of training that minimizes frame-to-frame differences between latent vectors of the same cell along its trajectory. A combination of these two losses was used to train the model in an end-to-end fashion.

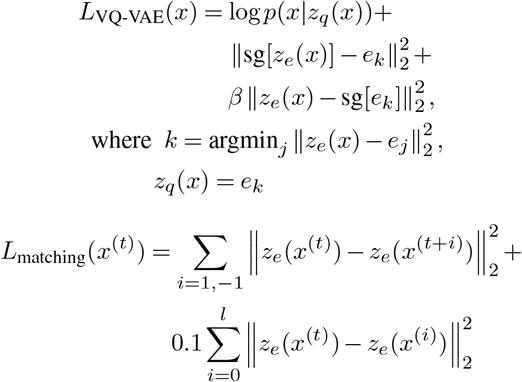

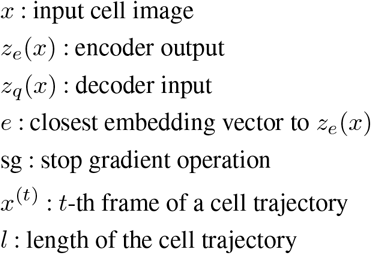

where *β* = 0.25 controls the amount of commitment loss regularized during training. In the training procedure, we applied only *L*_VQ-VAE_ in the initial phase. After 100 epochs (or when the model could generate reasonable reconstructions), we added in *L*_matching_ with a weight ratio of 0.0005 and trained the model till convergence.

Size of the latent space used in this study was consistent with the compression fold in its original work (31). Note that we didn’t push further to a much smaller latent space (i.e. *<*100D) due to the concern that smaller latent vector will consist of more conceptualized and higher-level representations of images, which would cast difficulties in interpretation and quantification.

Note that matching loss was only applied within mini-batch to avoid extensive calculation. During training we enforced a sample order that aligns most of the neighboring/same-trajectory cell pairs sequentially, so with a high chance a cell patch’s next or previous frame would exist in the same mini-batch.

#### PCA of latent vectors

PCA was fitted on *z*_*e*_(*x*) for all microglia cells in the training set, and applied to all training and test cells. Implementation from scikit-learn was used, and we extracted PCs that explain the top 50% of all variances: 48 components were extracted, explained variance of which is visualized in Fig. 2 - supplement 7A.

We also tried fitting and transforming *z*_*q*_(*x*) (data not shown), which generated very similar results and same interpretations for the top PCs. Distance metrics defined with either *z*_*e*_(*x*) or *z*_*q*_(*x*) showed similar results (Fig. 2 - supplement 9) as well. But since *z*_*e*_(*x*) contained more information before the quantization, we employed it throughout this work.

Other dimensionality reduction methods including tSNE (52) and UMAP (35) were also tested and showed no advantage over PCA as no clustering patterns were detected.

#### Gaussian Mixture Modeling of Trajectory Feature Vectors

To further unravel the morphodynamic features, as well as to link transcriptomic profiles of microglia, downstream clustering was performed on cell trajectories. Individual cell TFVs were composed of a shape descriptor and a motion descriptor. Top 48 PCs derived from PCA of latent vectors were used as the shape descriptor. Distance traveled per frame along the trajectory were averaged and its log mean taken as the motion descriptor. The log mean distance values were scaled to keep variance comparable with the top 2 PCs.

Gaussian Mixture Model (GMM) is utilized in the analysis to discover discrete states from TFVs. As GMM is capable of allocating different prior distributions to different perturbations, it allows the interpretation of morphodynamic differences between perturbation conditions as changes in the weight of each mixture component. We chose to use a 2-component GMM after testing clustering robustness (measured by Silhouette score) against number of components.

More specifically, we assumed two mixture components (two states) for microglia, and cells are sampled from one of the two components based on a Bernoulli distribution dependent on the perturbation condition. Given the TFV *x*_*i*_ and treatment group *y*_*i*_ of a cell, it has marginal probability as:

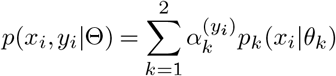

where 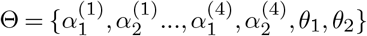 were parameters of the mixture model: *θ*_*k*_ is the center of *k*-th component, 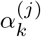 is the weight of *k*-th component in condition *j*. In the two-state model, 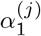 and 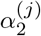 (summed to 1) formed the Bernoulli distribution for condition *j*. In the test set 4 conditions were used so 4 sets of component weights were fitted with samples from the corresponding groups.

Expectation-Maximization (EM) algorithm was employed to iteratively evaluate the parameters of GMM. E-step evaluates the membership weight of each cell based on the current set of parameters:

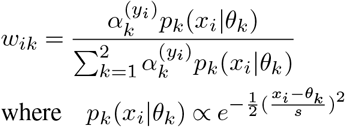

M-step recalculates the parameters based on new membership weights:

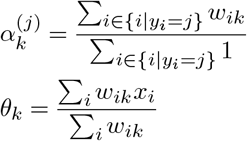

Two steps above were repeated till parameters converged. In the procedure we fixed the variance of mixture components to be the same as variance of TFVs to increase stability.

### scRNA Sequencing

Cells from three conditions (control, IFN beta and GBM supernatant) were dissociated using 0.25% trypsin solution, labeled with Multiseq barcodes (53) and processed for single cell mRNA sequencing using the 10x Genomics Chromium v3 Platform. CellRanger version 3 was used to generate the count matrix of cells by genes. Cells with more than 20% mitochondrial abundance or less than 500 UMI were removed. Doublets were removed during the demultiplexing of Multiseq barcodes.

SCTransform (54) was applied to the raw count matrix followed by PCA on the residuals. The top 30 principal components were used to find neighbors for Leiden clustering (55) and the UMAP (35) projection.

## Code and Data Availability

Open source python software for reconstruction of label-free optical properties is available at https://github.com/ mehta-lab/reconstruct-order and for analyzing cell states is available at https://github.com/czbiohub/dynamorph. The README file of dynamorph repository provides the link to example data.

## Acknowledgements

We thank Greg Huber for discussions around the physical models of cell states. J.Z. is supported by NSF CCF 1763191, NIH R21 MD012867-01, NIH P30AG059307, NIH U01MH098953, and grants from the Silicon Valley Foundation and the Chan-Zuckerberg Initiative. G. S. is supported by NIH NRSA F32 1F32MH118785. B. B. C., S-M.G., L-H. Y., and S.B.M. are supported by the Chan-Zuckerberg Biohub’s intramural program. This collaboration was supported by the Chan-Zuckerberg Biohub’s intercampus award to T. N. and J. Z.

## Supplementary Notes

### Encoding with partial inputs

Inputs from two channels (phase and retardance) were used to generate the shape embedding of individual cells in this study. To validate the general applicability of our method, we apply VQ-VAE on the same set of training/test cells but with only individual channel input of phase, retardance or bright field. Loss curves along the training process were plotted in Fig. 2 - supplement 4A.

Quantitative analysis on latent space suggests that phase and retardance channel could (when used individually) provide interpretable embedding for microglia in terms of both link with geometric properties (Fig. 2 - supplement 4B) and dynamics along trajectories (Fig. 2 - supplement 4C), while bright field images were more challenging to reconstruct and led to less interpretable results. Correlation matrices of the single-channel-input models indicate that top shape modes from phase or retardance models are capturing the same set of geometric properties as joint model. Bright field model captures the channel intensity, while being vague on other properties. In the distribution of pairwise frame differences, comparing with the median of random selected frames, joint model has the most stable embedding for same cell trajectories. All models retain the similarity hierarchy as discussed in Fig. 2 - supplement 9 to different extents.

Differential study on perturbations were performed in the same fashion. Feature vectors from phase or retardance inputs generate similar GMM clustering outcome as the joint model (Fig. 2 - supplement 4D), while bright field inputs struggle to separate two components between different perturbations. In the cross validation task of predicting perturbation treatment (Fig. 2 - supplement 4D), we see joint model has the highest overall accuracy and bright field model has the lowest.

Overall, we tested the single-channel-input models against the joint (phase+retardance) model used in this work from different aspects. Results suggest that phase or retardance model could provide cell embeddings that achieve the same goal as joint model but with slightly inferior quality. Bright field model provides less informative embedding and could have different downstream analysis performances.

### Comparison with other autoencoder variants

Following the same training/testing pipeline, we evaluated performances of two other representative autoencoder-based models, namely variational auto-encoder(56) and adversarial auto-encoder(57), on the reconstruction and interpretation tasks.

Loss curves during training are shown in Fig. 2 - supplement 3A, in which VAE and VQ-VAE has similar training behavior and converge after 300 epochs. AAE in contrast is much more unstable due to its adversarial model structure. AAE initially had relatively lower reconstruction loss but with much higher frame matching loss, suggesting a strong overfit. After inclusion of frame matching loss, we stopped its training after both loss achieved a relative plateau. During the whole process, discriminator accuracy was between 60 80%. Fig. 2 - supplement 2 illustrated some sample reconstructions from the three models, VQ-VAE has overall best reconstruction performance.

Fig. 2 - supplement 3B and C demonstrated evaluations on embedding quality based on shape interpretability and dynamics along trajectories. VAE has similar performance on the correlation matrix, though both models are worse in terms of embedding stability on frames from same cell.

For differential study on perturbations, GMM outcomes from model embeddings are very close (not shown). Fig. 2 - supplement 3D presents the confusion matrix for permutation treatment predictor, on which both models have worse accuracy than VQ-VAE.

Overall we find that these other auto-encoder variants are suitable for encoding the shape and discovery of cell states, but VQ-VAE provided better reconstruction and embedding quality.

### Predicting treatment conditions with Trajectory Feature Vectors

Given the TFVs built upon shape and motion of microglia, we explore the possibility of predicting perturbation condition purely based on microglia’s morphodynamic response. In this task, TFVs are used to predict the perturbation condition from which the cell is cultured. A 10-fold cross validation strategy is used, and gradient boosted tree is selected for the prediction task after comparing multiple machine learning models. During the evaluation, model (with default parameter) is trained with 9 folds of cell trajectories from the perturbation experiment and evaluated on the remaining fold (randomly split). In total 10 models are trained and results are aggregated to generate confusion matrix reported in Fig. 3 - supplement 2.

Among all 4 perturbation conditions, model has the best accuracy in control subset cells, while the other 3 disease-relevant conditions are more intertwined. Confusion between control subset and IFN beta subset is the lowest, indicating a large morphodynamic difference between these two groups. This is also reflected by the GMM clustering outcome as the two subsets are composed of cells of different states. Both GBM and IL17 subsets have 20% cells being predicted to come from control group, which hint these could be the proportion of microglia not activated by the stimuli.

Overall accuracy of this prediction task is 0.58 when using shape features from VQ-VAE with phase and retardance channel inputs. This could be compared with other model setups reported in Fig. 2 - supplement 4D and Fig. 2 - supplement 3D. As this task evaluates how embedding differentiate between perturbations, the accuracy score could partially reflect the embedding quality.

### Correlation between morphodynamic states and transcriptomic clusters

Two morphodynamic states were detected through the GMM clustering step in DynaMorph. We compared this result with the transcriptomes of cultured microglia to identify the key molecular sub-types that could correspond to the two states. In this experiment, cells from three conditions (control, IFN beta and GBM) were processed for single cell mRNA sequencing. Note that the majority of the measured cells (∼90%) are from control subset.

We generated UMAP projections of both morphodynamic features (Fig. 4 - supplement 3A) and scRNA-seq results to parallel the comparison, which are plotted in Fig. 4 - supplement 3B, C, D. A similar two-component clustering was performed on scRNA data, using Leiden clustering (55). The distributions of state/cluster assignments in each subset are plotted as bars in Fig. 4 - supplement 3E. Notably, cells from the control subset form a large group (cluster 2) and three small isolated groups (combined as cluster 1), in which the isolated groups could be further separated and interpreted through a more fine-grained clustering procedure. (Fig. 4 - supplement 4A) Cluster 1 accounts for ∼15% of all control cells, this relative abundance ratio coincides with the distribution of cells in states 1 and 2 defined on the morphodynamic space. Interestingly, exposure to IFN beta, which is enriched for morphodynamic state-1, also shows enrichment in one of the sub-group (1-2 in Fig. 4 - supplement 4) of transcriptomic cluster 1. In contrast, microglia that were treated with supernatant from GBM cultures, are transcriptomically more similar to the majority of control microglia, consistent with more subtle differences in morphodynamic states.

To address this further, we identified the top differentially-expressed genes (Fig. 4 - supplement 4B) for each transcriptomic cluster. Cluster 1-2, which is one the minor groups in control subset and enriched for cells exposed to IFN beta, has up-regulated genes encoding antiviral proteins (RSAD2, MX1, etc.) and interferon-induced proteins (IFIT1, IFIT2, IFIT3, etc.). Another cluster 1-1 is also enriched for cytoskeleton and microtubule-related genes (MAP1B, STMN1, TUBA1A, etc.), which would usually cause structural changes in cells. We hence speculate that the observed shape and density differences between different morphodynamic states could be correlated with the corresponding cell’s expression changes.

Note that this analysis only focuses on population level agreement between morphodynamics and gene expression profiles. A single cell level validation of the discovery: behavior/morphodynamic change paralleling transcriptome change will be an important aspect of our future work.

## Supplementary figures

**Fig. 1 - supplement 1.**
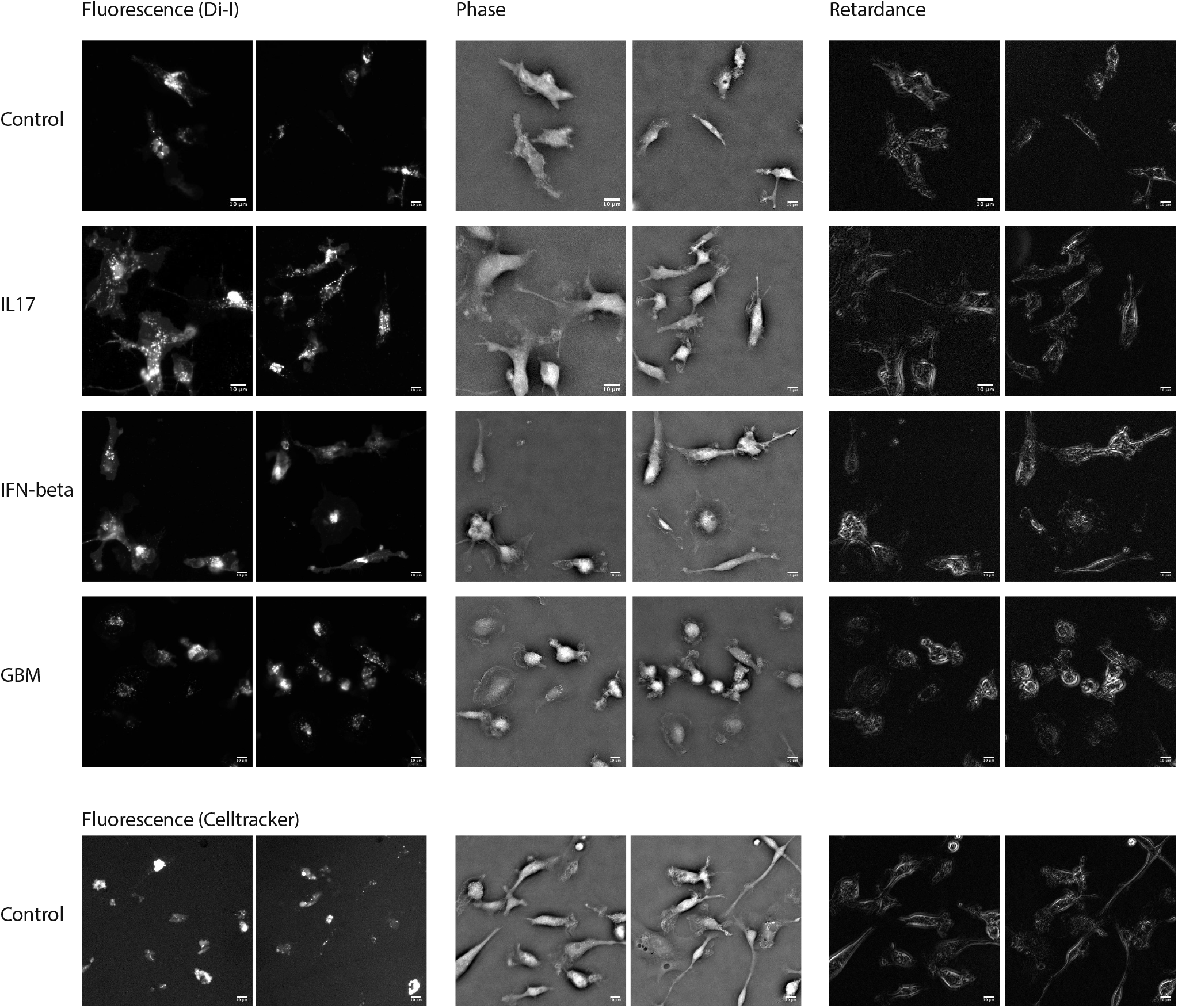
Comparison of microglia shape as visualized with vital dyes (Di-I and CellTracker CM-DiI) and label-free measurements (phase and retardance): Two examples from each condition are shown. Di-I and CellTracker are a lipophilic carbocyanine derivatives that internalize through different protocols, but should penetrate the membrane and persist in the cytoplasm enough to outline cell contours. Both dyes under all perturbation conditions show similar internalization and sequestration of the dyes. All scale bars are 10 um.

**Fig. 1 - supplement 2.**
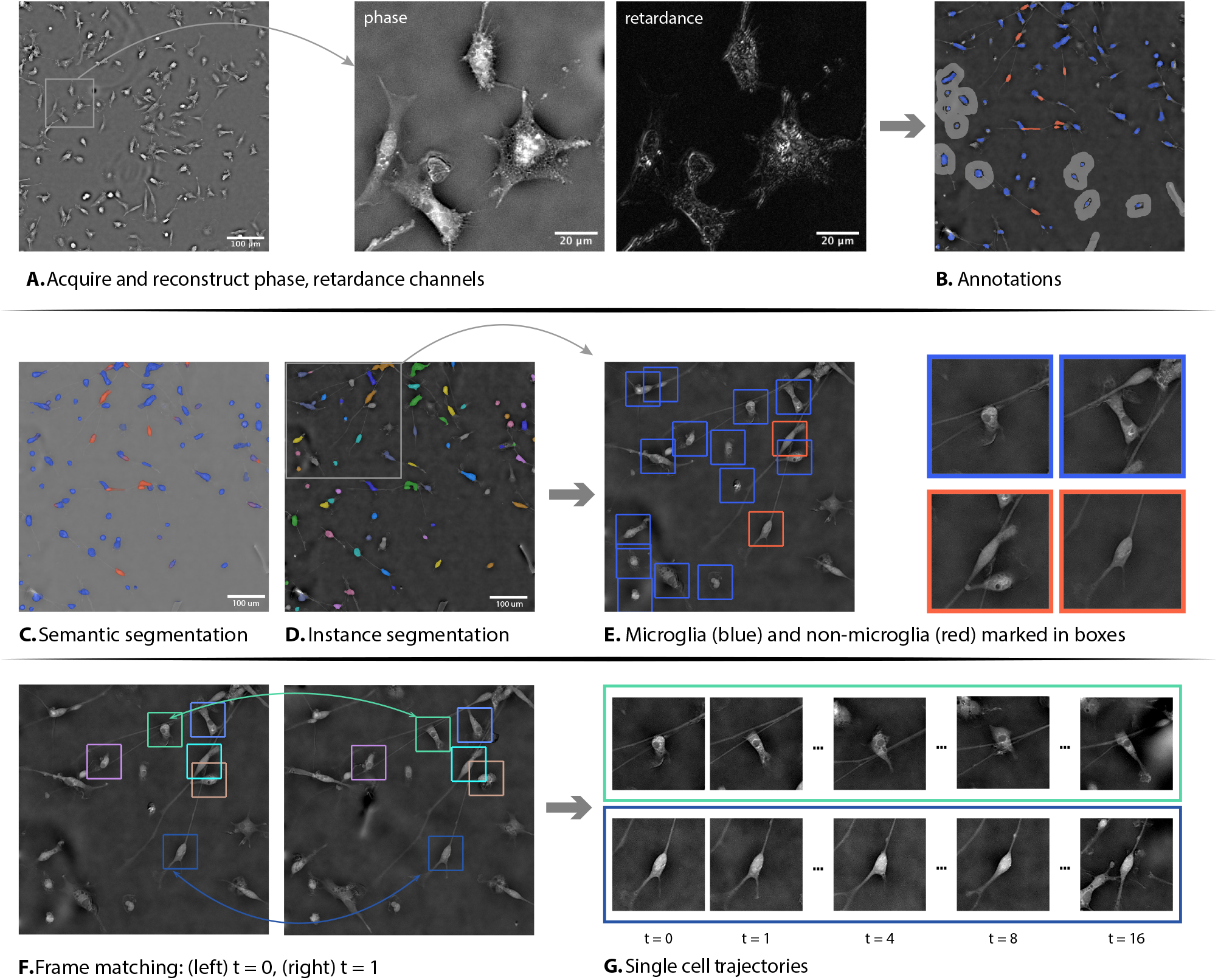
Identification and tracking of microglia with sparse supervision: **(A)** Cell shape and dynamics are visualized without label based on density (phase) and structural anisotropy (retardance). **(B, C)** Sparse human annotations enable microglia segmentation using a combination of conventional machine learning (ilastik/random forest) to separate foreground and background, and deep convolutional networks (U-Net) to separate microglia and contaminating non-microglia cells. **(D, E)** Segmentation masks are further clustered through DBSCAN to form individual cell masks, from which the position and surrounding patch of the cell are extracted. In the encoding pipeline we extracted boxes of 83*µm* 83*µm* around the cell centers for the patches. Note that stringent filtering was applied on the cell masks to exclude potential artifact and overlapping/contacting cells, leaving only masks/bounding boxes of individual cells. **(F)** Individual cells in adjacent time-points are linked through solving a linear assignment problem. **(G)** Connected components through multiple frames form single cell trajectories. Representative trajectories for microglia (top) and progenitor cells (bottom) are shown.

**Fig. 1 - supplement 3.**
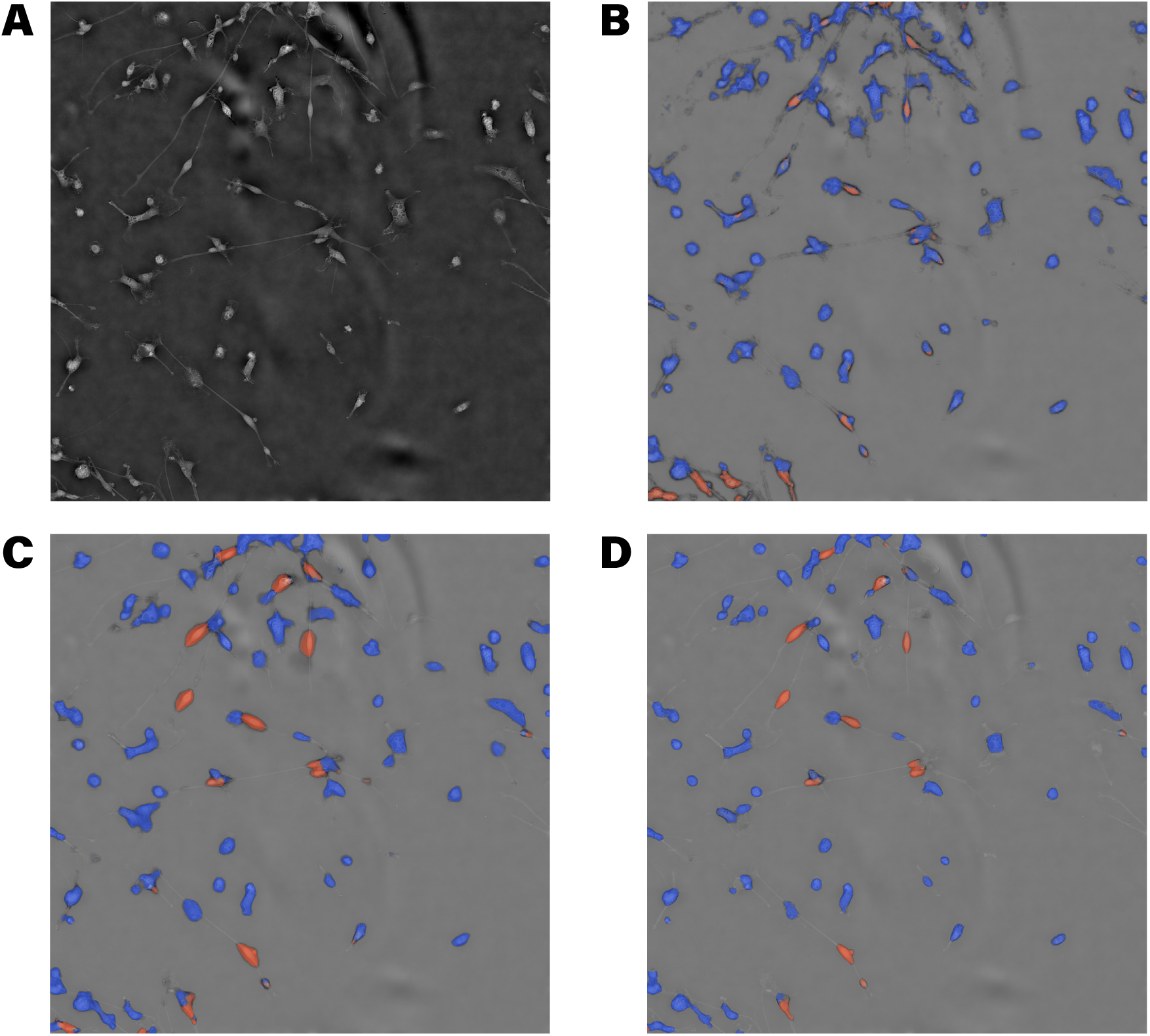
Comparison of different segmentation results: **(A)** Raw input of an unannotated slice, phase channel. **(B)** Prediction map from a random forest classifier (ilastik) trained on human annotations. Note that predictions are more noisy and chimeric. **(C)** Prediction map from a U-Net trained on human annotations. Note that most edges exceed the boundaries of cells. **(D)** Prediction map from a U-Net trained on human annotations and random forest augmented background annotations.

**Fig. 2 - supplement 1.**
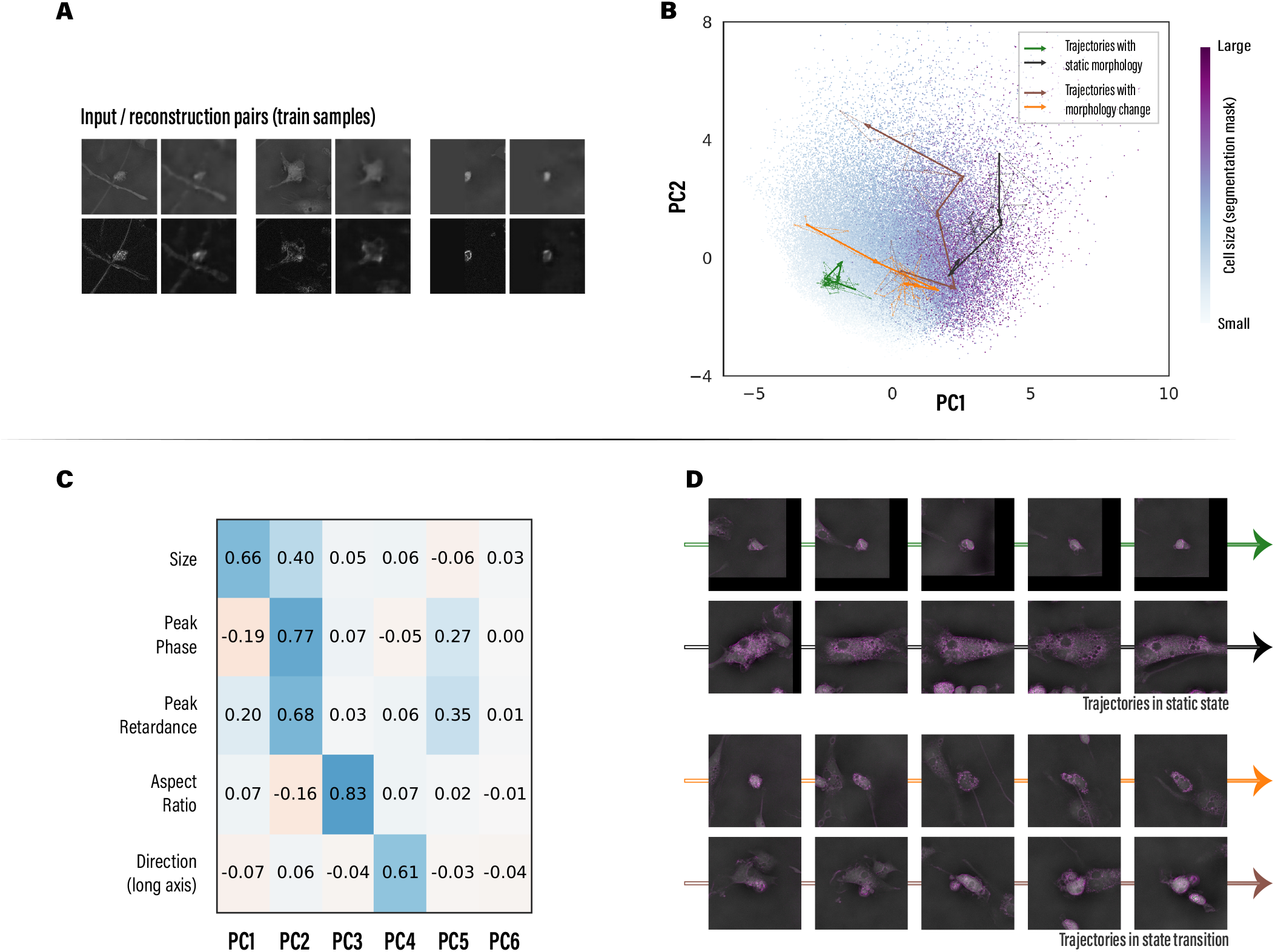
Reconstruction and encodings of samples from the training set: **(A)** Example patches from the training set and their reconstructions by VQ-VAE **(B)** Top 2 PCs of all samples from the training set, 4 representative trajectories are plotted. **(C)** Spearman’s rank correlation coefficients calculated between top PCs and geometric properties. Note that results are highly similar to test set samples. **(D)** Representative trajectories are visualized.

**Fig. 2 - supplement 2.**
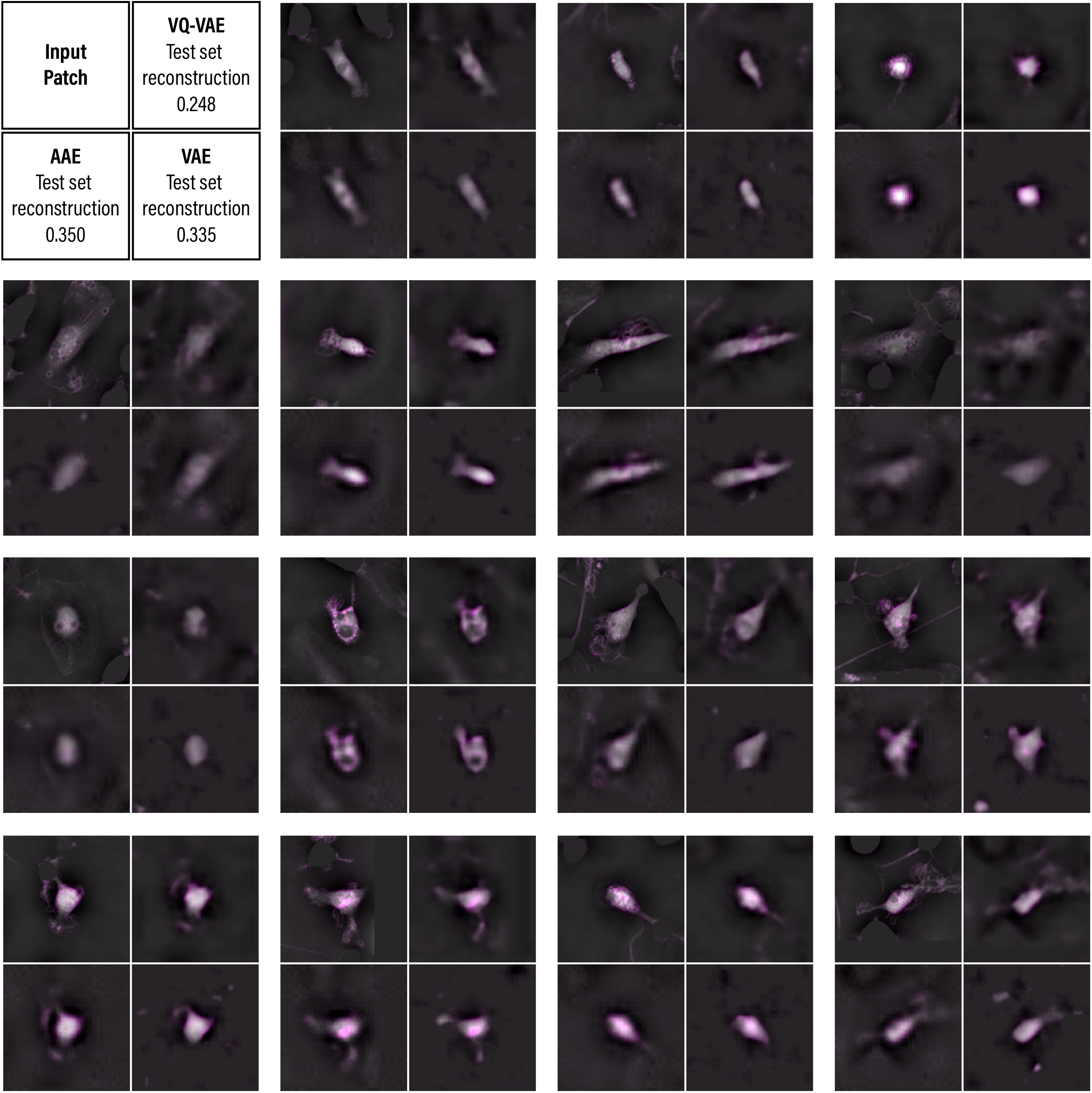
Sample cell patches randomly selected from the test set and reconstructions from auto-encoder variants: Each group contains 4 patches, respectively the original input and reconstructions from VQ-VAE, VAE and AAE. Order of the patches and reconstruction loss averaged over all test set data are noted in the top left panel. Note that all variants have latent space of the same size, and VQ-VAE has overall best reconstruction and image compression fold.

**Fig. 2 - supplement 3.**
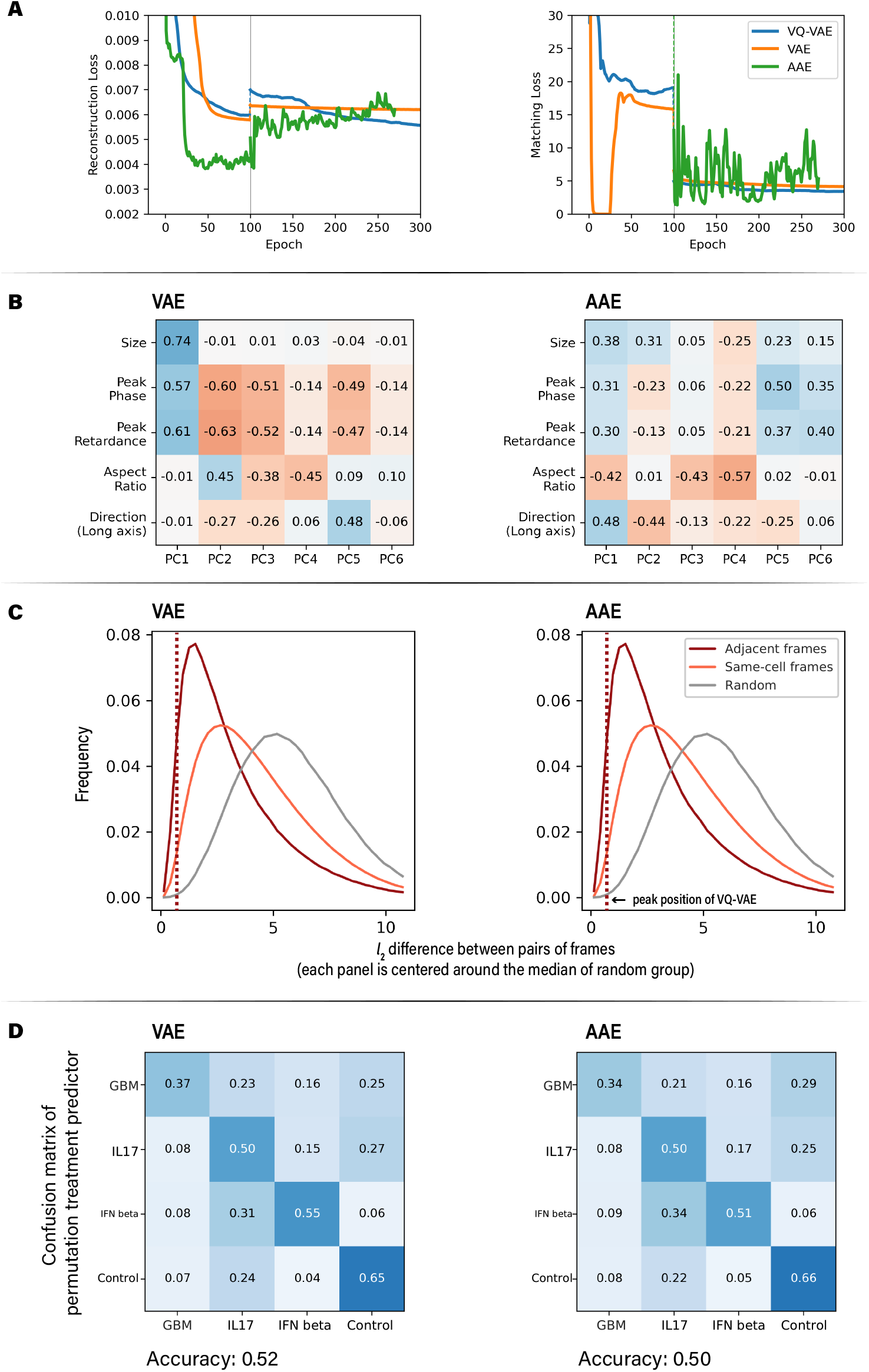
Comparison of different auto-encoder variants, compare with “phase+retardance model” plots in Fig. 2 - supplement 4 for VQ-VAE performance. **(A)** Reconstruction and frame-matching loss curves during training. Grey vertical lines indicate the inclusion of frame matching loss. VAE and VQ-VAE converged after 300 epochs. AAE had unstable loss due to its model structure. **(B)** Correlations between top PCs and selected geometric properties of the test set data. **(C)** Distributions of latent vector differences between pairs of frames. Both panels are centered at the median of random group, red dashed lines indicate the peak position of VQ-VAE (phase+retardance) model. **(D)** Results on differential study of perturbations: confusion matrix for the perturbation condition predictor (VQ-VAE has accuracy 0.58).

**Fig. 2 - supplement 4.**
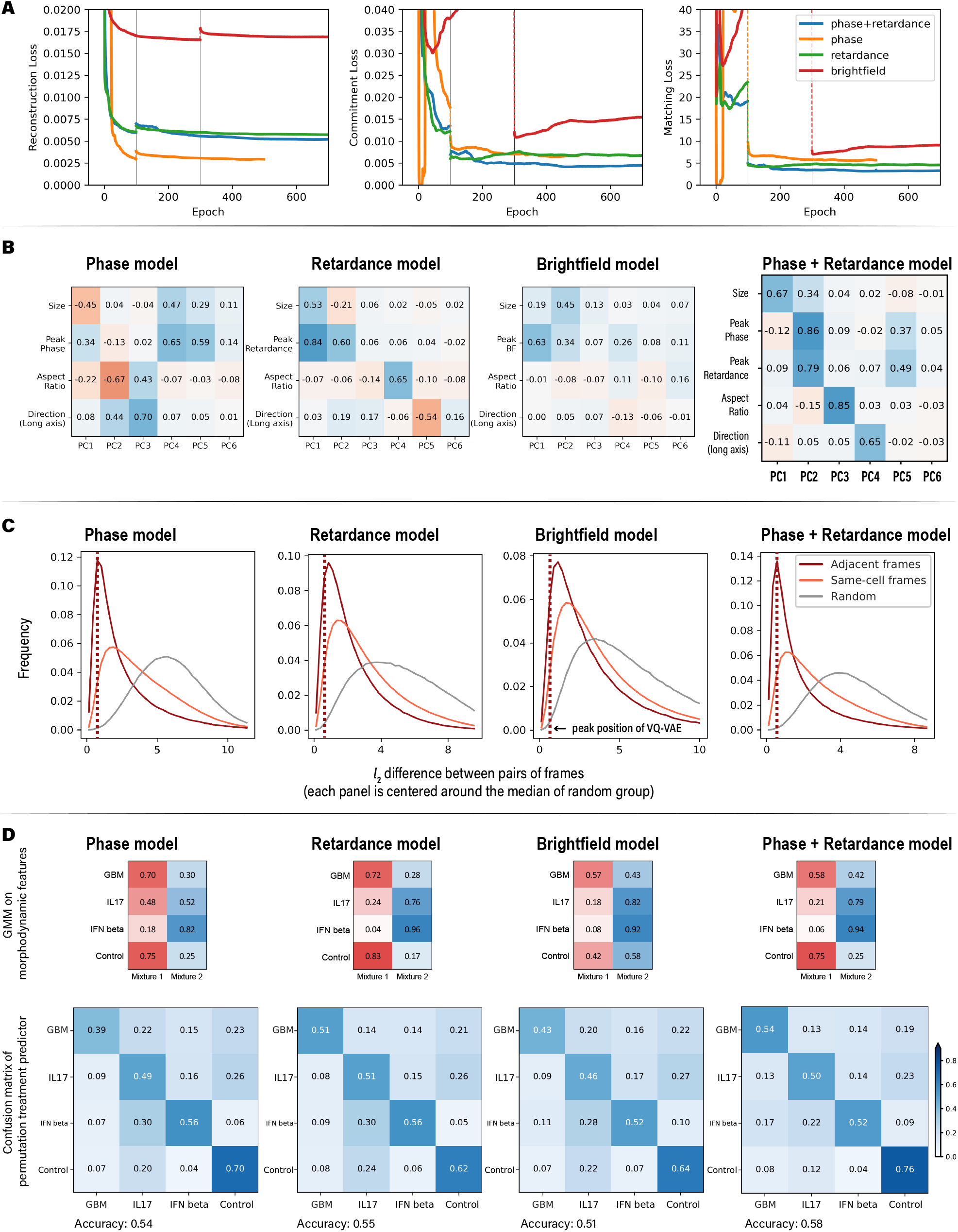
Comparison of models trained with individual channel inputs. **(A)** Reconstruction, commitment and frame-matching loss curves during training. Grey vertical lines indicate the inclusion of frame matching loss. All models reached convergence after 600 epochs. Models with either phase or retardance channel input reached similar level of loss as the joint model. **(B)** Correlations between top PCs and selected geometric properties of the test set data. (same plot as in Figure 2D) Top shape modes from single-channel-input models found relations with one or more geometric properties. **(C)** Distributions of latent vector differences between pairs of frames (same plot as in Fig. 2 - supplement 9). Note that all 4 panels are centered at the median of random group, and red dashed lines indicate the peak position of joint model. **(D)** Results on differential study of perturbations: distribution of detected GMM components (top row), confusion matrix for the perturbation condition predictor calculated in the same procedure as Fig. 3 - supplement 2 (bottom row).

**Fig. 2 - supplement 5.**
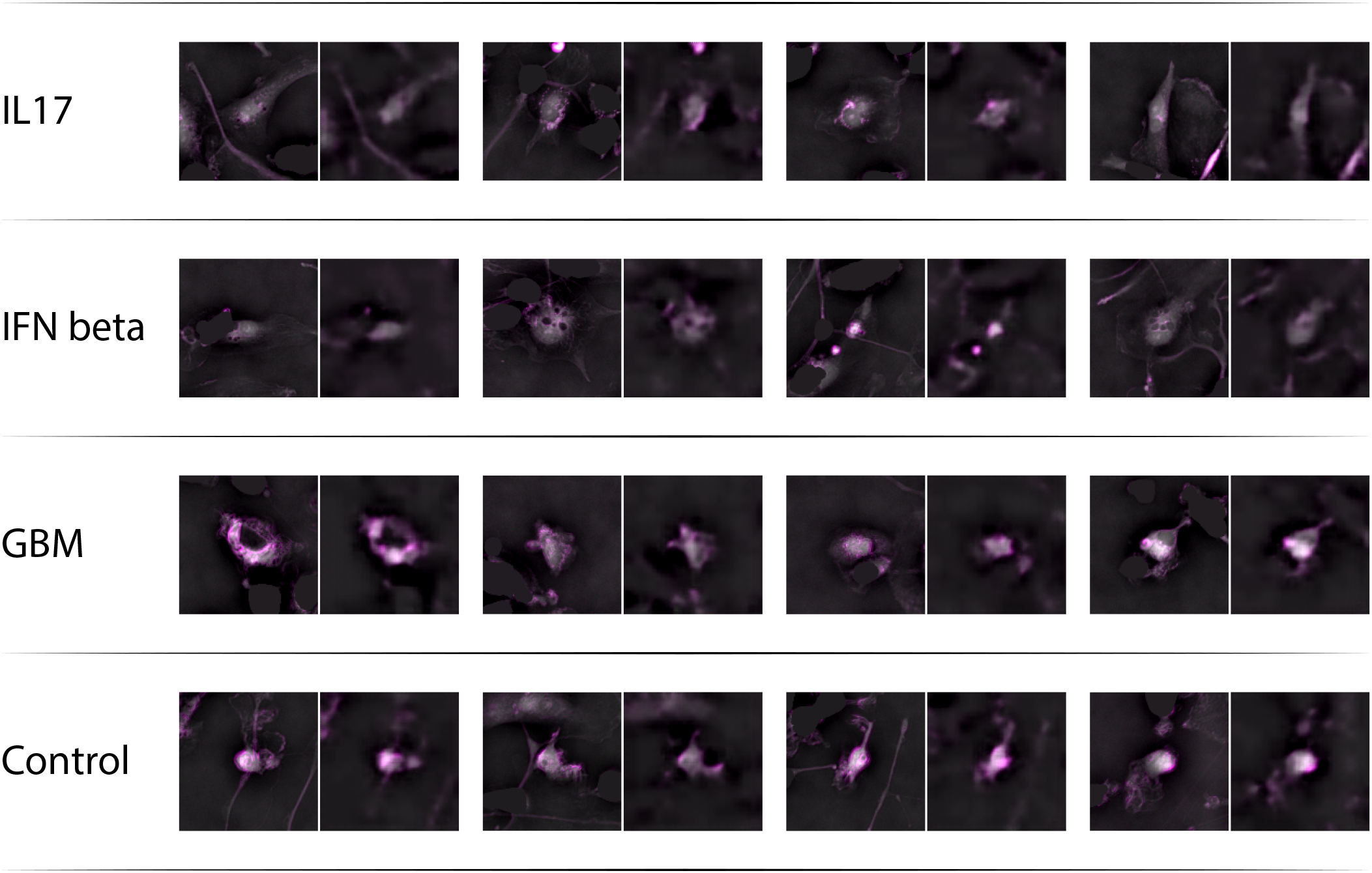
Illustrative cell patches of cells imaged under different perturbations (IL17, IFN beta, GBM, Control) and their reconstructions by VQ-VAE.

**Fig. 2 - supplement 6.**
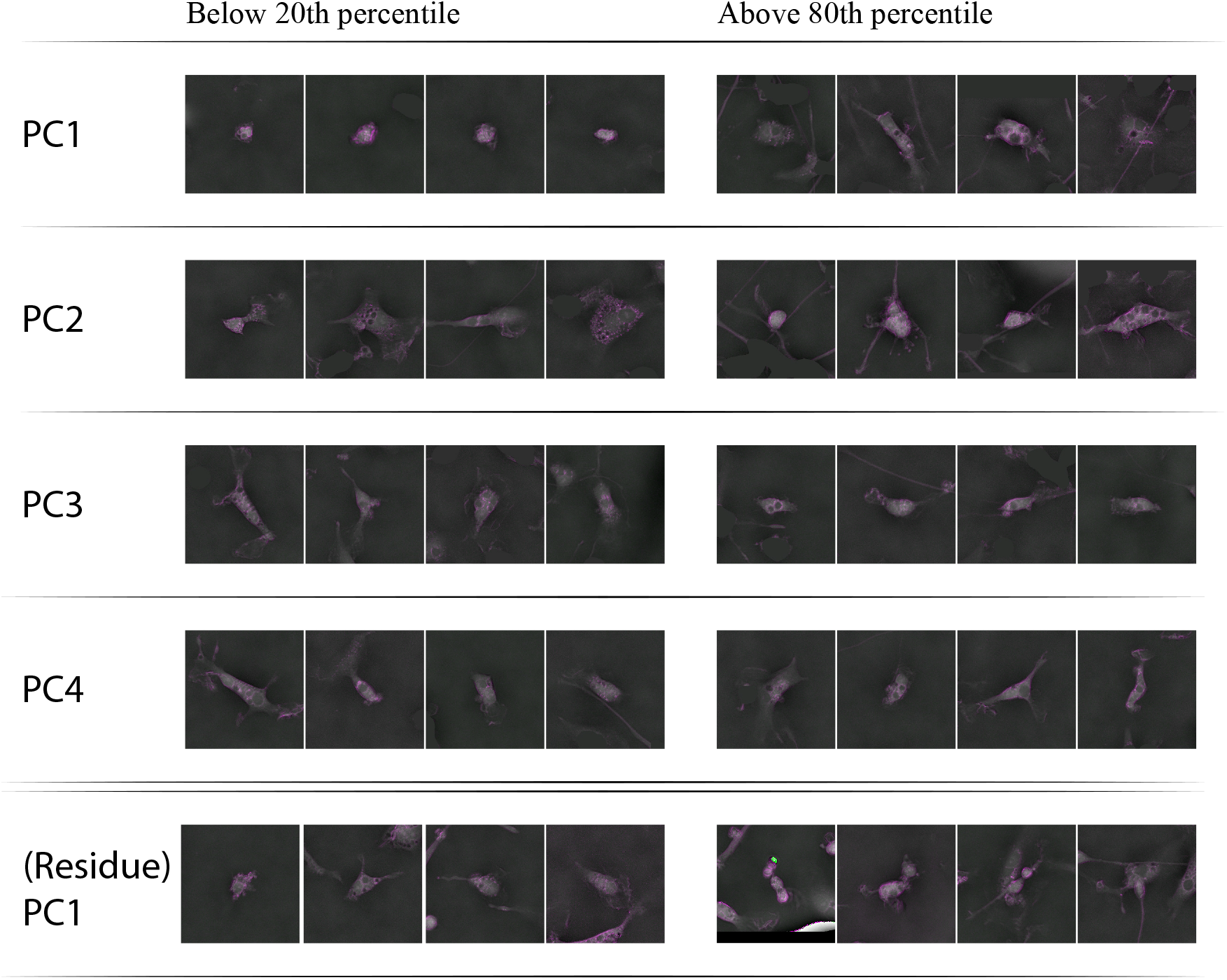
Cell patches randomly sampled from the upper and lower 20 percentiles along each principal component axis (PC1 to 4, residue PC1). Patches are visualized as combination of phase channel (as grey scale) and retardance channel (as magenta shades). Samples along PC1 show different cell sizes; samples along PC2 show different phase strengths; samples along PC3 and PC4 (controlled for PC1 and PC2 values) show different orientations. We further calculated a residue latent space after excluding all known geometric properties of cells (listed in Figure 2D), samples along the first principal component of the residue space (rPC1) are illustrated in the last row: samples with lower rPC1 values tend to involve less contact/interaction, while higher rPC1 value correlate with complex environments including overlapping cells, cell on the border of imaging view, imaging artifact, etc.

**Fig. 2 - supplement 7.**
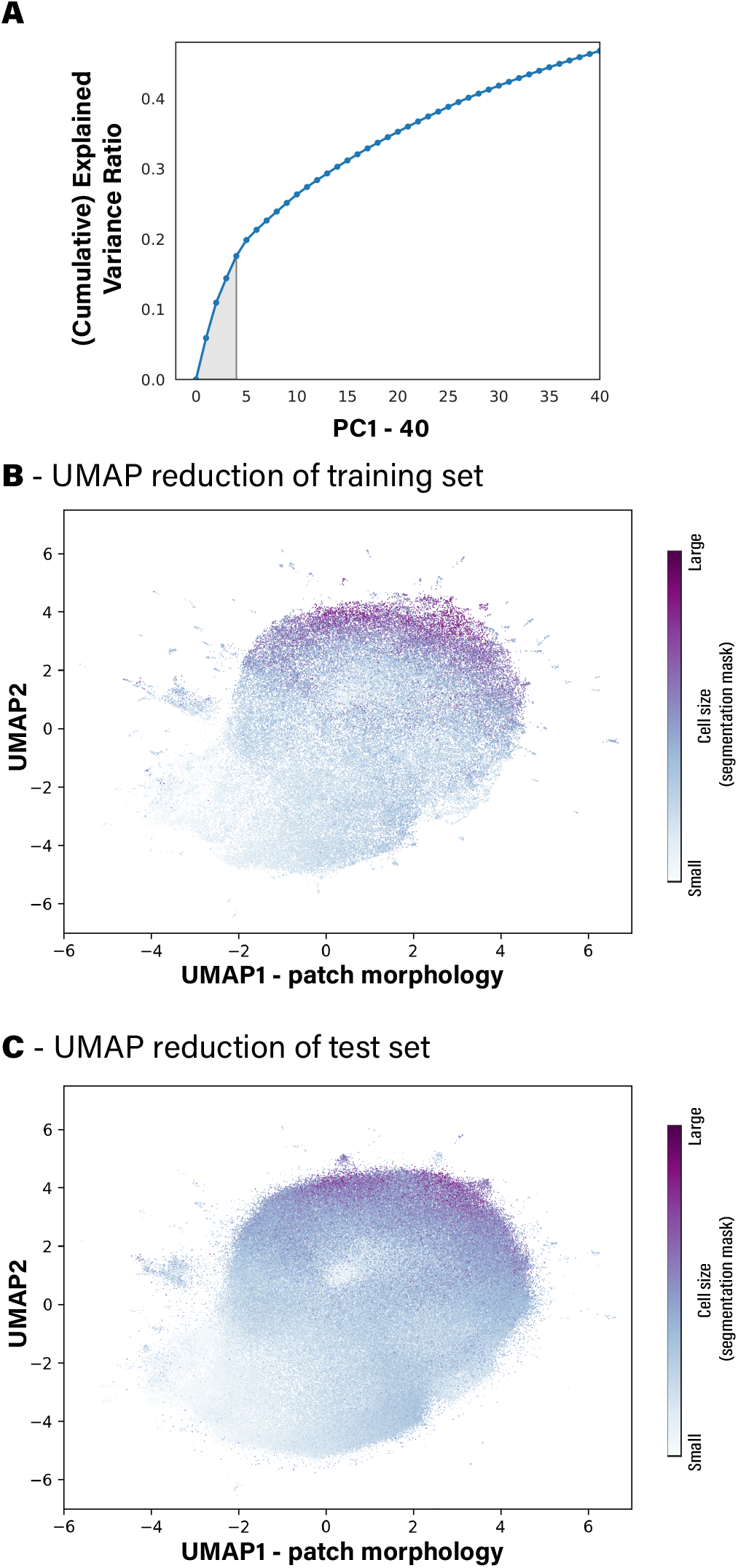
Learned Morphologial Space is high-dimensional: **(A)** Explained variance ratios of the top 40 principal components of the shape space. Note that analysis in this work focuses only on the first 4 principal components. (shaded area, explained 17% of all variance). **(B)** and **(C)** UMAP reduction (2 components) on sample latent vectors from the training set and the test set. No clustering patterns are found after tuning parameters, indicating that the space is highly continuous.

**Fig. 2 - supplement 8.**
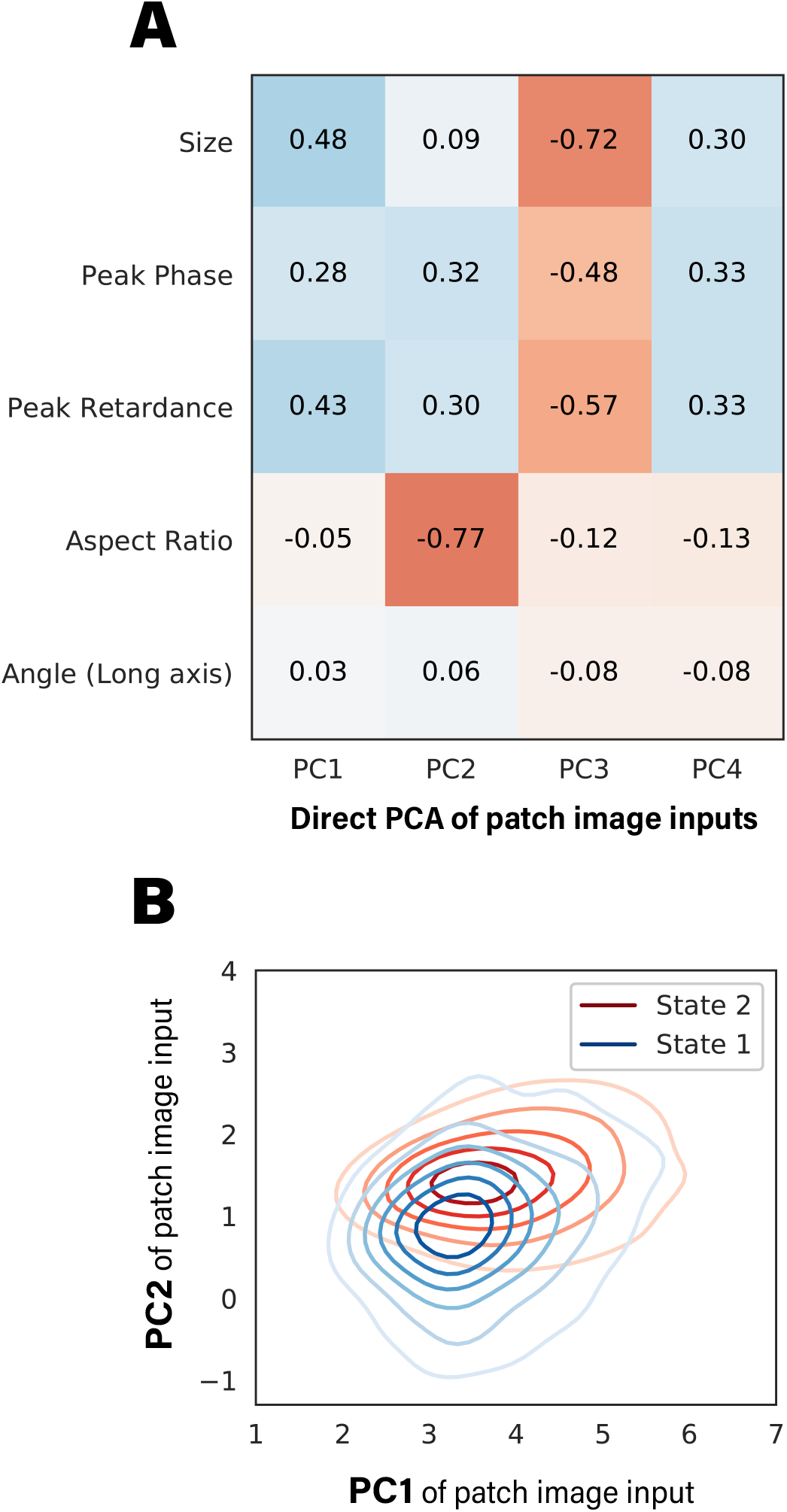
PCA was directly fitted on input images without encoding from the training dataset and applied to test samples. **(A)** Resulting PCs were analyzed for correlations to geometric properties. Spearman’s rank correlation coefficients were calculated between the same set of geometric properties and top image PCs. Correlations are weaker and more interrelated and thus harder to separate and interpret. **(B)** Two GMM states were defined using motion descriptor (speed) and image PCs. KDE plot of trajectories from the two states is illustrated, note that state separation is much worse than GMM states defined with latent vector PCs.

**Fig. 2 - supplement 9.**
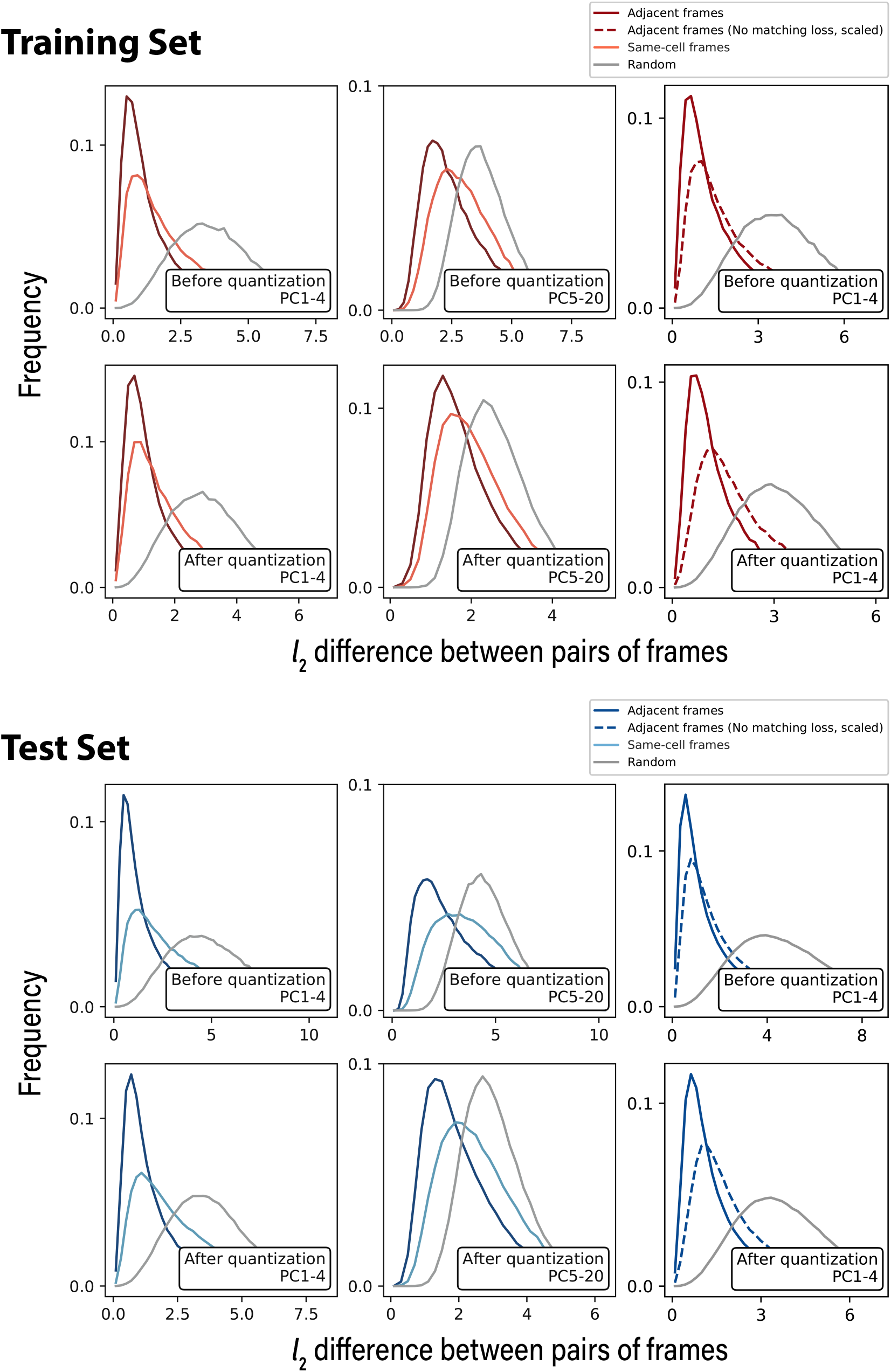
Distributions of latent vector differences between pairs of frames: On both training set (upper panels) and test set (lower panels), we calculated differences between pairs of frames for three conditions: adjacent frames in trajectories (deep red/blue), non-adjacent pair of frames from the same trajectory (light red/blue) and random pairs of frames selected from all static patches (grey). Left and middle columns show that distances in the latent space match well with our prior of shape similarities (adjacent > same-cell > random pair), and such trend is much more significant in top PCs. Right column compares the distribution between latent representations generated with (solid line) and without (dashed line) matching loss introduced, which suggests that matching loss helped in regularizing the latent space. Quantization step in the VQ-VAE doesn’t significantly interfere with the distance metric.

**Fig. 3 - supplement 1.**
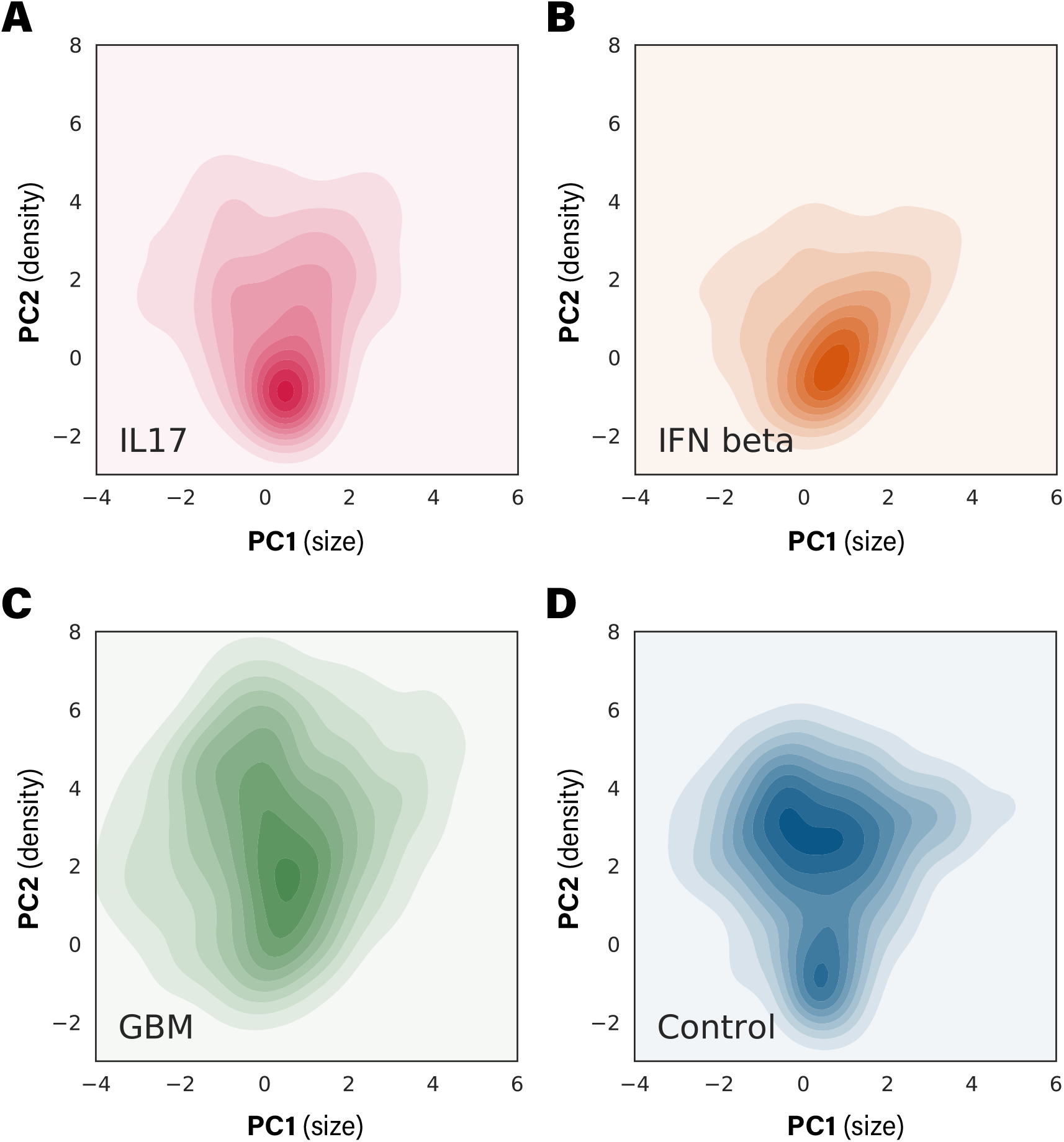
Kernel density estimate (KDE) plots of trajectory-averaged PC1 and PC2 from the test dataset.

**Fig. 3 - supplement 2.**
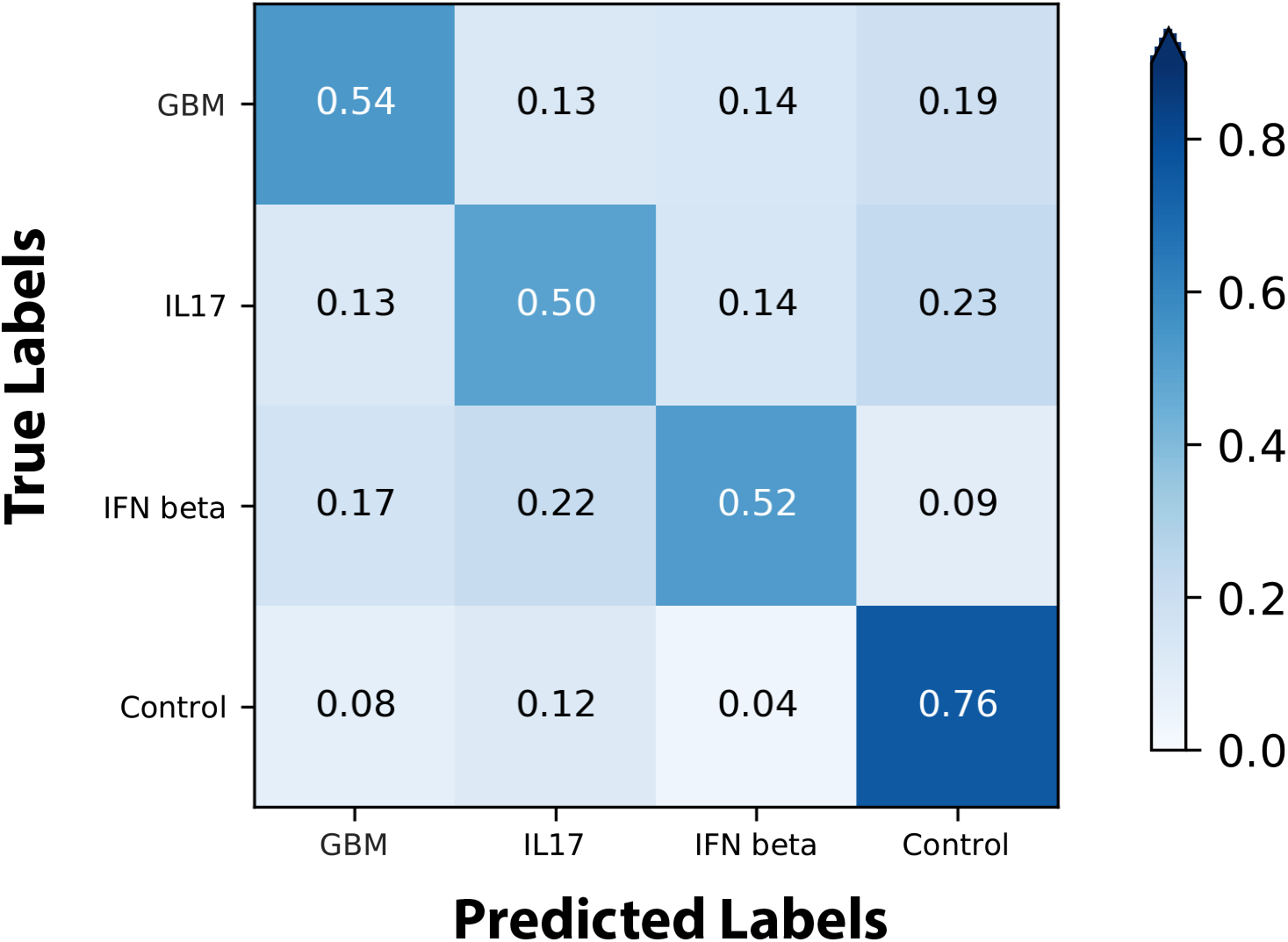
Confusion matrix of classifiers that predict condition/subset from Trajectory Feature Vectors. We employed gradient boosted trees with default hyper-parameters as the predictor models and trained them under 10-fold cross validation.

**Fig. 4 - supplement 1.**
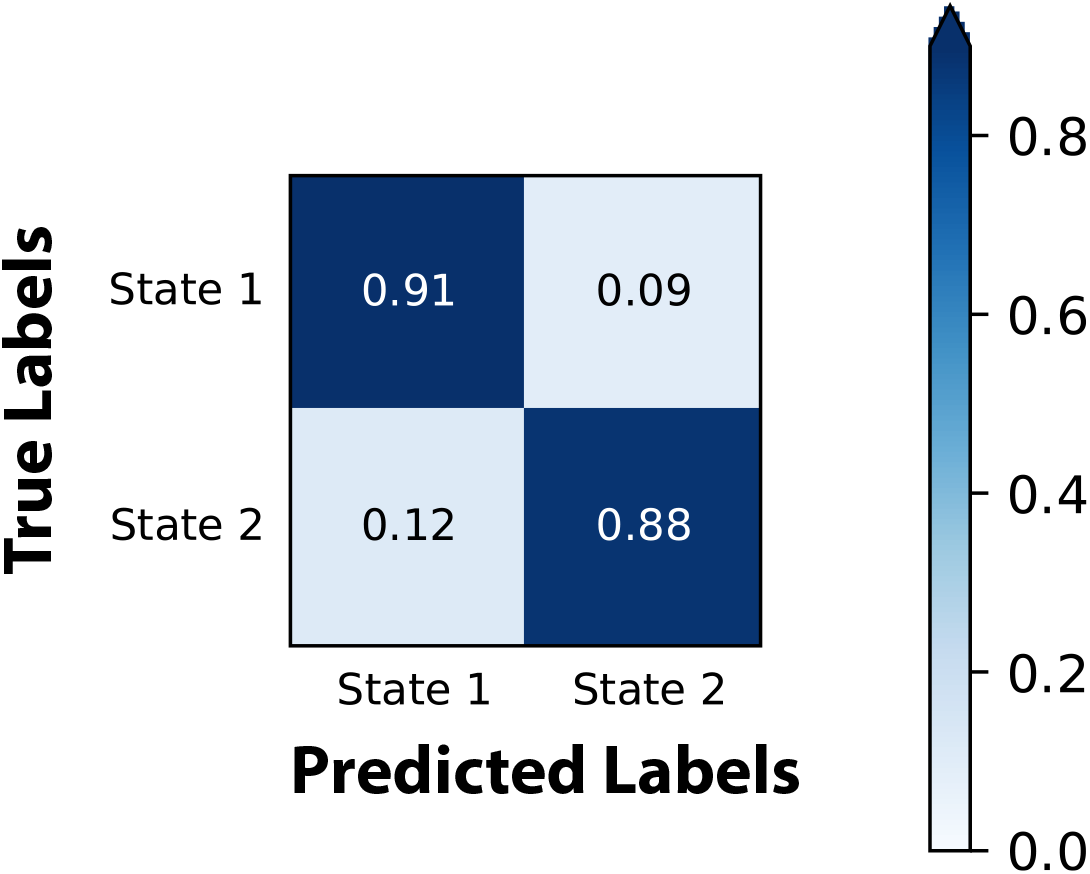
Confusion matrix of classifiers that predict GMM state from Trajectory Feature Vectors. We employed gradient boosted trees with default hyper-parameters as the predictor models and trained them under 10-fold cross validation.

**Fig. 4 - supplement 2.**
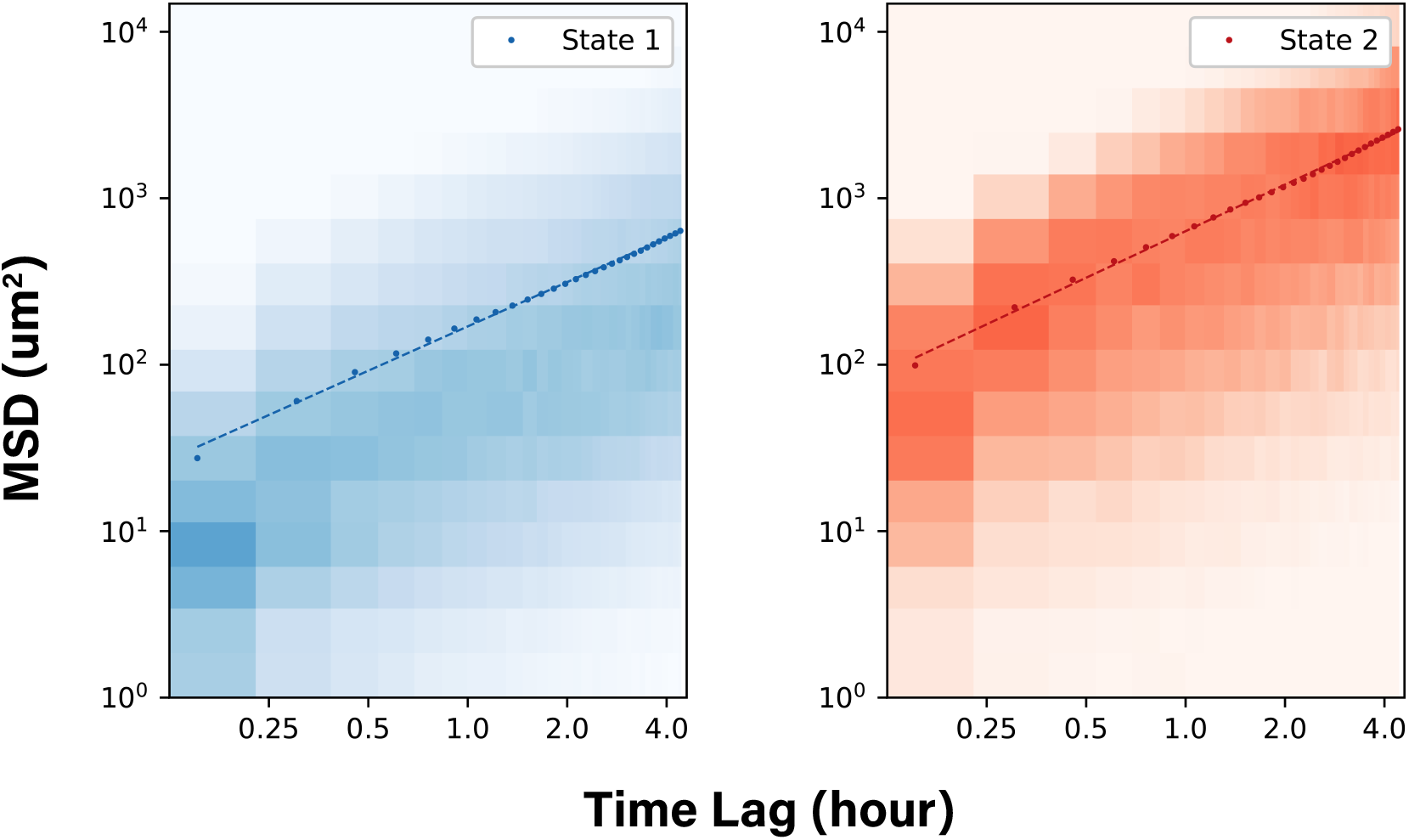
Density map of squared cumulative displacements calculated over trajectories from the two GMM states. Mean square displacements (MSD) are plotted, log-log fit results of the MSD could be found in Figure 4E

**Fig. 4 - supplement 3.**
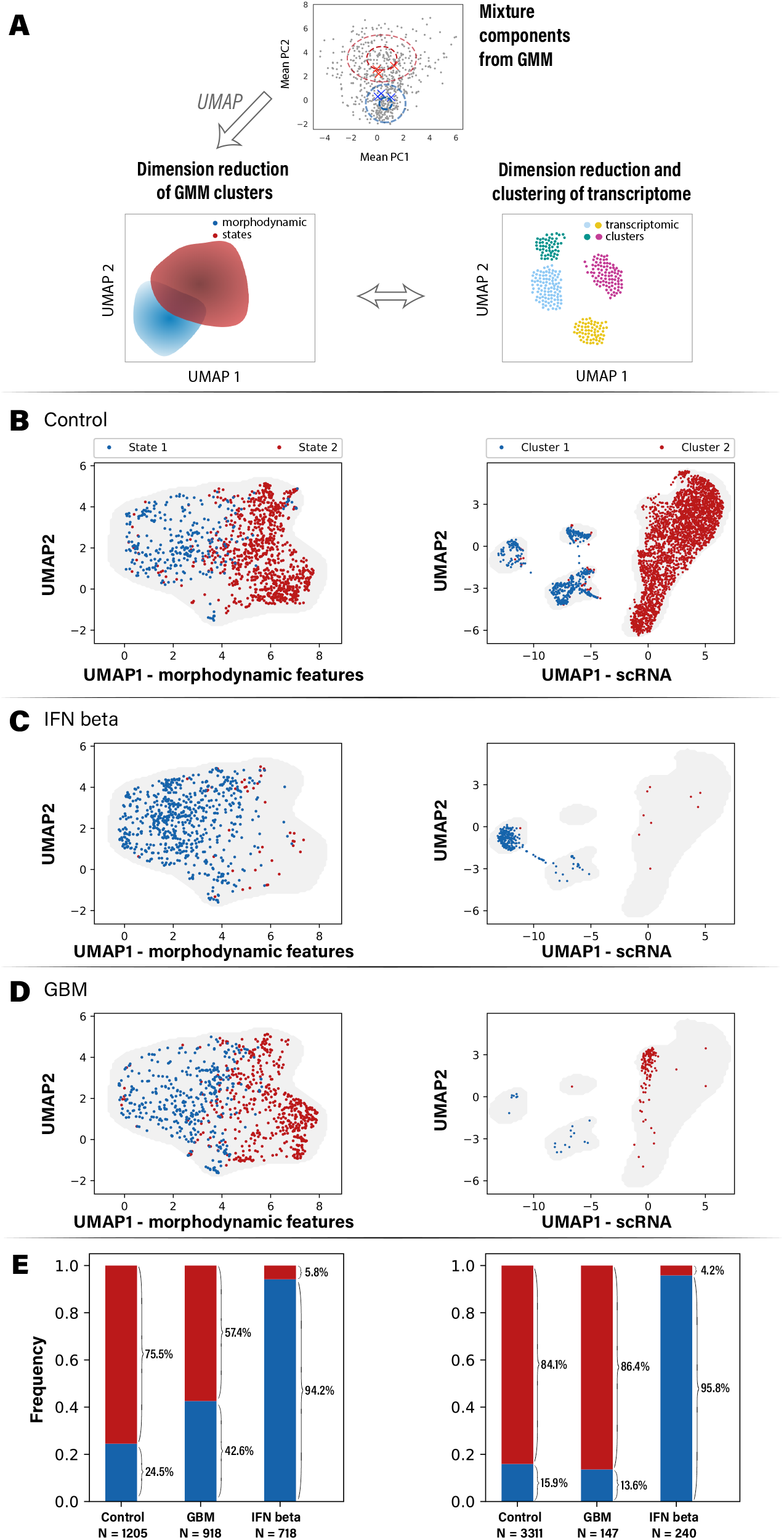
Comparison of morphodynamic states and gene expression per perturbation. **(A)** We compared relative abundance of state-1 and state-2 with clusters of gene expression by computing consistent low dimensional projections (UMAP) of the Trajectory Feature Vectors and transcriptome. Scatter plots of cell populations from three perturbations (**(B)** Control; **(C)** IFN beta; **(D)** GBM) are shown. Each dot represents a trajectory from imaging or a cell measured for transcriptome, and grey shades show the outlines of the distributions. Left panels show the UMAP projections of combination of shape features (first 48 PCs) and motility feature (speed, mean distance) derived from imaging. GMM clustering results are illustrated as the colors of the dots. Right panels show UMAP projections of single-cell RNA sequencing (scRNA-seq) results, on which a similar two-state clustering is performed. **(E)** Distributions of cluster/state assignments of cells from each test condition are plotted as bar graphs. Interestingly, in control and IFN beta conditions, the occupancy ratios between morphodynamic state-1 and state-2 matched with occupancy ratios of gene expression clusters 1 and 2.

**Fig. 4 - supplement 4.**
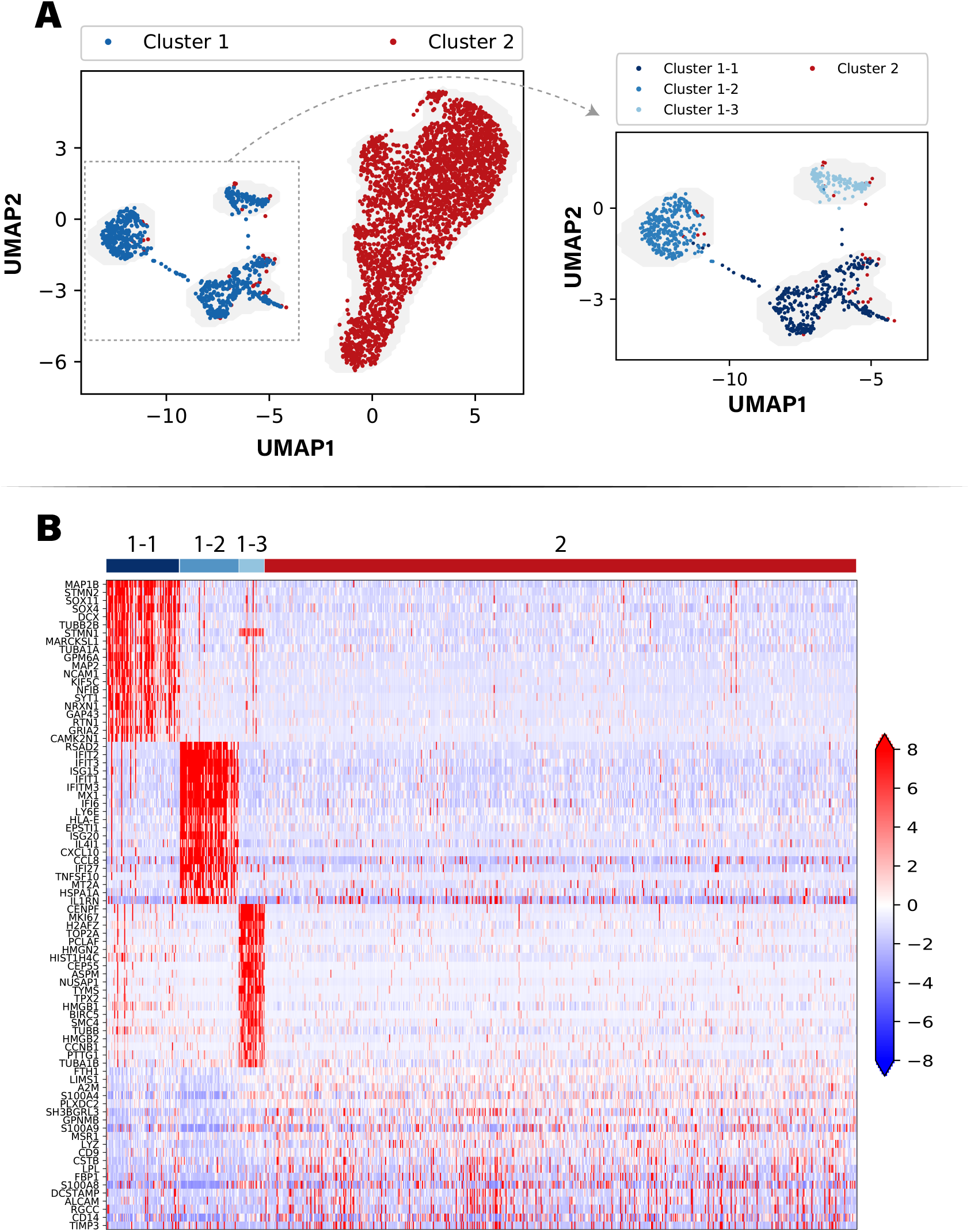
Details of scRNA clustering: **(A)** UMAP projection and clustering of microglia cells based on scRNA-seq. Note that cells from all three conditions are included. Note that cluster 1 is further separated into 3 sub-clusters by applying the same method, in total identifying four molecularly distinct clusters. **(B)** Heatmap of the top differentially expressed genes in each cluster. Cluster 2 includes primarily control microglia and microglia treated with GBM supernatant. Cluster 1-1 is characterized by non-neuronal genes. Cluster 1-2 has high expression of interferon response genes. Cluster 1-3 is defined by high expression of cell cycle genes consistent with a dividing population.

## Supplementary tables

**Supplementary Table 1.**
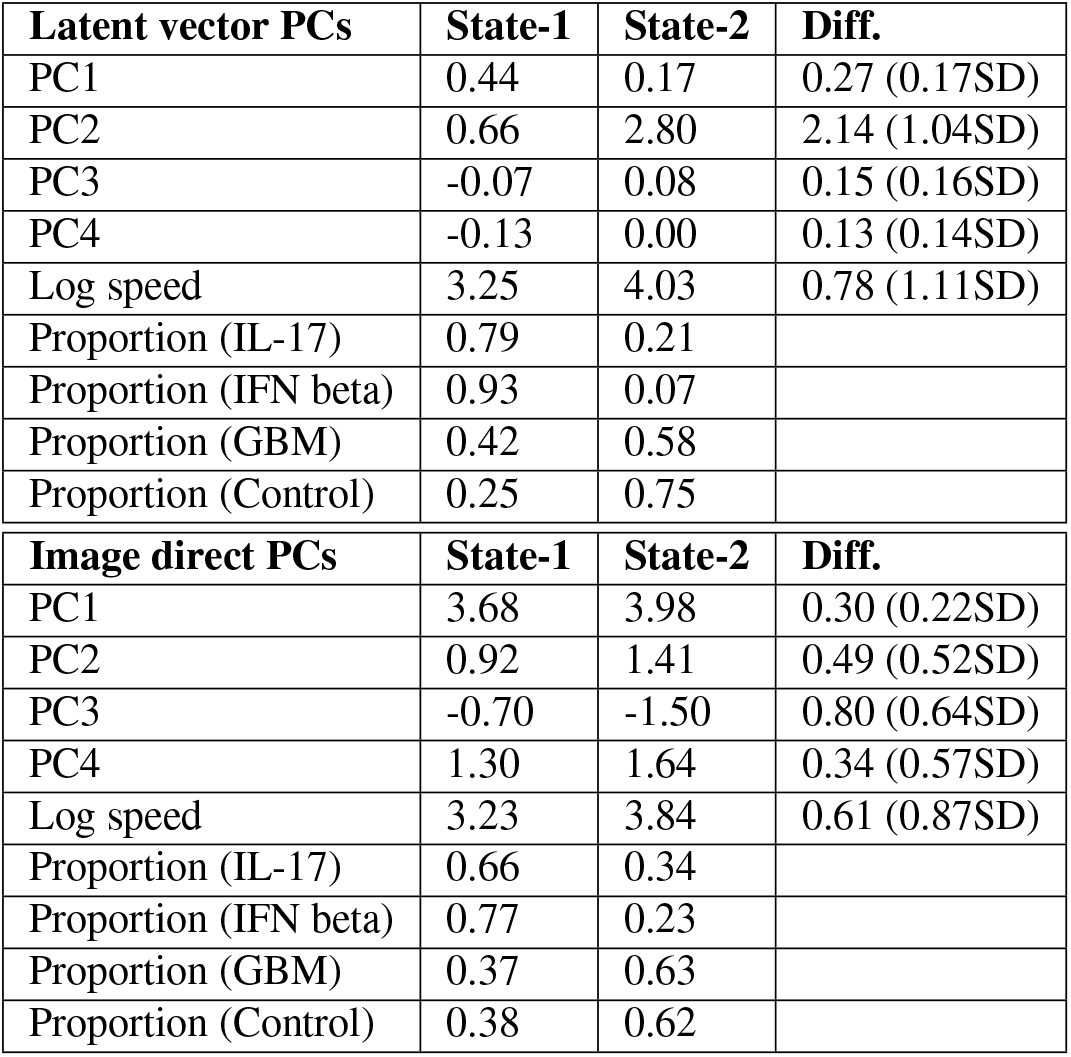
Details of the mixture components detected by GMM using morphodynamic features. “Diff.” column shows the difference in values and standard deviation (SD) between two states. Clusters of latent vector PCs are mainly separated by PC2 and Log speed. Test subsets show different distributions of the two morphodynamic states. Clustering (with the same parameter and procedure) on image direct PCs is less clear, separations between mixture components and test subsets are weaker.

## Supplementary videos

Each video set described below can be downloaded as a zip file from this google drive folder.

**Video Set 1**. 24 hour long videos of phase (density) and retardance (anisotropy) of microglia that illustrate distinct behaviors.

**Video Set 2**. Video clips of representative cell trajectories visualized in Figure 2 and Fig. 2 - supplement 1: we show full fields of view with bounding box and enlarged view of the cell. Note that among the 8 trajectories, 4 are more stable in appearance (corresponding to green and black arrows in Figure 2 and Fig. 2 - supplement 1), 4 undergo significant shape change (orange and brown arrows).

**Video Set 3**. Video clips of representative cell trajectories from each subset of the test dataset: 4 trajectories are provided for each treatment condition (control, GBM, IL17, IFN beta).

**Video Set 4**. Video clips of representative cell trajectories from each of the two morphodynamic state defined by GMM: 5 trajectories are provided for each state.

**Video Set 5**. Video clips of representative cell trajectories that underwent state transitions.

**Video Set 6**. Video clips of sample progenitor cells.

## References

1. Mark-Anthony Bray, Shantanu Singh, Han Han, Chadwick T. Davis, Blake Borgeson, Cathy Hartland, Maria Kost-Alimova, Sigrun M. Gustafsdottir, Christopher C. Gibson, and Anne E. Carpenter, “Cell Painting, a high-content image-based assay for morphological profiling using multiplexed fluorescent dyes,” Nature Protocols, vol. 11, no. 9, pp. 1757–1774, Sept. 2016.

2. Meghan K. Driscoll, Erik S. Welf, Andrew R. Jamieson, Kevin M. Dean, Tadamoto Isogai, Reto Fiolka, and Gaudenz Danuser, “Robust and automated detection of subcellular morphological motifs in 3D microscopy images,” Nature Methods, vol. 16, no. 10, pp. 1037–1044, Oct. 2019.

3. Z. Pincus and J. A. Theriot, “Comparison of quantitative methods for cell-shape analysis,” Journal of Microscopy, vol. 227, no. 2, pp. 140–156, 2007.

4. David A. Van Valen, Takamasa Kudo, Keara M. Lane, Derek N. Macklin, Nicolas T. Quach, Mialy M. DeFelice, Inbal Maayan, Yu Tanouchi, Euan A. Ashley, and Markus W. Covert, “Deep Learning Automates the Quantitative Analysis of Individual Cells in Live-Cell Imaging Experiments,” PLoS computational biology, vol. 12, no. 11, pp. e1005177, Nov. 2016.

5. Assaf Zaritsky, Andrew R. Jamieson, Erik S. Welf, Andres Nevarez, Justin Cillay, Ugur Eskiocak, Brandi L. Cantarel, and Gaudenz Danuser, “Inter-pretable deep learning of label-free live cell images uncovers functional hallmarks of highly-metastatic melanoma,” bioRxiv, p. 2020.05.15.096628, May 2020.

6. Caleb K. Chan, Amalia Hadjitheodorou, Tony Y.-C. Tsai, and Julie A. Theriot, “Quantitative comparison of principal component analysis and unsupervised deep learning using variational autoencoders for shape analysis of motile cells,” bioRxiv, p. 2020.06.26.174474, June 2020.

7. Kinneret Keren, Zachary Pincus, Greg M. Allen, Erin L. Barnhart, Gerard Marriott, Alex Mogilner, and Julie A. Theriot, “Mechanism of shape determination in motile cells,” Nature, vol. 453, no. 7194, pp. 475–480, May 2008.

8. Wallace F. Marshall, “Origins of cellular geometry,” BMC Biology, vol. 9, no. 1, pp. 57, Aug. 2011.

9. Shahriar Shadkhoo and Madhav Mani, “The Role of Cytoplasmic Interactions in the Collective Polarization of Tissues and its Interplay with Cellular Geometry,” bioRxiv, p. 289520, Mar. 2018.

10. P. Rezaie, G. Trillo-Pazos, J. Greenwood, I. P. Everall, and D. K. Male, “Motility and Ramification of Human Fetal Microglia in Culture: An Investigation Using Time-Lapse Video Microscopy and Image Analysis,” Experimental Cell Research, vol. 274, no. 1, pp. 68–82, Mar. 2002.

11. Assaf Zaritsky, Erik S. Welf, Yun-Yu Tseng, M. Angeles Rabadán, Xavier Serra-Picamal, Xavier Trepat, and Gaudenz Danuser, “Seeds of Locally Aligned Motion and Stress Coordinate a Collective Cell Migration,” Biophysical Journal, vol. 109, no. 12, pp. 2492–2500, Dec. 2015.

12. Erick Moen, Enrico Borba, Geneva Miller, Morgan Schwartz, Dylan Bannon, Nora Koe, Isabella Camplisson, Daniel Kyme, Cole Pavelchek, Tyler Price, Takamasa Kudo, Edward Pao, William Graf, and David Van Valen, “Accurate cell tracking and lineage construction in live-cell imaging experiments with deep learning,” bioRxiv, p. 803205, Oct. 2019.

13. Beate Neumann, Thomas Walter, Jean-Karim Hériché, Jutta Bulkescher, Holger Erfle, Christian Conrad, Phill Rogers, Ina Poser, Michael Held, Urban Liebel, Cihan Cetin, Frank Sieckmann, Gregoire Pau, Rolf Kabbe, Annelie Wünsche, Venkata Satagopam, Michael H. A. Schmitz, Catherine Chapuis, Daniel W. Gerlich, Reinhard Schneider, Roland Eils, Wolfgang Huber, Jan-Michael Peters, Anthony A. Hyman, Richard Durbin, Rainer Pepperkok, and Jan Ellenberg, “Phenotypic profiling of the human genome by time-lapse microscopy reveals cell division genes,” Nature, vol. 464, no. 7289, pp. 721–727, Apr. 2010.

14. Chawin Ounkomol, Sharmishtaa Seshamani, Mary M. Maleckar, Forrest Collman, and Gregory R. Johnson, “Label-free prediction of three-dimensional fluorescence images from transmitted-light microscopy,” Nature Methods, vol. 15, no. 11, pp. 917, Nov. 2018.

15. Eric M. Christiansen, Samuel J. Yang, D. Michael Ando, Ashkan Javaherian, Gaia Skibinski, Scott Lipnick, Elliot Mount, Alison O’Neil, Kevan Shah, Alicia K. Lee, Piyush Goyal, William Fedus, Ryan Poplin, Andre Esteva, Marc Berndl, Lee L. Rubin, Philip Nelson, and Steven Finkbeiner, “In silico labeling: Predicting fluorescent labels in unlabeled images,” Cell, vol. 173, no. 3, pp. 792–803.e19, Apr. 2018.

16. Syuan-Ming Guo, Li-Hao Yeh, Jenny Folkesson, Ivan E Ivanov, Anitha P Krishnan, Matthew G Keefe, Ezzat Hashemi, David Shin, Bryant B Chhun, Nathan H Cho, Manuel D Leonetti, May H Han, Tomasz Nowakowski, and Shalin B Mehta, “Revealing architectural order with quantitative label-free imaging and deep learning,” eLife, vol. 9, pp. e55502, July 2020, Publisher: eLife Sciences Publications, Ltd.

17. Bryan He, Ludvig Bergenstråhle, Linnea Stenbeck, Abubakar Abid, Alma Andersson, Åke Borg, Jonas Maaskola, Joakim Lundeberg, and James Zou, “Integrating spatial gene expression and breast tumour morphology via deep learning,” Nature Biomedical Engineering, pp. 1–8, June 2020.

18. Jacob C. Kimmel, Amy Y. Chang, Andrew S. Brack, and Wallace F. Marshall, “Inferring cell state by quantitative motility analysis reveals a dynamic state system and broken detailed balance,” PLOS Computational Biology, vol. 14, no. 1, pp. e1005927, Jan. 2018.

19. Hirofumi Kobayashi, Keith C. Cheveralls, Manuel D. Leonetti, and Loic A. Royer, “Self-Supervised Deep-Learning Encodes High-Resolution Features of Protein Subcellular Localization,” bioRxiv, p. 2021.03.29.437595, Mar. 2021, Publisher: Cold Spring Harbor Laboratory Section: New Results.

20. Anne E. Carpenter and David M. Sabatini, “Systematic genome-wide screens of gene function,” Nature Reviews Genetics, vol. 5, no. 1, pp. 11–22, Jan. 2004.

21. Keara Lane, David Van Valen, Mialy M. DeFelice, Derek N. Macklin, Takamasa Kudo, Ariel Jaimovich, Ambrose Carr, Tobias Meyer, Dana Pe’er, Stéphane C. Boutet, and Markus W. Covert, “Measuring Signaling and RNA-Seq in the Same Cell Links Gene Expression to Dynamic Patterns of NF-ŸB Activation,” Cell Systems, vol. 4, no. 4, pp. 458–469.e5, Apr. 2017.

22. Karren Dai Yang, Anastasiya Belyaeva, Saradha Venkatachalapathy, Karthik Damodaran, Adityanarayanan Radhakrishnan, Abigail Katcoff, G. V. Shivashankar, and Caroline Uhler, “Multi-Domain Translation between Single-Cell Imaging and Sequencing Data using Autoencoders,” bioRxiv, p. 2019.12.13.875922, Dec. 2019.

23. Kaytlyn A. Gerbin, Tanya Grancharova, Rory Donovan-Maiye, Melissa C. Hendershott, Jackson Brown, Stephanie Q. Dinh, Jamie L. Gehring, Matthew Hirano, Gregory R. Johnson, Aditya Nath, Angelique Nelson, Charles M. Roco, Alexander B. Rosenberg, M. Filip Sluzewski, Matheus P. Viana, Calysta Yan, Rebecca J. Zaunbrecher, Kimberly R. Cordes Metzler, Vilas Menon, Sean P. Palecek, Georg Seelig, Nathalie Gaudreault, Theo Knijnenburg, Susanne M. Rafelski, Julie A. Theriot, and Ruwanthi N. Gunawardane, “Cell states beyond transcriptomics: Integrating structural organization and gene expression in hiPSC-derived cardiomyocytes,” bioRxiv, p. 2020.05.26.081083, May 2020.

24. Michael W Salter and Beth Stevens, “Microglia emerge as central players in brain disease,” Nature medicine, vol. 23, no. 9, pp. 1018, 2017.

25. Chintan Chhatbar, Claudia N Detje, Elena Grabski, Katharina Borst, Julia Spanier, Luca Ghita, David A Elliott, Marta Joana Costa Jordao, Nora Mueller, James Sutton, et al., “Type i interferon receptor signaling of neurons and astrocytes regulates microglia activation during viral encephalitis,” Cell reports, vol. 25, no. 1, pp. 118–129, 2018.

26. Starlee Lively and Lyanne C Schlichter, “Microglia responses to pro-inflammatory stimuli (lps, ifn”+ tnf–) and reprogramming by resolving cytokines (il-4, il-10),” Frontiers in Cellular Neuroscience, vol. 12, pp. 215, 2018.

27. Axel Nimmerjahn, Frank Kirchhoff, and Fritjof Helmchen, “Resting microglial cells are highly dynamic surveillants of brain parenchyma in vivo,” Science, vol. 308, no. 5726, pp. 1314–1318, 2005.

28. Louis-Philippe Bernier, Christopher J Bohlen, Elisa M York, Hyun B Choi, Alireza Kamyabi, Lasse Dissing-Olesen, Jasmin K Hefendehl, Hannah Y Collins, Beth Stevens, Ben A Barres, et al., “Nanoscale surveillance of the brain by microglia via camp-regulated filopodia,” Cell reports, vol. 27, no. 10, pp. 2895–2908, 2019.

29. Stephen C Noctor, Elisa Penna, Hunter Shepherd, Christian Chelson, Nicole Barger, Verónica Martínez-Cerdeño, and Alice F Tarantal, “Periven-tricular microglial cells interact with dividing precursor cells in the nonhuman primate and rodent prenatal cerebral cortex,” Journal of Comparative Neurology, vol. 527, no. 10, pp. 1598–1609, 2019.

30. David V Hansen, Jan H Lui, Philip RL Parker, and Arnold R Kriegstein, “Neurogenic radial glia in the outer subventricular zone of human neocortex,” Nature, vol. 464, no. 7288, pp. 554–561, 2010.

31. Aaron van den Oord and Oriol Vinyals, “Neural discrete representation learning,” in Advances in Neural Information Processing Systems, 2017, pp. 6306–6315.

32. Ali Razavi, Aaron van den Oord, and Oriol Vinyals, “Generating Diverse High-Fidelity Images with VQ-VAE-2,” in Advances in Neural Information Processing Systems 32, H. Wallach, H. Larochelle, A. Beygelzimer, F. d\textquotesingle Alché-Buc, E. Fox, and R. Garnett, Eds., pp. 14866–14876. Curran Associates, Inc., 2019.

33. Will Zou, Shenghuo Zhu, Kai Yu, and Andrew Y Ng, “Deep learning of invariant features via simulated fixations in video,” in Advances in neural information processing systems, 2012, pp. 3203–3211.

34. Florian Schroff, Dmitry Kalenichenko, and James Philbin, “Facenet: A unified embedding for face recognition and clustering,” arXiv preprint 1503.03832, 2015.

35. Leland McInnes, John Healy, and James Melville, “Umap: Uniform manifold approximation and projection for dimension reduction,” arXiv preprint 1802.03426, 2018.

36. Anyong Yu, Haizhen Duan, Tianxi Zhang, Yong Pan, Zhi Kou, Xiaojun Zhang, Yuanlan Lu, Song Wang, and Zhao Yang, “Il-17a promotes microglial activation and neuroinflammation in mouse models of intracerebral haemorrhage,” Molecular immunology, vol. 73, pp. 151–157, 2016.

37. Jun Kawanokuchi, Kouki Shimizu, Atsumi Nitta, Kiyofumi Yamada, Tetsuya Mizuno, Hideyuki Takeuchi, and Akio Suzumura, “Production and functions of il-17 in microglia,” Journal of neuroimmunology, vol. 194, no. 1-2, pp. 54–61, 2008.

38. Gloria B Choi, Yeong S Yim, Helen Wong, Sangdoo Kim, Hyunju Kim, Sangwon V Kim, Charles A Hoeffer, Dan R Littman, and Jun R Huh, “The maternal interleukin-17a pathway in mice promotes autism-like phenotypes in offspring,” Science, vol. 351, no. 6276, pp. 933–939, 2016.

39. Thomas Blank and Marco Prinz, “Type i interferon pathway in cns homeostasis and neurological disorders,” Glia, vol. 65, no. 9, pp. 1397–1406, 2017.

40. Giuseppa Mudò, Monica Frinchi, Domenico Nuzzo, Pietro Scaduto, Fulvio Plescia, Maria F Massenti, Marta Di Carlo, Carla Cannizzaro, Giovanni Cassata, Luca Cicero, et al., “Anti-inflammatory and cognitive effects of interferon-—1a (ifn—1a) in a rat model of alzheimer’s disease,” Journal of neuroinflammation, vol. 16, no. 1, pp. 1–16, 2019.

41. David Gosselin, Dylan Skola, Nicole G Coufal, Inge R Holtman, Johannes CM Schlachetzki, Eniko Sajti, Baptiste N Jaeger, Carolyn O’Connor, Conor Fitzpatrick, Martina P Pasillas, et al., “An environment-dependent transcriptional network specifies human microglia identity,” Science, vol. 356, no. 6344, 2017.

42. Jerome H Friedman, “Greedy function approximation: a gradient boosting machine,” Annals of statistics, pp. 1189–1232, 2001.

43. Olaf Ronneberger, Philipp Fischer, and Thomas Brox, “U-net: Convolutional networks for biomedical image segmentation,” in International Conference on Medical image computing and computer-assisted intervention. Springer, 2015, pp. 234–241.

44. Stuart Berg, Dominik Kutra, Thorben Kroeger, Christoph N. Straehle, Bernhard X. Kausler, Carsten Haubold, Martin Schiegg, Janez Ales, Thorsten Beier, Markus Rudy, Kemal Eren, Jaime I. Cervantes, Buote Xu, Fynn Beuttenmueller, Adrian Wolny, Chong Zhang, Ullrich Koethe, Fred A. Hamprecht, and Anna Kreshuk, “Ilastik: Interactive machine learning for (bio)image analysis,” Nature Methods, vol. 16, no. 12, pp. 1226–1232, Dec. 2019.

45. Kaiming He, Xiangyu Zhang, Shaoqing Ren, and Jian Sun, “Deep residual learning for image recognition,” in Proceedings of the IEEE conference on computer vision and pattern recognition, 2016, pp. 770–778.

46. Diederik P Kingma and Jimmy Ba, “Adam: A method for stochastic optimization,” arXiv preprint 1412.6980, 2014.

47. Martin Ester, Hans-Peter Kriegel, Jörg Sander, and Xiaowei Xu, “A density-based algorithm for discovering clusters in large spatial databases with noise.,” in Knowledge Discovery and Data Mining, 1996, vol. 96, pp. 226–231.

48. Joseph Redmon, Santosh Divvala, Ross Girshick, and Ali Farhadi, “You only look once: Unified, real-time object detection,” in Proceedings of the IEEE conference on computer vision and pattern recognition, 2016, pp. 779–788.

49. Kaiming He, Georgia Gkioxari, Piotr Dollár, and Ross Girshick, “Mask r-cnn,” in Proceedings of the IEEE international conference on computer vision, 2017, pp. 2961–2969.

50. Khuloud Jaqaman, Dinah Loerke, Marcel Mettlen, Hirotaka Kuwata, Sergio Grinstein, Sandra L. Schmid, and Gaudenz Danuser, “Robust single-particle tracking in live-cell time-lapse sequences,” Nature Methods, vol. 5, no. 8, pp. 695–702, Aug. 2008.

51. Adam Paszke, Sam Gross, Soumith Chintala, Gregory Chanan, Edward Yang, Zachary DeVito, Zeming Lin, Alban Desmaison, Luca Antiga, and Adam Lerer, “Automatic differentiation in pytorch,” in NIPS-W, 2017.

52. Laurens van der Maaten and Geoffrey Hinton, “Visualizing data using t-sne,” Journal of machine learning research, vol. 9, no. Nov, pp. 2579–2605, 2008.

53. Christopher S McGinnis, David M Patterson, Juliane Winkler, Daniel N Conrad, Marco Y Hein, Vasudha Srivastava, Jennifer L Hu, Lyndsay M Murrow, Jonathan S Weissman, Zena Werb, et al., “Multi-seq: sample multiplexing for single-cell rna sequencing using lipid-tagged indices,” Nature methods, vol. 16, no. 7, pp. 619, 2019.

54. Christoph Hafemeister and Rahul Satija, “Normalization and variance stabilization of single-cell rna-seq data using regularized negative binomial regression,” Genome biology, vol. 20, no. 1, pp. 1–15, 2019.

55. Vincent A Traag, Ludo Waltman, and Nees Jan van Eck, “From louvain to leiden: guaranteeing well-connected communities,” Scientific reports, vol. 9, no. 1, pp. 1–12, 2019.

56. Diederik P. Kingma and Max Welling, “Auto-encoding variational bayes,” arXiv preprint 1312.6114, 2013.

57. Alireza Makhzani, Jonathon Shlens, Navdeep Jaitly, Ian Goodfellow, and Brendan Frey, “Adversarial autoencoders,” arXiv preprint 1511.05644, 2015.

