## Supplementary figures and images for "DynaMorph: self-supervised learning of morphodynamic states of live cells"

### sample_B4-Site_0_63.gif

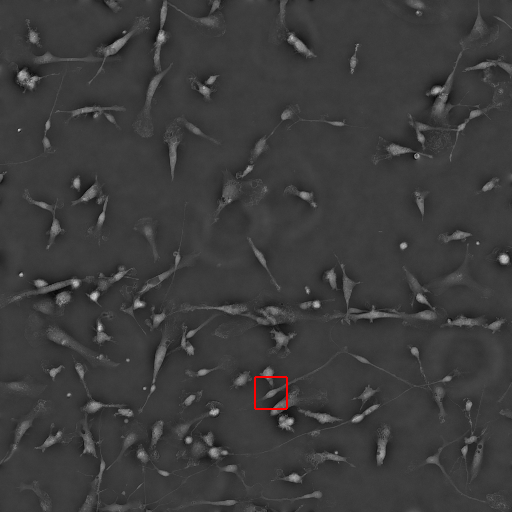

### sample_B4-Site_3_50.gif

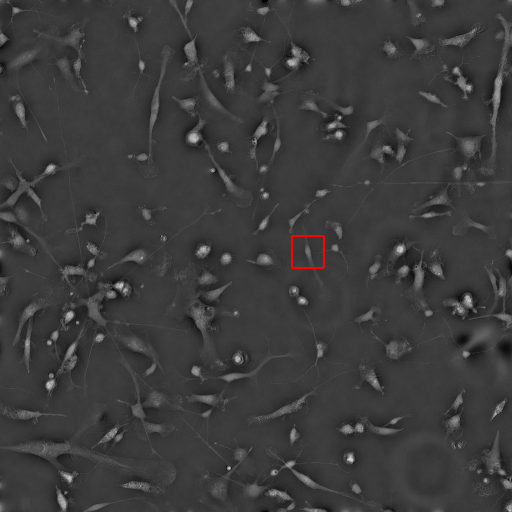

### supp_video_Control_sample_bbox_Control-Site_1_441.gif

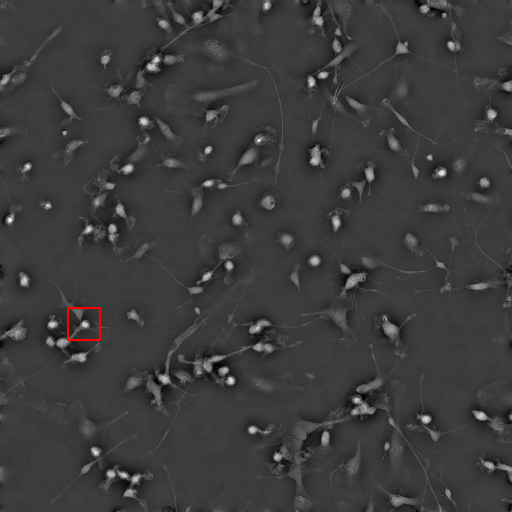

### supp_video_Control_sample_bbox_Control-Site_2_202.gif

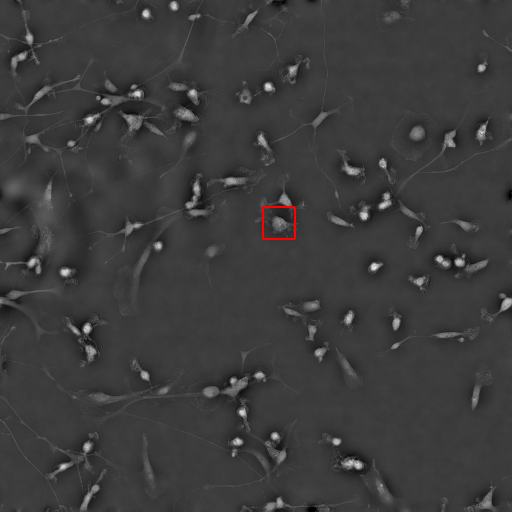

### supp_video_Control_sample_bbox_Control-Site_3_253.gif

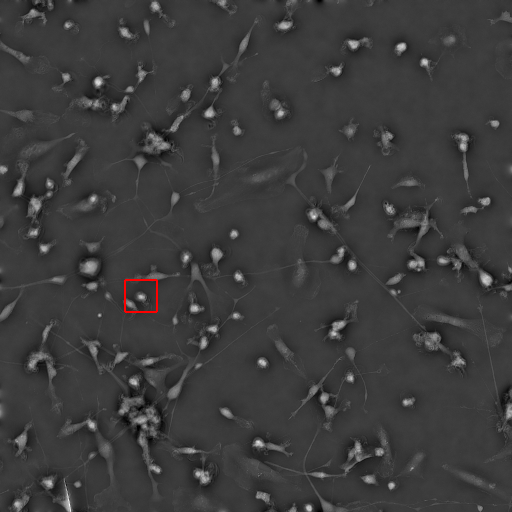

### supp_video_Control_sample_bbox_Control-Site_3_349.gif

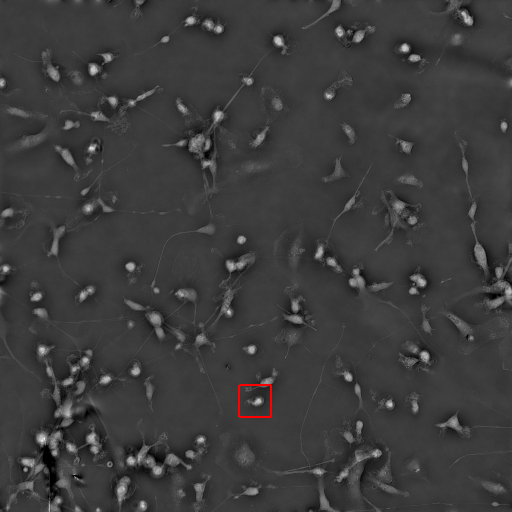

### supp_video_Control_sample_movie_Control-Site_1_441.gif

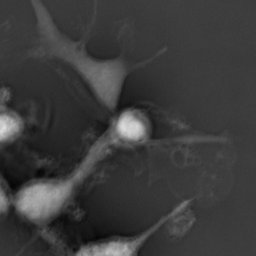

### supp_video_Control_sample_movie_Control-Site_2_202.gif

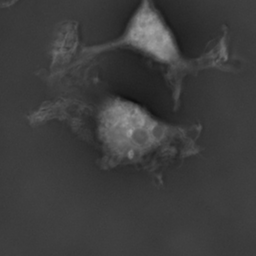

### supp_video_Control_sample_movie_Control-Site_3_253.gif

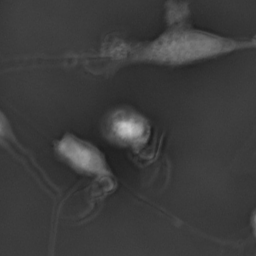

### supp_video_Control_sample_movie_Control-Site_3_349.gif

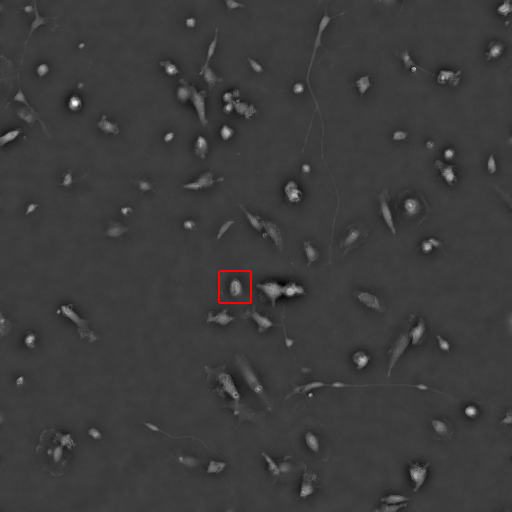

### supp_video_GBM_sample_bbox_GBM-Site_0_214.gif

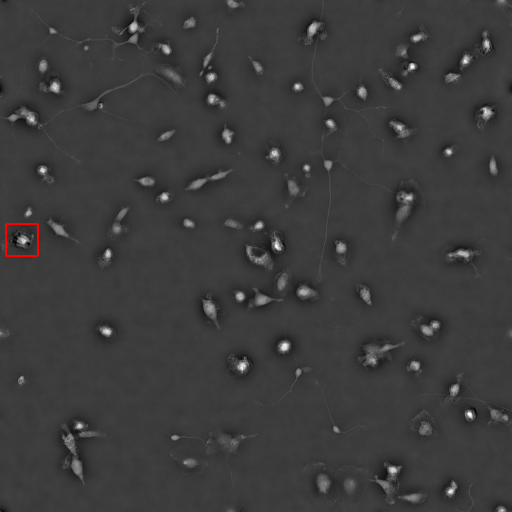

### supp_video_GBM_sample_bbox_GBM-Site_1_84.gif

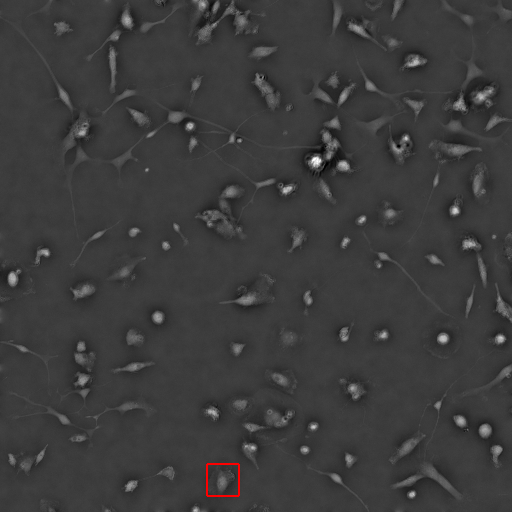

### supp_video_GBM_sample_bbox_GBM-Site_2_39.gif

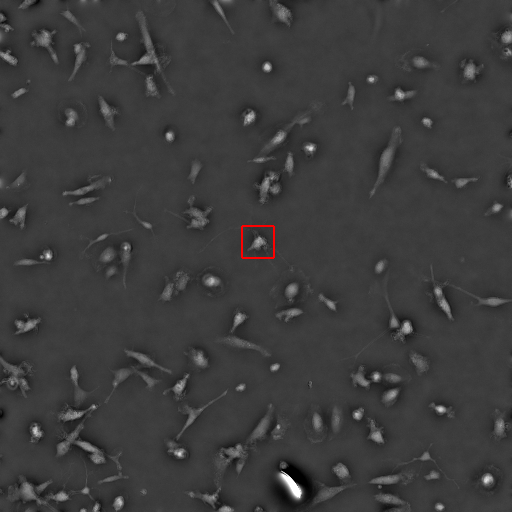

### supp_video_GBM_sample_movie_GBM-Site_0_43.gif

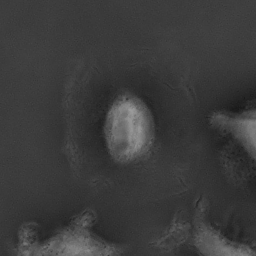

### supp_video_GBM_sample_movie_GBM-Site_0_214.gif

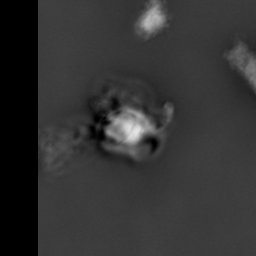

### supp_video_GBM_sample_movie_GBM-Site_1_84.gif

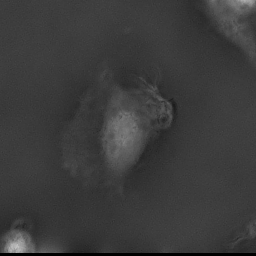

### supp_video_GBM_sample_movie_GBM-Site_2_39.gif

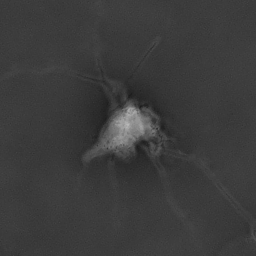

### supp_video_IFNbeta_sample_bbox_IFNbeta-Site_1_252.gif

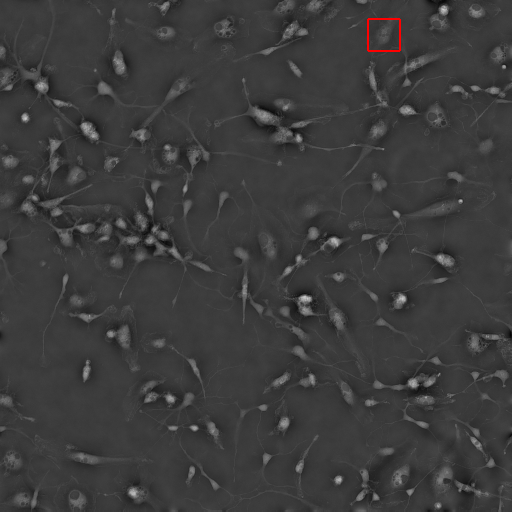

### supp_video_IFNbeta_sample_bbox_IFNbeta-Site_2_243.gif

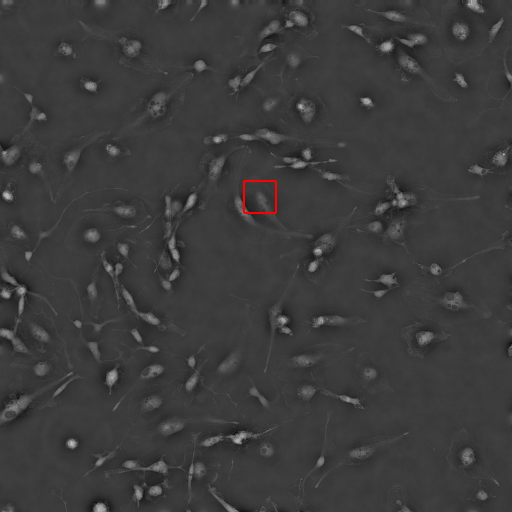

### supp_video_IFNbeta_sample_bbox_IFNbeta-Site_6_72.gif

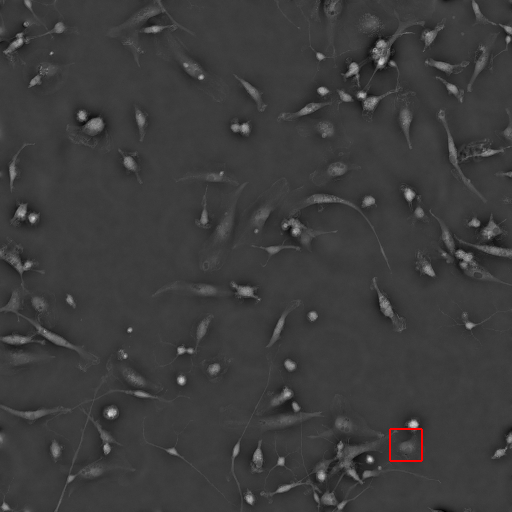

### supp_video_IFNbeta_sample_bbox_IFNbeta-Site_7_28.gif

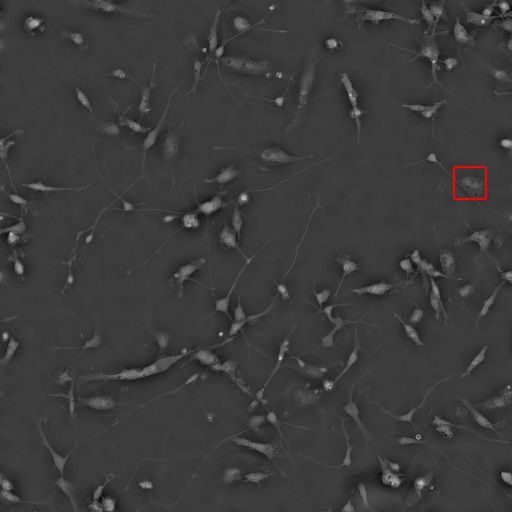

### supp_video_IFNbeta_sample_movie_IFNbeta-Site_1_252.gif

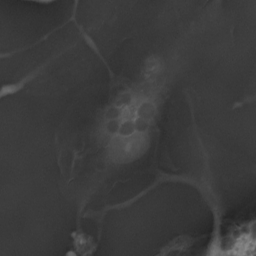

### supp_video_IFNbeta_sample_movie_IFNbeta-Site_2_243.gif

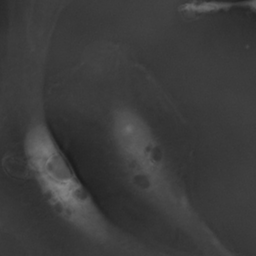

### supp_video_IFNbeta_sample_movie_IFNbeta-Site_6_72.gif

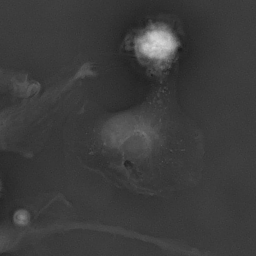

### supp_video_IFNbeta_sample_movie_IFNbeta-Site_7_28.gif

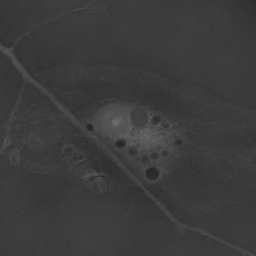

### supp_video_IL17_sample_bbox_IL17-Site_0_89.gif

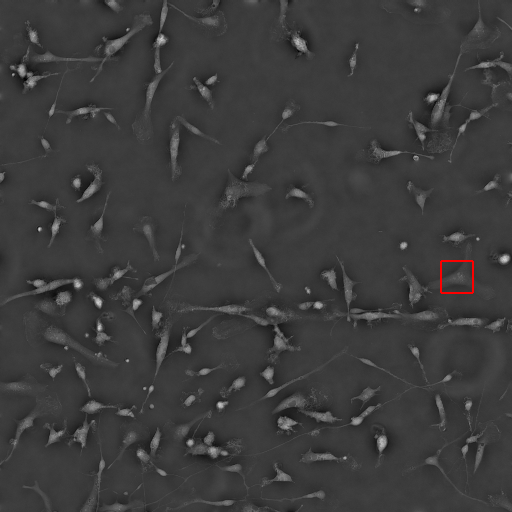

### supp_video_IL17_sample_bbox_IL17-Site_2_51.gif

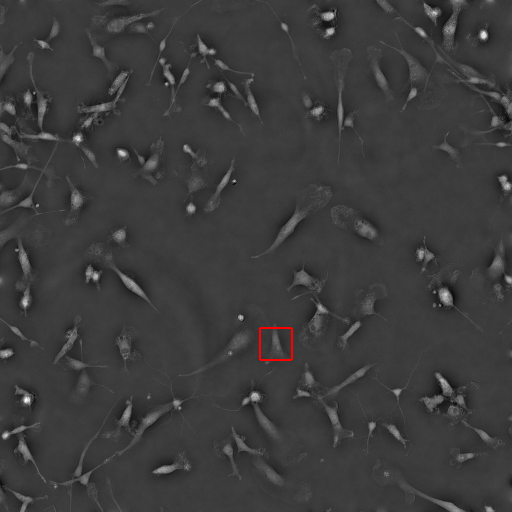

### supp_video_IL17_sample_bbox_IL17-Site_2_282.gif

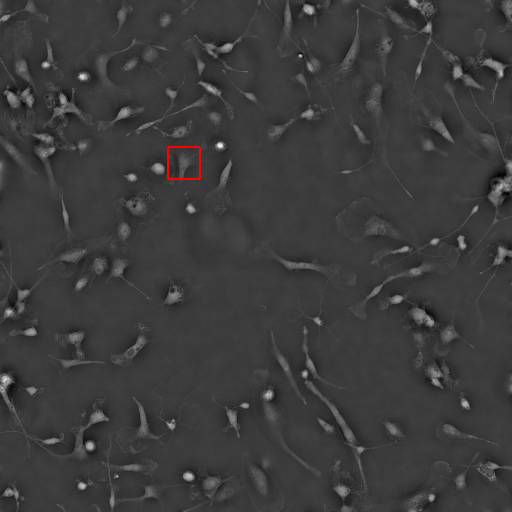

### supp_video_IL17_sample_bbox_IL17-Site_3_30.gif

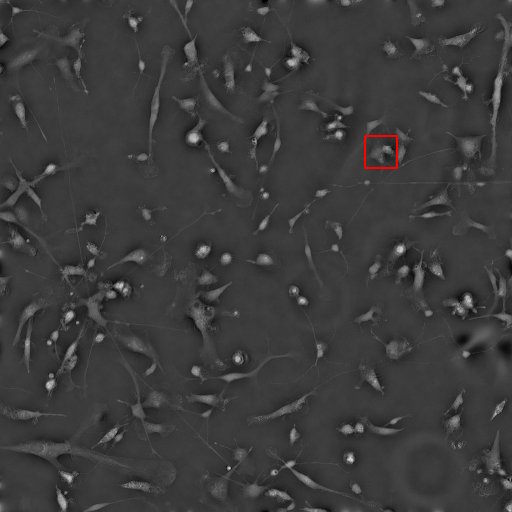

### supp_video_IL17_sample_movie_IL17-Site_0_89.gif

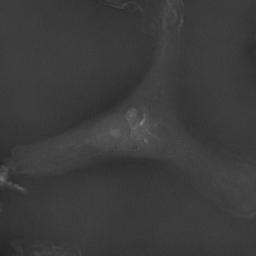

### supp_video_IL17_sample_movie_IL17-Site_2_51.gif

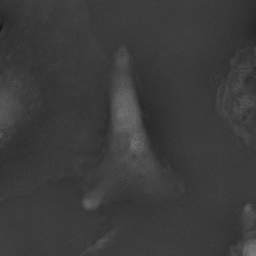

### supp_video_IL17_sample_movie_IL17-Site_2_282.gif

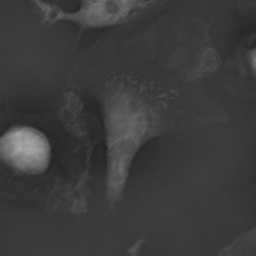

### supp_video_IL17_sample_movie_IL17-Site_3_30.gif

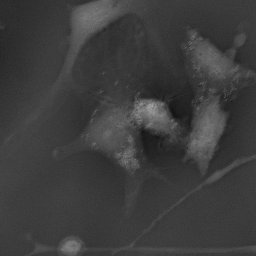

### supp_video_PCA_test_sample_bbox_black.gif

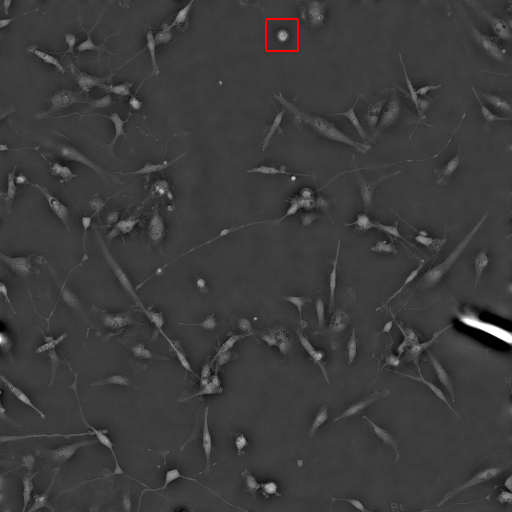

### supp_video_PCA_test_sample_bbox_brown.gif

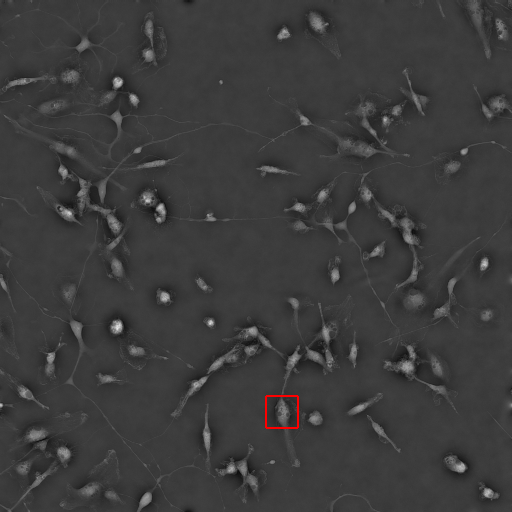

### supp_video_PCA_test_sample_bbox_green.gif

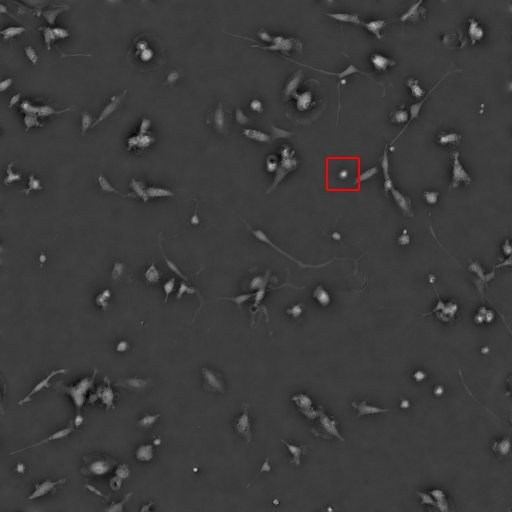

### supp_video_PCA_test_sample_bbox_orange.gif

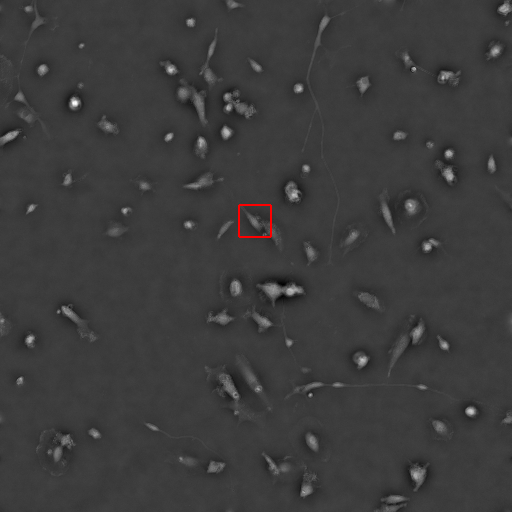

### supp_video_PCA_test_sample_movie_black.gif

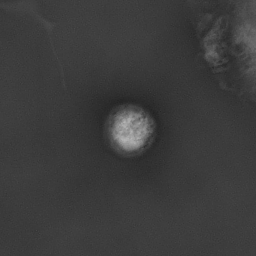

### supp_video_PCA_test_sample_movie_brown.gif

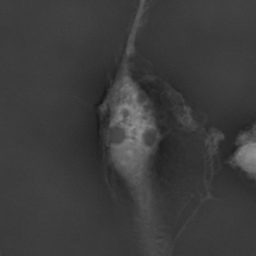

### supp_video_PCA_test_sample_movie_green.gif

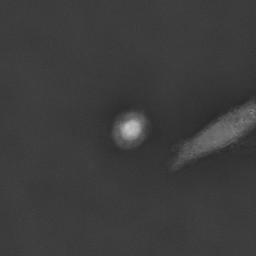

### supp_video_PCA_test_sample_movie_orange.gif

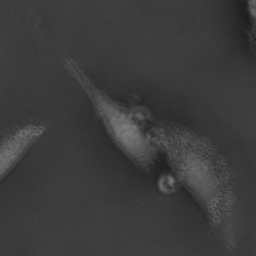

### supp_video_PCA_train_sample_bbox_black.gif

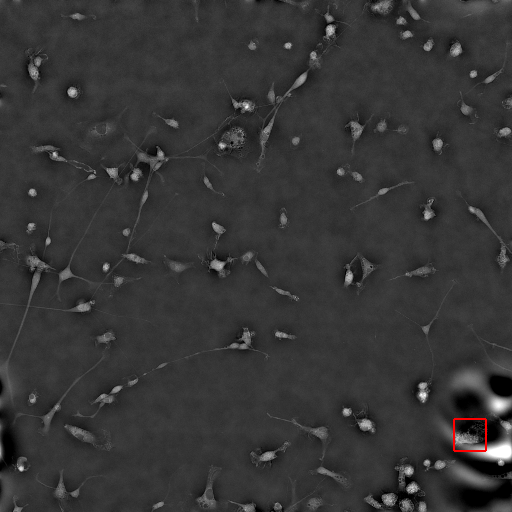

### supp_video_PCA_train_sample_bbox_brown.gif

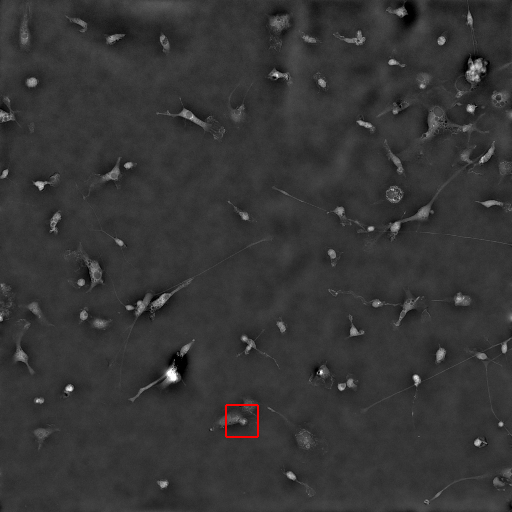

### supp_video_PCA_train_sample_bbox_green.gif

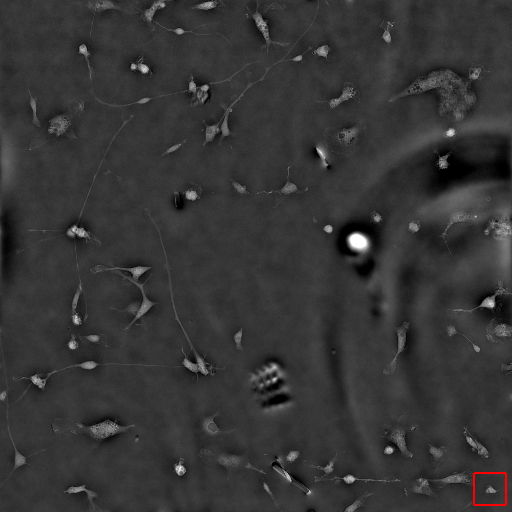

### supp_video_PCA_train_sample_bbox_orange.gif

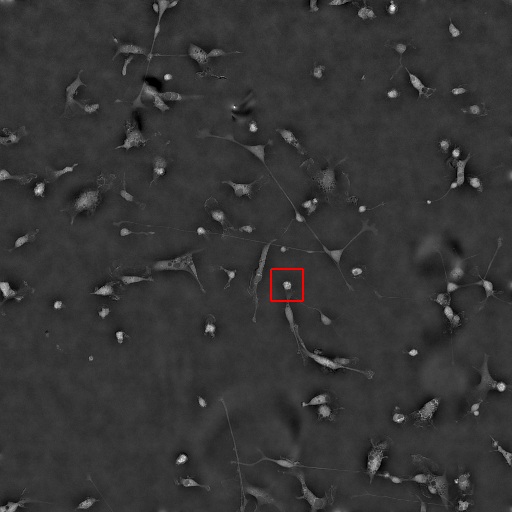

### supp_video_PCA_train_sample_movie_black.gif

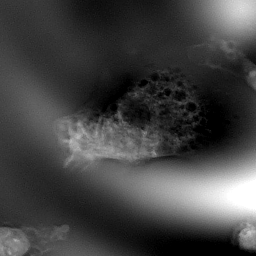

### supp_video_PCA_train_sample_movie_brown.gif

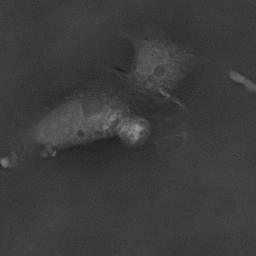

### supp_video_PCA_train_sample_movie_green.gif

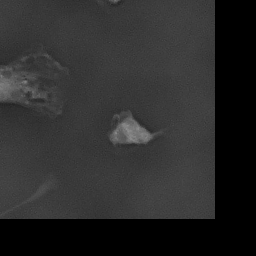

### supp_video_PCA_train_sample_movie_orange.gif

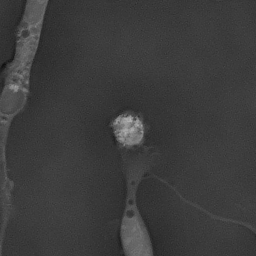

### supp_video_state_1_sample_bbox_Control-Site_2_253.gif

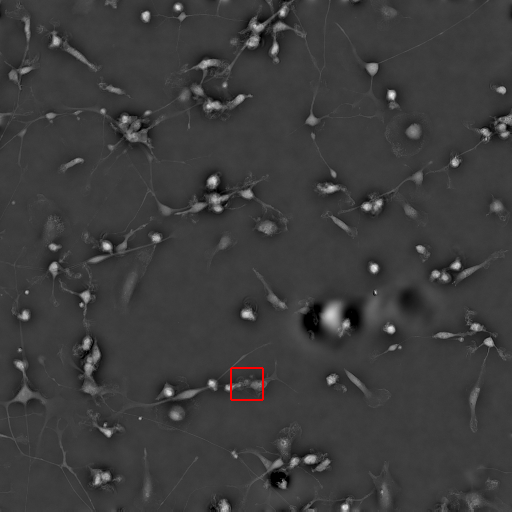

### supp_video_state_1_sample_bbox_Control-Site_5_29.gif

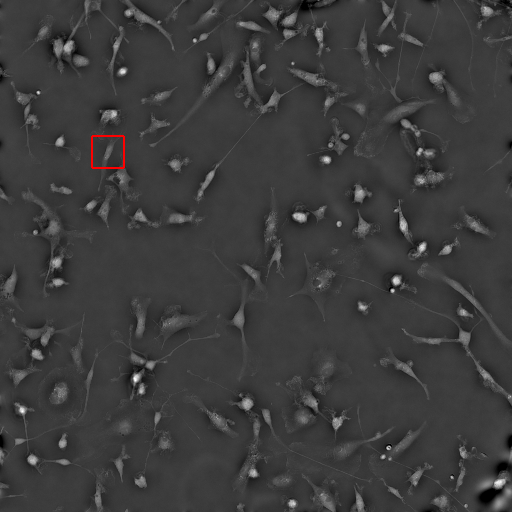

### supp_video_state_1_sample_bbox_GBM-Site_2_93.gif

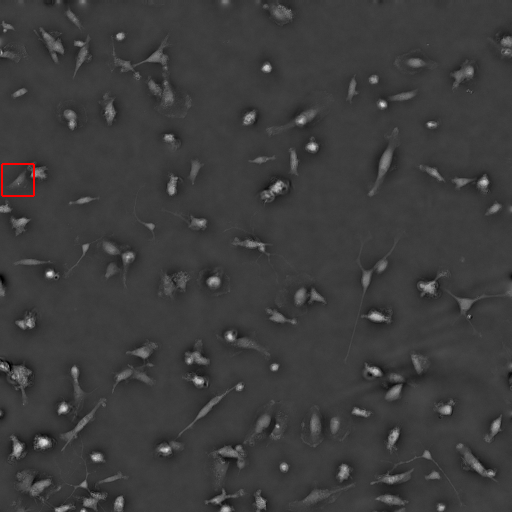

### supp_video_state_1_sample_bbox_IFNbeta-Site_1_288.gif

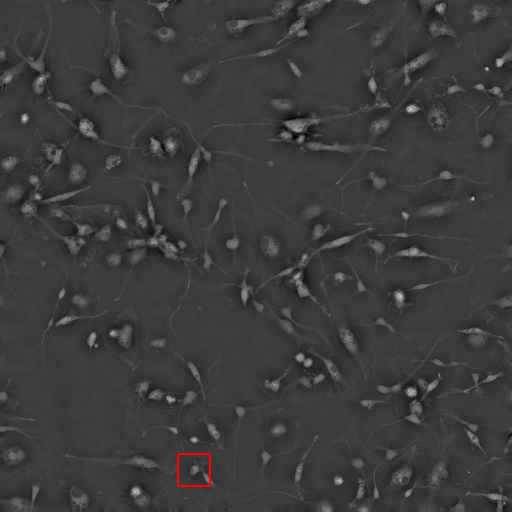

### supp_video_state_1_sample_bbox_IFNbeta-Site_7_67.gif

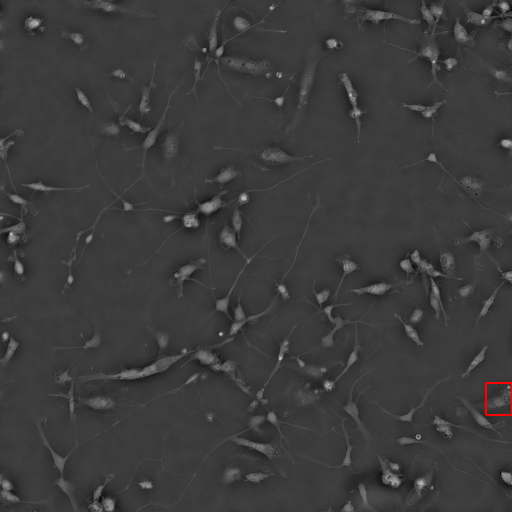

### supp_video_state_1_sample_movie_Control-Site_2_253.gif

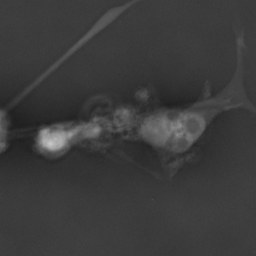

### supp_video_state_1_sample_movie_Control-Site_5_29.gif

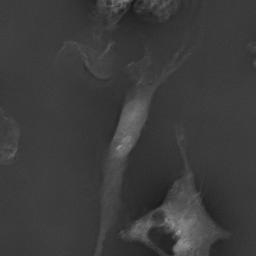

### supp_video_state_1_sample_movie_GBM-Site_2_93.gif

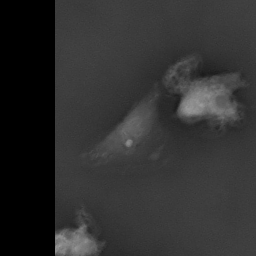

### supp_video_state_1_sample_movie_IFNbeta-Site_1_288.gif

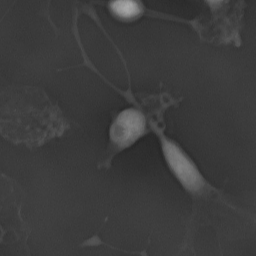

### supp_video_state_1_sample_movie_IFNbeta-Site_7_67.gif

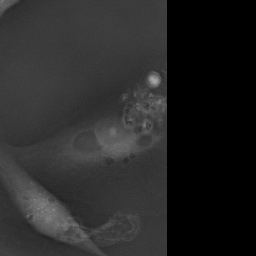

### supp_video_state_2_sample_bbox_Control-Site_2_154.gif

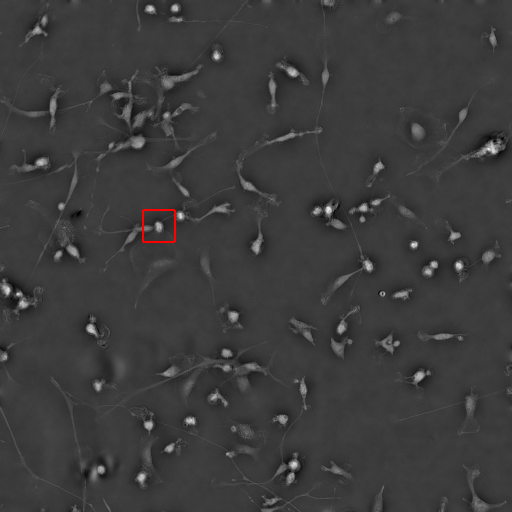

### supp_video_state_2_sample_bbox_Control-Site_2_299.gif

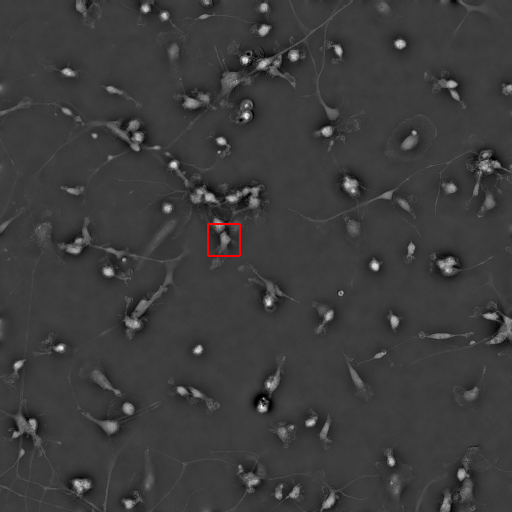

### supp_video_state_2_sample_bbox_GBM-Site_1_86.gif

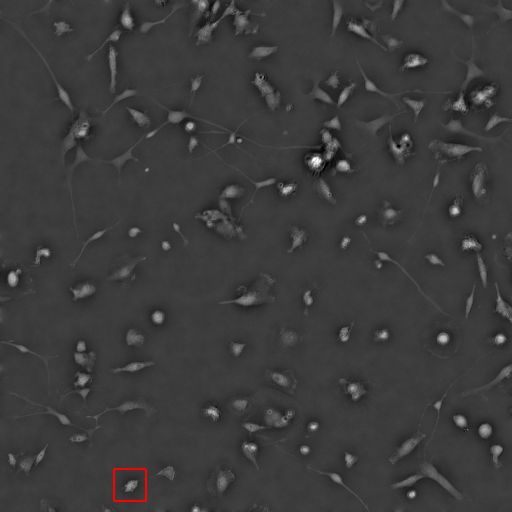

### supp_video_state_2_sample_bbox_GBM-Site_2_191.gif

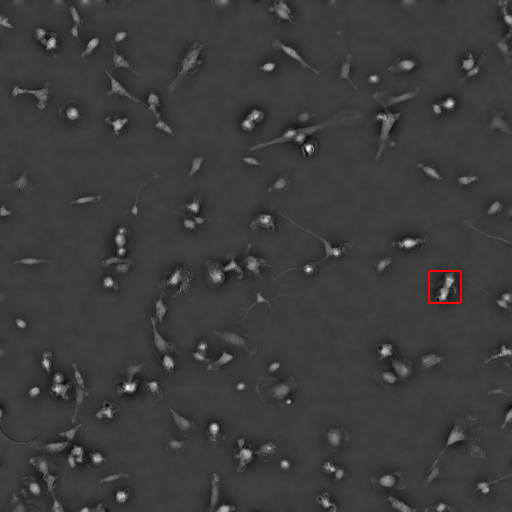

### supp_video_state_2_sample_bbox_GBM-Site_5_159.gif

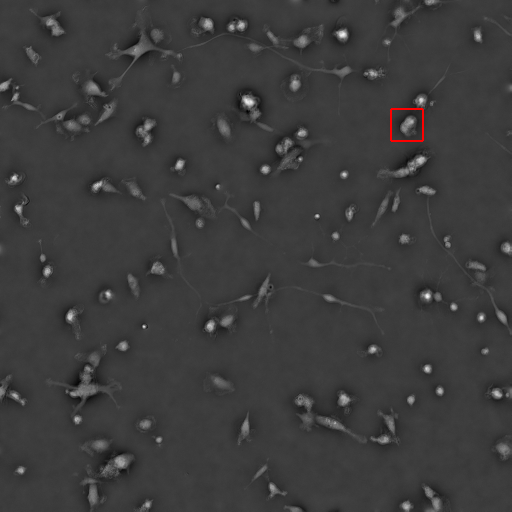

### supp_video_state_2_sample_movie_Control-Site_2_154.gif

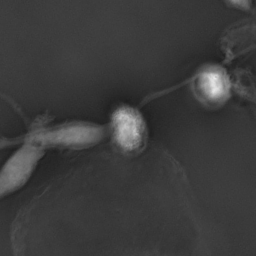

### supp_video_state_2_sample_movie_Control-Site_2_299.gif

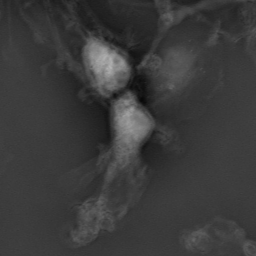

### supp_video_state_2_sample_movie_GBM-Site_1_86.gif

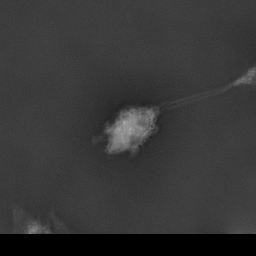

### supp_video_state_2_sample_movie_GBM-Site_2_191.gif

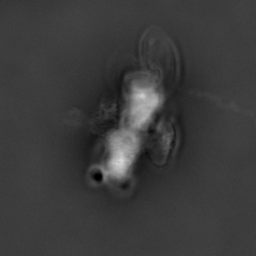

### supp_video_state_2_sample_movie_GBM-Site_5_159.gif

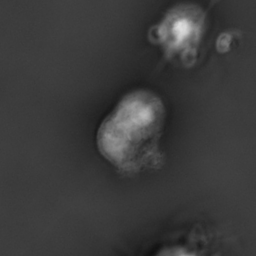

### supp_video_state_transition_bbox_Control-Site_1_5.gif

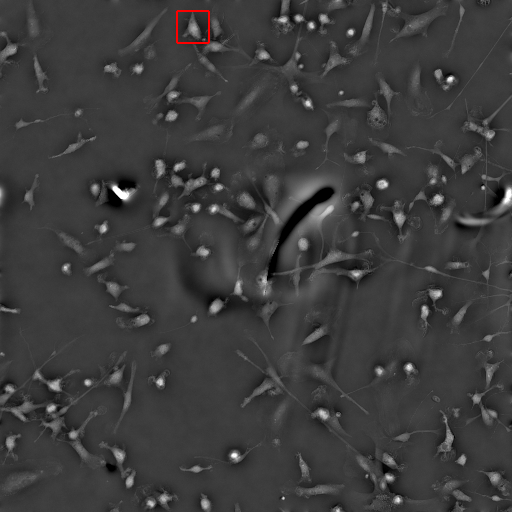

### supp_video_state_transition_bbox_Control-Site_3_22.gif

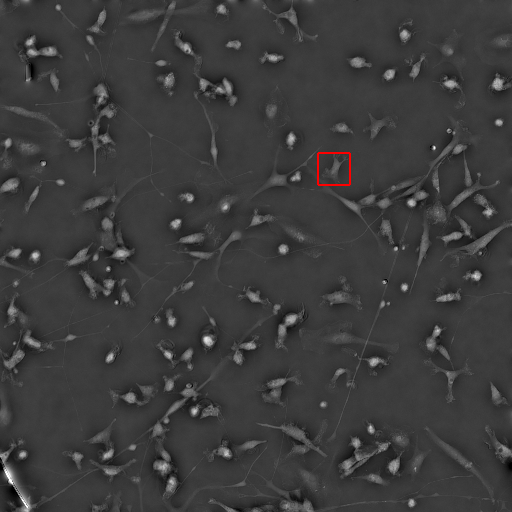

### supp_video_state_transition_bbox_Control-Site_5_74.gif

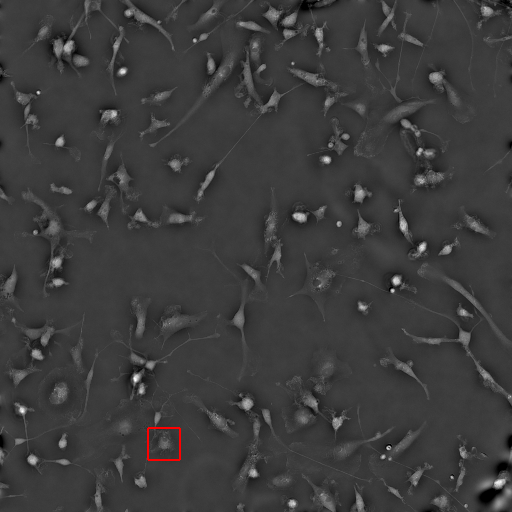

### supp_video_state_transition_bbox_Control-Site_5_189.gif

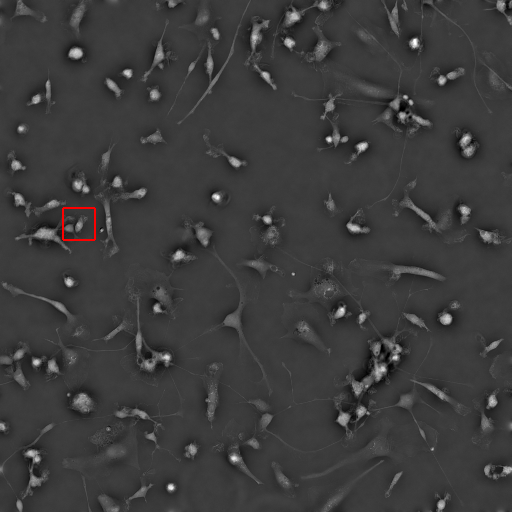

### supp_video_state_transition_bbox_Control-Site_5_329.gif

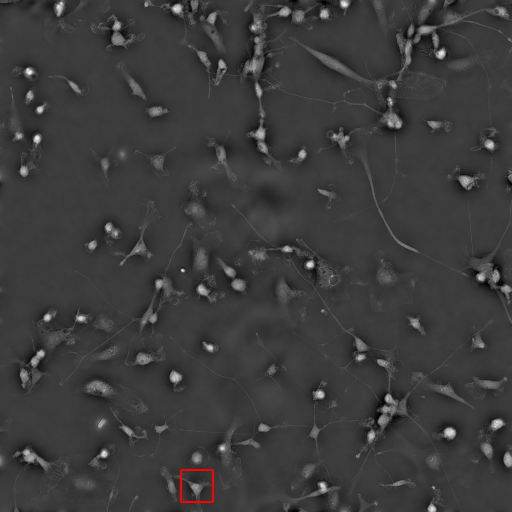

### supp_video_state_transition_bbox_GBM-Site_0_48.gif

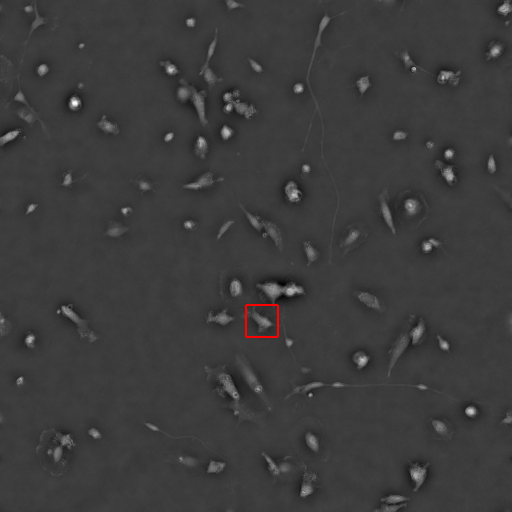

### supp_video_state_transition_bbox_GBM-Site_1_30.gif

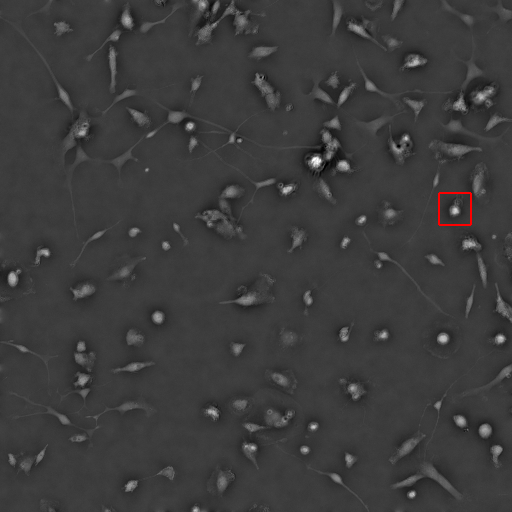

### supp_video_state_transition_bbox_GBM-Site_1_74.gif

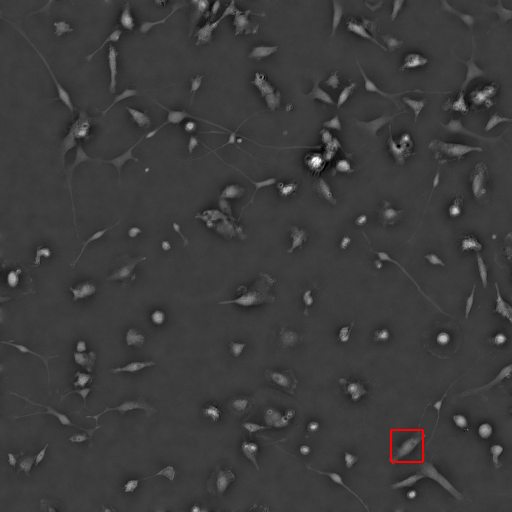

### supp_video_state_transition_bbox_GBM-Site_5_125.gif

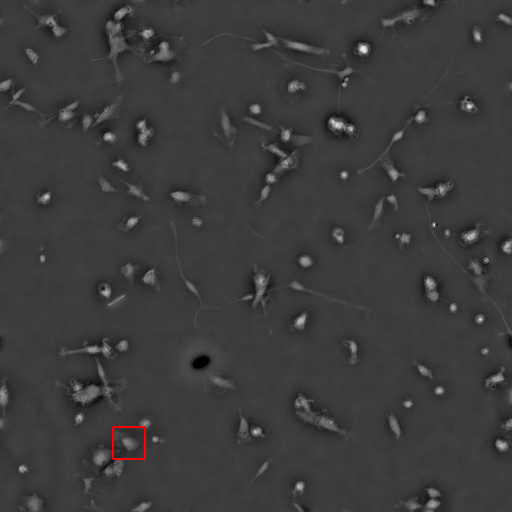

### supp_video_state_transition_bbox_GBM-Site_5_161.gif

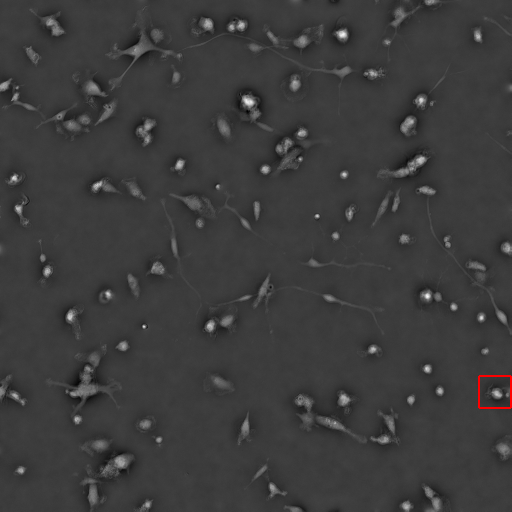

### supp_video_state_transition_movie_Control-Site_1_5.gif

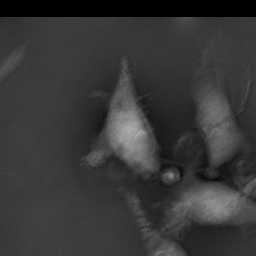

### supp_video_state_transition_movie_Control-Site_3_22.gif

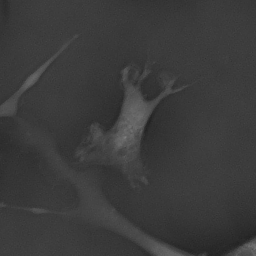

### supp_video_state_transition_movie_Control-Site_5_74.gif

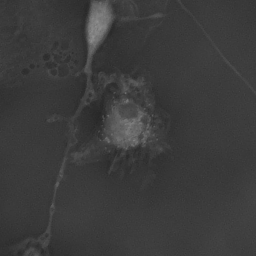

### supp_video_state_transition_movie_Control-Site_5_189.gif

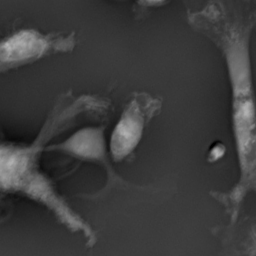

### supp_video_state_transition_movie_Control-Site_5_329.gif

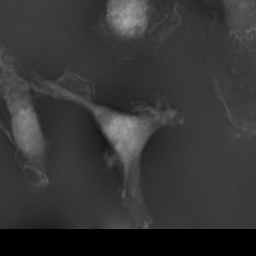

### supp_video_state_transition_movie_GBM-Site_0_48.gif

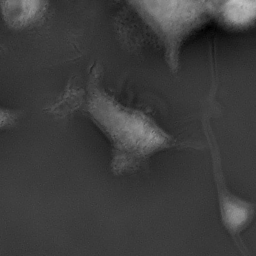

### supp_video_state_transition_movie_GBM-Site_1_30.gif

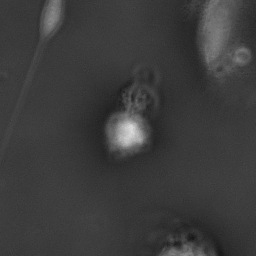

### supp_video_state_transition_movie_GBM-Site_1_74.gif

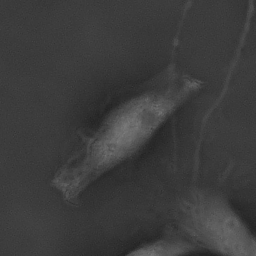

### supp_video_state_transition_movie_GBM-Site_5_125.gif

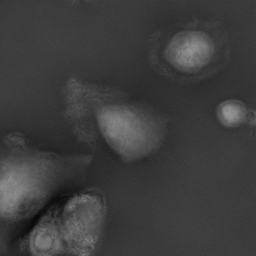

### supp_video_state_transition_movie_GBM-Site_5_161.gif

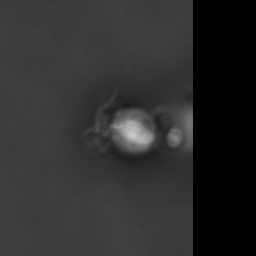
